# The localization of chitin synthase mediates the patterned deposition of chitin in developing Drosophila bristles

**DOI:** 10.1101/718841

**Authors:** Paul N. Adler

## Abstract

The insect exoskeleton is a complex structure that is a key for the life style of this very successful group of animals. It contains proteins, lipids and the N-acetyl glucosamine polymer chitin. Chitin is synthesized by the enzyme chitin synthase. In most body regions, chitin fibrils are found in a stack of parallel arrays that can be detected by transmission electron microscopy. Each array is rotated with respect to the layers above and below. In sensory bristles, chitin primarily accumulates in bands parallel to the proximal/distal axis of the bristle. These bands are visible by confocal microscopy providing experimental advantages. We have used this cell type and an edited chitin synthase gene to establish that the bands of chitin are closely associated with stripes of chitin synthase, arguing the localization of chitin synthase plays an important role in mediating the patterned deposition of chitin. This is reminiscent of what has been seen for chitin and chitin synthase in fungi and between cellulose and cellulose synthase in plants. Several genes are known to be essential for proper chitin deposition. We found one of these, *Rab11* is required for the insertion of chitin synthase into the plasma membrane and a second, *duskylike* is required for plasma membrane chitin synthase to localize properly into stripes. We also established that the actin cytoskeleton is required for the proper localization of chitin synthase and chitin in developing sensory bristles.

## Introduction

Chitin is an abundant and widespread extracellular polymer found in many types of eukaryotic organisms from fungi to vertebrates. It is synthesized by the multi-pass transmembrane enzyme Chitin Synthase (CS). This enzyme has principally been studied in fungi and insects, where chitin plays important structural roles. In fungi, chitin is a constituent of the cell wall and the number of CS genes is quite variable (Merzendorfer, 2011). For example, S. cereviase has 3 CS genes (Gohlke et al., 2017) while Aspergillius fumigatus has 8 (Muszkieta et al., 2014). Chitin in fungal cell walls is not uniformly distributed and in these systems, different CS’s appear to have different subcellular localizations and to mediate chitin synthesis in different parts of the cell wall including the bud ring in budding yeast (Cabib and Bowers, 1971; Foltman et al., 2018). In insects, chitin is a major component of the cuticular exoskeleton, the apical surface of trachea and the peritrophic membrane that lines the gut (Merzendorfer, 2011). In the cuticle, it is typically in parallel arrays while in the peritrophic membrane it is a fibrous mesh. There are two CS genes in insects, one functions in the formation of the cuticular exoskeleton and tracheal lining and the other synthesizes the chitin found in the peritrophic membrane (Merzendorfer, 2011). In Drosophila the *kkv* gene encodes the CS enzyme required for the synthesis of cuticle chitin (Moussian et al., 2005; Ostrowski et al., 2002).

The most conserved region in all chitin synthases is the catalytic domain (con1) (Dorfmueller et al., 2014; Nagahashi et al., 1995; Yabe et al., 1998) and this region is essential and sufficient for chitobiose synthesis by SC-CHS2. A second conserved region (con2) is essential for the synthesis of long chito-oligosaccharides, and seems likely to be essential for the translocation of growing chitin chains (Dorfmueller et al., 2014; Yabe et al., 1998). Both of these regions are thought to be cytoplasmic in yeast CHS2, although there is evidence for two transmembrane domains separating the catalytic site from at least the c terminal most part of Con2 (Gohlke et al., 2017). The number of inferred transmembrane domains varies from ∼5 to 18 with fungal CS proteins generally predicted to contain many fewer putative transmembrane domains than insect CS proteins (Gohlke et al., 2017; Merzendorfer, 2011) (Merzendorfer and Zimoch, 2003). In the case of fungal chitin synthases direct experimental data established that the computer programs for predicting transmembrane domains are useful but not able to predict accurately membrane protein topology (Gohlke et al., 2017). We report here experimental evidence that the amino terminus of Drosophila Kkv is in the cytoplasm and the carboxy terminus in the extracellular space.

The arrangement of chitin in insect cuticle may differ in different structures. Over most of the cuticle it is in layers of parallel arrays of chitin fibrils with each layer rotated with respect to its neighbors above and below (Bouligand, 1972; Moussian, 2013; Moussian et al., 2006a). As assayed by confocal microscopy, in the cuticle that covers the shaft of sensory bristles chitin is most abundant in bands that run parallel to the proximal-distal axis of the bristle (Nagaraj and Adler, 2012). In transmission electron micrographs, we did not see evidence for the presence of chitin layers in bristles or in hairs (trichomes). Whether this represents a true difference or is a consequence of a higher density of cuticle proteins masking the layers remains uncertain. In this paper, we make use of the bristle shaft as a model cell type to study patterned chitin deposition in insects. The large size of these polypoid cells makes them favorable for this purpose.

Chitin fibrils are insoluble at physiological pH (Elieh-Ali-Komi and Hamblin, 2016), which restricts models for patterned chitin deposition (Fig 1A). One possibility is that in insects as in fungi CS is localized in a patterned way to specific membrane domains and chitin deposition is directly patterned by this. In insect epidermal cells rows of elevated membrane called undulae have been proposed to be the site of chitin deposition (Moussian et al., 2007) (Moussian et al., 2006a). The tips of the undulae are associated with secreted extracellular matrix (Moussian et al., 2006a) (Adler, 2017) but it is not clear if this material is composed of chitin, cuticle proteins, other extracellular proteins/carbohydrate or more than one (or all) of these. Interestingly, prominent undulae are not seen during the deposition of some chitin containing cuticle, for example the cuticle that covers wing hairs or bristles (Adler, 2017; Sobala and Adler, 2016). Thus, it seems unlikely that undulae per se are essential for chitin or cuticle deposition. An alternative model is that the synthesis of chitin is not patterned but that chitin binding proteins bind to chitin fibrils as they are extruded through the membrane and serve as carriers to mediate the movement of the chitin to the correct place in the developing cuticle. There are a large number of proteins encoded by insect genomes that contain a chitin binding domain and could be part of such a system (Karouzou et al., 2007; Willis, 2010). In such a model, it seems likely that one or more unidentified proteins are first deposited in a patterned way and they interact with the chitin binding protein-chitin complex to guide the location for chitin fibril deposition. A third model is that CS containing exosomes/chitosomes are secreted and these are guided to the correct location for by interactions between exosome membrane proteins and one or more cuticle components. There are suggestions in support of this sort of model in the literature but evidence for exosomes has not been reported in transmission EM studies on cuticle deposition in Drosophila (e.g. (Sobala and Adler, 2016)).

**Figure 1.**
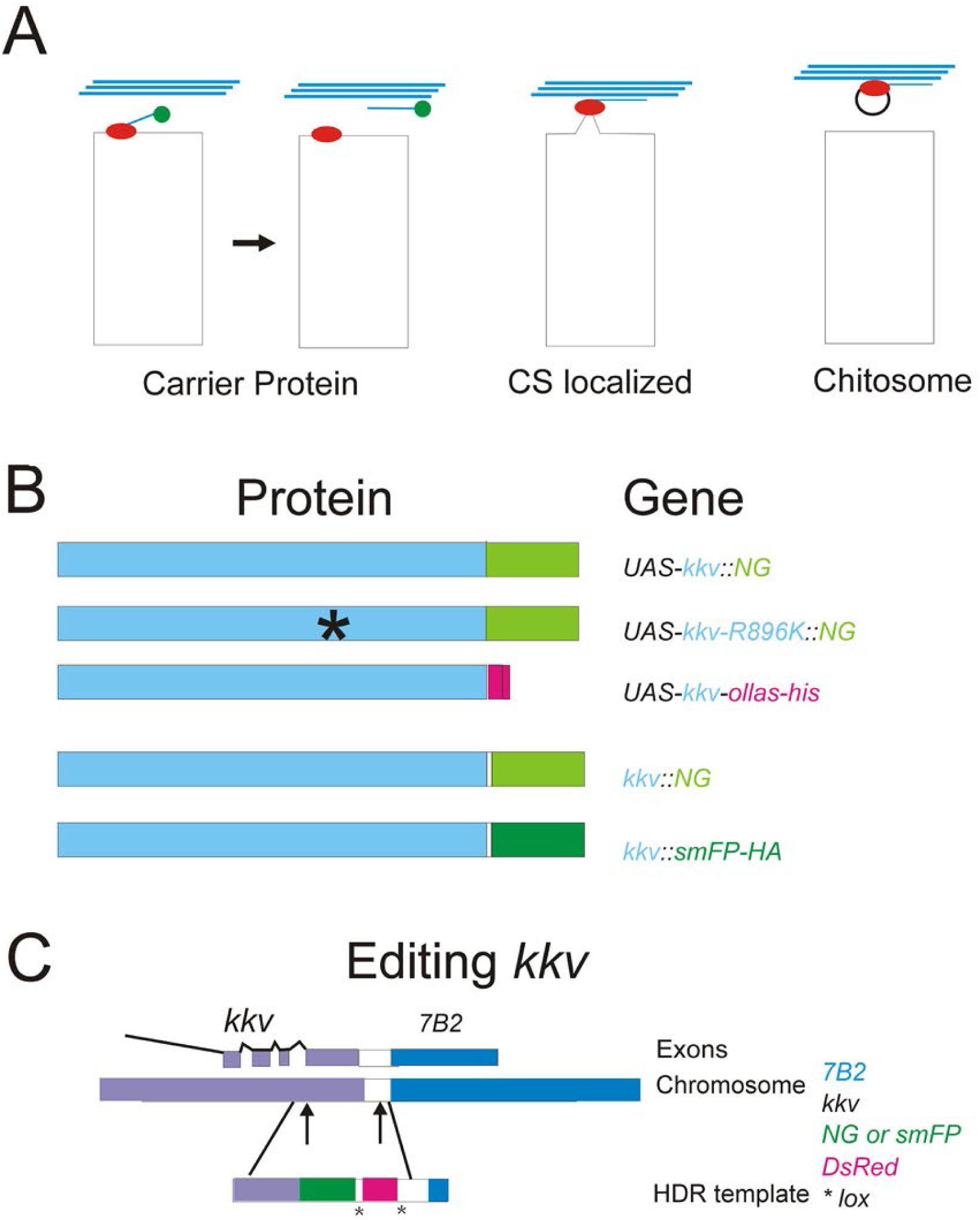
A. Models for chitin deposition and Chitin Synthase. B. Cartoons showing the proteins encoded by both UAS transgenes and edited genes. The asterisks indicates the R896K mutation. C. The approach used for the editing of *kkv*. The upward arrows indicate the target locations of the two guide RNAs. The upper part of the panel shows the 3’ end of *kkv.*

We used the accumulation of chitin bands in sensory bristle cuticle (Nagaraj and Adler, 2012) to examine the relationship between the localization of Chitin Synthase (Kkv) to the patterned accumulation of chitin. We find Kkv is closely associated with these chitin bands during cuticle deposition. Notably, this is true even when both patterns are highly abnormal. The accumulation of Kkv is not smooth like the chitin bands but is punctate. To clarify the text we use stripes to describe the accumulation of Kkv and bands to describe chitin. We previously identified several genes whose function was essential for the accumulation of bristle chitin in parallel bands (Nagaraj and Adler, 2012). Knocking down the function of two of these genes resulted in a failure of the accumulation of Kkv in stripes. These observations link the patterning of extracellular chitin to the patterning of Kkv localization in the apical plasma membrane of epithelial cells and begin the identification of genes that mediate both the insertion of Kkv into the plasma membrane and the patterned organization of Kkv in stripes along the proximal distal axis of the bristle.

## Results

### Generation and characterization of transgenes and edited genes that encode tagged Kkv

As a first step in examining the role of CS localization in the patterning of chitin deposition we generated a series of new genetic reagents consisting of 4 different *UAS-kkv* transgenes and two edits of the endogenous *kkv* gene (Fig 1BC) (see Methods for details).

In one of the UAS transgenes the *kkv* open reading frame was tagged on the C terminus by the bright mNeonGreen (NG) fluorescent protein (Shaner et al., 2013) (*UAS-kkv::NG*). In a second it was tagged by the ollas epitope tag (Park et al., 2008) and his_6_ (*UAS-kkv-OH*). Overexpression of these transgenes resulted in mild and limited gain of function phenotypes (see Supplementary Text File 1 and Fig S1). We also examined a variant of the NG tagged protein that contained the amino acid change found in the amorphic *kkv*-1 allele (R896K) (Moussian et al., 2005). In the fourth, multiple changes converted the catalytic domain to change convert it to the equivalent mosquito sequence.

We used CRISPR/Cas9 and Homology Dependent Repair (HDR) to edit the endogenous *kkv* gene to add two different C terminal tails (see Methods for details). In one we added the mNeonGreen fluorescent protein (NG) (Shaner et al., 2013) while in the second we added *smFP-HA* (Viswanathan et al., 2015) (Fig 1BC), which is a variant of super folder GFP with multiple HA tags inserted into loops of GFP. Both *kkv::NG* and *kkv::smFP-HA* were homozygous viable and showed no mutant phenotypes under a stereo microscope. Homozygous *kkv::NG* fly cuticle appeared normal when examined by compound light microscopy or scanning electron microscopy (Fig 2EF, S2CD – compare to EF). This was also true for *kkv::NG/DF* and *kkv::NG/kkv*^*1*^. The data establish the edited gene and protein are close to if not functionally equivalent to wild type. Homozygous *kkv::smFP-HA* flies displayed a phenotype of thin and bent hairs wing hairs (Figs 2EF, S2A). The phenotype appeared slightly stronger in *kkv::smFP-HA*/*Df* flies but we did not attempt to quantify this (Fig S2AB). In *ap-Gal4; UAS-kkv::NG kkv::smFP-HA/kkv::smGFP-HA* and *ap-Gal4; UAS-kkv::NG kkv::smFP-HA/Df(3R)ED5156* flies the wing hair phenotype was rescued in the dorsal wing cells where *ap* drives expression of *kkv::NG* but not in the ventral wing cells that served as an internal control (Fig 2GH). Similar results were obtained when *UAS-kkv-OH* was substituted for *UAS-kkv::NG.* We examined pupae that expressed Kkv::NG and Kkv::smFP-HA by in vivo confocal imaging. The level of fluorescence was much higher for Kkv::NG (we estimate it as being ∼15X brighter (see Methods and Fig S3)) and hence we used it for all of the protein localization experiments described below. These data are consistent with *kkv::smFP-HA* being a viable hypomorphic allele of *kkv* where a lower than normal level of protein accumulates. The hair phenotype is likely due to this structure requiring a higher level of chitin for normal morphogenesis. Consistent with this hypothesis wing trichomes are also the most sensitive cuticular structure to knocking down *kkv* expression using RNAi (pna, unpublished).

**Figure 2.**
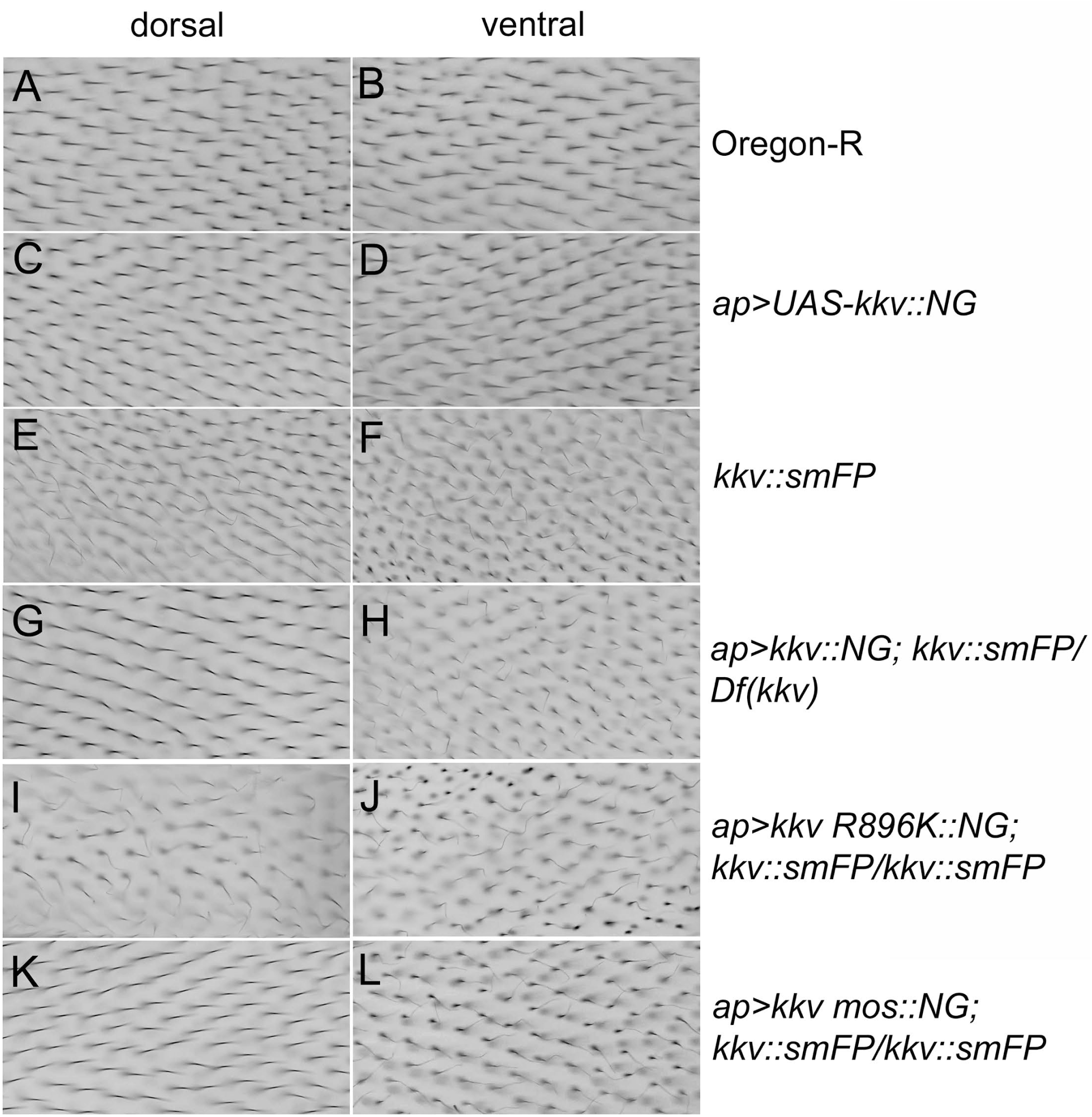
The rescue of the wing phenotype of *kkv::smFP* by the expression of a UAS transgene driven by *ap-Gal4.* A small region from the dorsal and ventral surface of the wing is shown for all genotypes. This region is from the posterior region. Note the thin bent hairs on both surfaces of the *kkv::smFP* homozygotes (EF) compared to those in wild type (AB) and *ap>kkv::NG* (CD) wings. Note the rescue in the dorsal surface of *ap>kkv::NG; kkv::smFP/Df* wings (GH). This is due to *ap* only driving expression of UAS transgenes in the dorsal surface cells. Note the failure to see rescue in *ap>kkv R896K::NG; kkv::smFP* wings (IJ) establishing that only the expression of a functional Kkv protein provides rescue. The rescue with *ap>kkv mos::NG* can be seen in the image of the dorsal surface of such wings (K) compared to the ventral surface (L).

### Subcellular Localization of Kkv

Kkv::NG localized to the apical surface of the large polytene salivary gland cells in *ptc>kkv::NG* larvae (Fig S4I). In *ptc>kkv::NG* wing discs we observed the expected stripe of expression (Fig S5A) in the middle of the wing. We saw no evidence for the secretion of Kkv::NG in these experiments. The stripe was obvious in living wing discs and in fixed wing discs stained with anti-NG antibodies, establishing the specificity of the antibodies. A similar specificity was observed for a rabbit polyclonal antibody (anti-Kkv-M) made against a region from the central part of the Kkv protein (aa1097-1246) and an anti-ollas monoclonal antibody. As expected (Maue et al., 2009; Moussian et al., 2015; Zimoch and Merzendorfer, 2002) Kkv::NG was preferentially localized apically in wing disc cells (Fig S5B, arrow). In *ap>kkv::NG* pupal wings Kkv::NG was localized external to the actin filaments found in the center of growing hairs (Fig 3G-L) (Adler et al., 2013; Turner and Adler, 1998; Wong and Adler, 1993). This close localization is reminiscent of apical F-actin and chitin in wing hairs (Adler et al., 2013) and in late stages of trachea development in Drosophila embryos (Ozturk-Colak et al., 2016).

**Figure 3.**
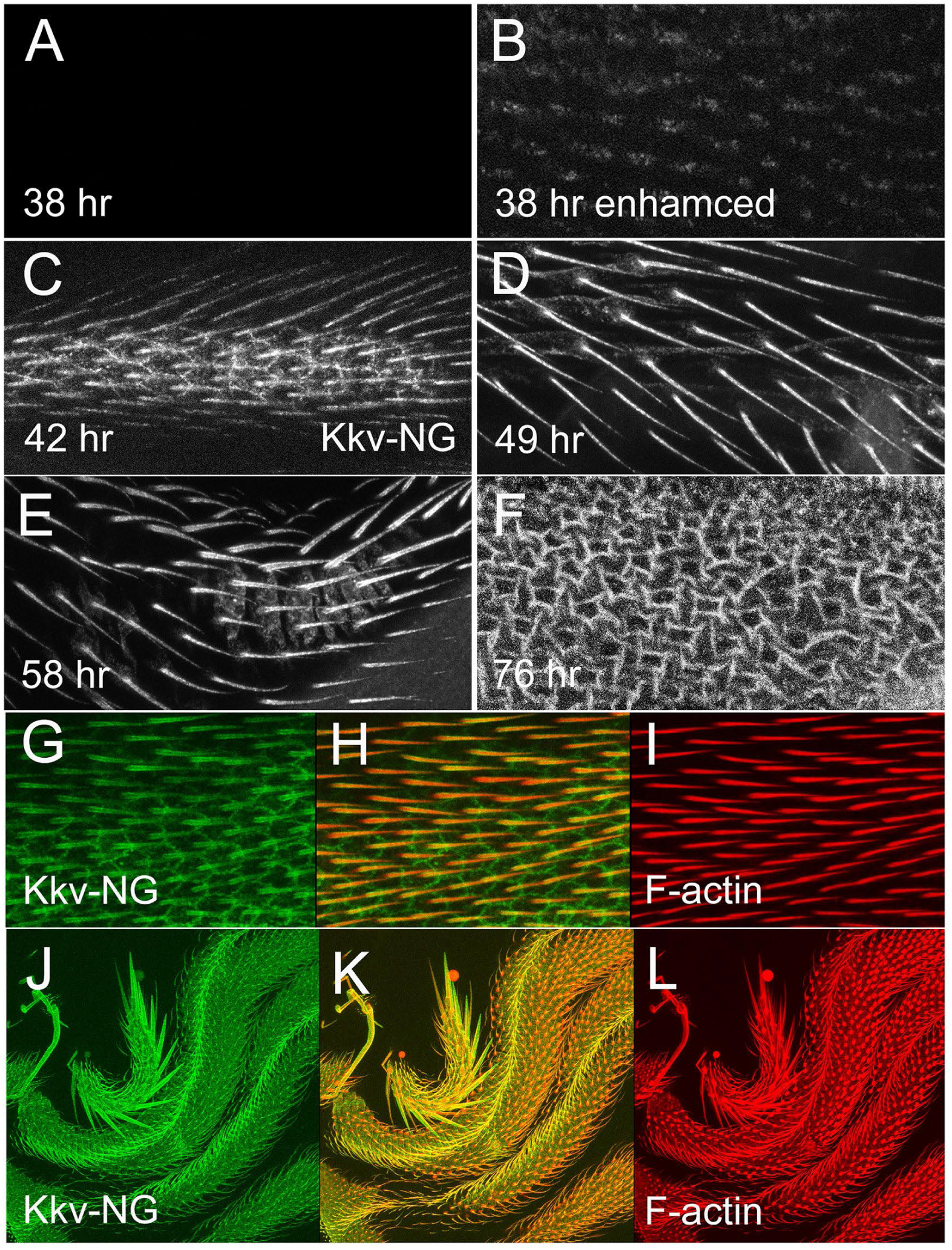
Localization of Kkv in the pupal wing. A-F. In vivo images of *kkv::NG* pupal wings. All except B are shown with the same microscope settings. Note the clear labeling of the hairs in wings from 42-58 hr. In the oldest wings the hairs are fainter and the image is from the focal plane where the pedestals are obvious. We did not detect hair labeling in the youngest wings (A) unless the image was enhance by brightening in ImageJ or Photoshop (B). G,H,I. Shown is a *ap-Gal4/+; UAS-kkv::NG* pupal 36 hr wing fixed and F-actin stained with phalloidin (red). Note the NeonGreen is external to the F-actin. One can also see that the NeonGreen signal is in the hair membrane and does not extend to the center of the hair. J,K,L. A fixed 48 hr *ap-Gal4/+; UAS-kkv::NG* wing stained for both NeonGreen (J-green) and F-actin (L-phalloidin - red). Note the bright disc of F-actin staining at the base of the hair is not associated with an accumulation of Kkv::NG.

The use of the Neon Green (and Ollas-His_6_) tag to localize Kkv requires that the tag remains associated with the enzyme. Since chitin synthases are often cleaved and this has been linked to enzyme activation (Broehan et al., 2007; Merzendorfer and Zimoch, 2003) (Zhang and Zhu, 2013) this is a concern (we have also seen evidence for cleavage of Kkv on Western blots - pna, preliminary results). To test if the tags remained associated with the enzyme we stained pupae with both a commercially available anti-NG monoclonal and our anti-Kkv-M rabbit polyclonal antibody. We observed a clear co-localization in stripes of puncta along the proximal distal axis in bristles (Fig 4 DEF). This established that the neon green tag from the fusion protein is an accurate reporter for the Kkv protein. We obtained similar results using anti-Ollas and anti-Kkv antibodies (Fig 4 GHI).

**Figure 4.**
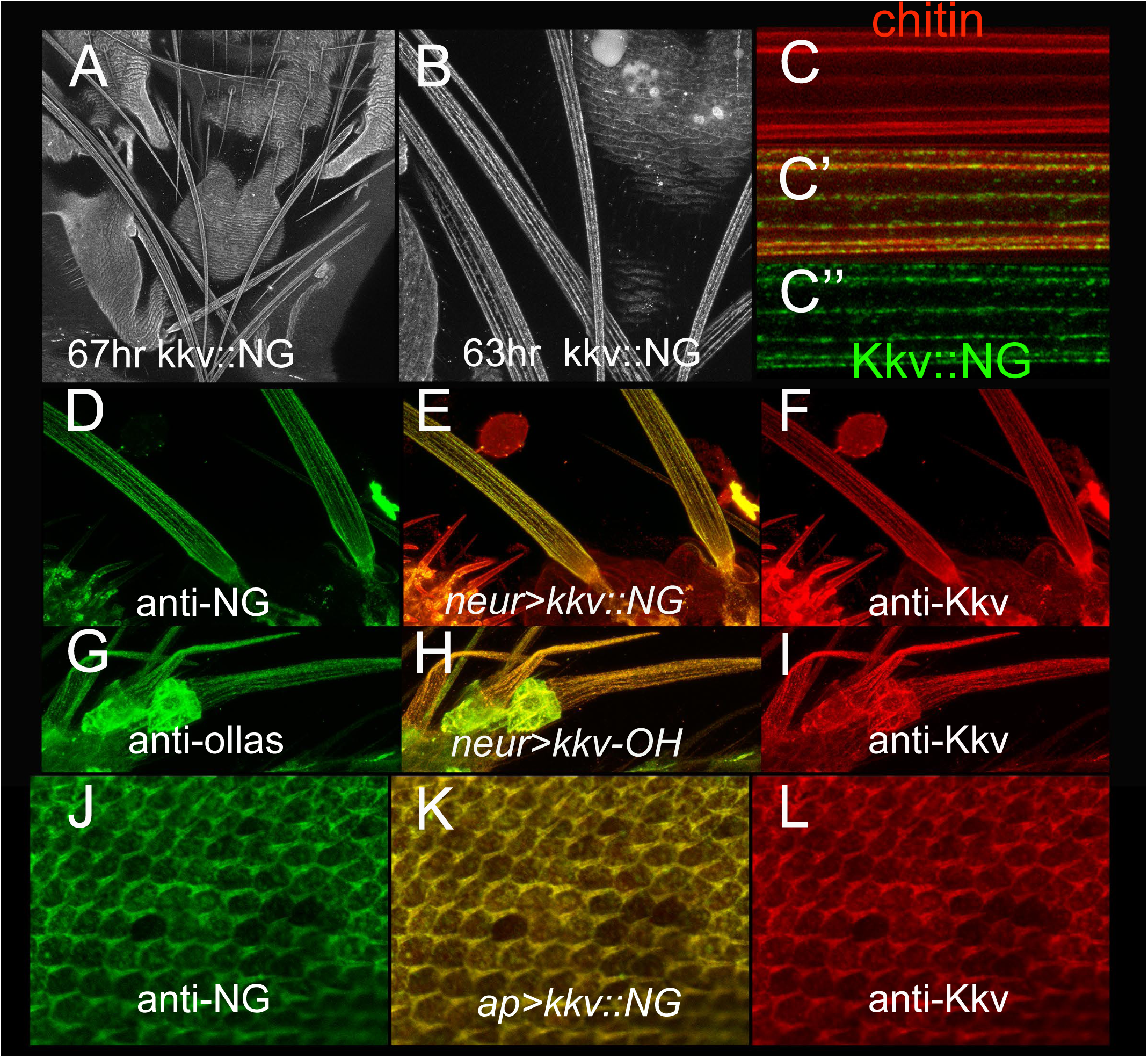
Localization of Kkv. A. A low magnification image of a *kkv::NG* notum. B. A higher magnification image of part of a *kkv::NG* notum. Note the stripes of Kkv::NG along the proximal distal axis of the bristles. C (C’ and C”). A high magnification image of a bristle from a *UAS-ChtVis/+; neur-Gal4/kkv::NG/+* bristle. Note the relatively smooth bands of chitin (red) and the punctate stripes of Kkv::NG (green) and the association between the two. D,E,F. Two bristles from a *neur-Gal4/kkv::NG* pupae immunostained for NeonGreen (green-D) and Kkv (red-F). E is the merged image and shows the high degree of co-immunostaining. G,H,I. Bristles from a *neur-Gal4/kkv-OH* pupae immunostained for ollas (green-G) and Kkv (red-I). H is the merged image and shows the high degree of co-immunostaining. J,K,L. Pupal wings from *ap-Gal4/+; UAS-kkv::NG* immunostained for NeonGreen (green-J) and Kkv (red-L). K is the merged imaged and shows a high degree of co-localization.

### Localization of Kkv encoded by the edited endogenous gene

We first examined the accumulation of Kkv::NG in pupal wings as this tissue is best characterized for the timing of cuticle deposition and gene expression (Adler et al., 2013; Sobala and Adler, 2016). Previously we found that we could first detect chitin in developing hairs wing hairs around 42 hr after white prepupae (awp) (Adler et al., 2013). We observed Kkv::NG in developing hairs in living pupal wings at 42, 49 and 58 hrs awp (after white pupae) (Fig 3C). The level of fluorescence was lower in 42 hr hairs than in 49 hr hairs. We could also detect Kkv::NG fluorescence in the apical surface of wing cells at these times with higher levels at cell boundaries. This was a bit surprising since the procuticle deposition starts later (∼ 56-58 hr awp) in the wing blade (Adler et al., 2013; Sobala and Adler, 2016). In digitally enhanced images we also observed evidence for Kkv::NG in 38 hr awp wings in the proximal part of the hair (Fig 3B). In older wings (e.g. 76 hr) during the middle of procuticle deposition the apical membrane fluorescence was stronger than at earlier times and we could see the pedestals that the hairs are found on at late stages (Fig 3F) (Mitchell et al., 1990; Sobala and Adler, 2016).

We next examined Kkv::NG in thoracic bristles by in vivo imaging from ∼40-80 hr awp. Previously we found that chitin accumulated in bands along the proximal distal axis of thoracic bristles starting around 42 hr awp (Nagaraj and Adler, 2012). The level of Kkv::NG fluorescence in younger than 50hr awp bristles was lower than in older bristles. (Fig S6). At both early and later stages, Kkv::NG fluorescence had a punctate appearance within an overall pattern of stripes along the proximal distal axis of the bristle (Figs 4A-C, 5J, S5E, S6). As development proceeded, the stripes became more complete although they never reached the completeness and smoothness seen with chitin bands. In older animals (>70 hrs) the pattern became somewhat less distinct with more inter-stripe fluorescence (Fig S6). When we examined orthogonal views of bristle image stacks the stripes of Kkv were obvious (Fig 6A). Unless stated otherwise we examined bristles from 50-65 hr old animals as these showed the most dramatic “stripe pattern”. Control experiments with Oregon-R pupae established that the florescence we were observing was due to the edited *kkv* gene and not to autofluorescence (Fig S5C-F). A similar, albeit perhaps a bit less precise stripe pattern was seen in developing *neur-Gal4*, UAS-*Kkv::NG* bristles.

**Figure 5.**
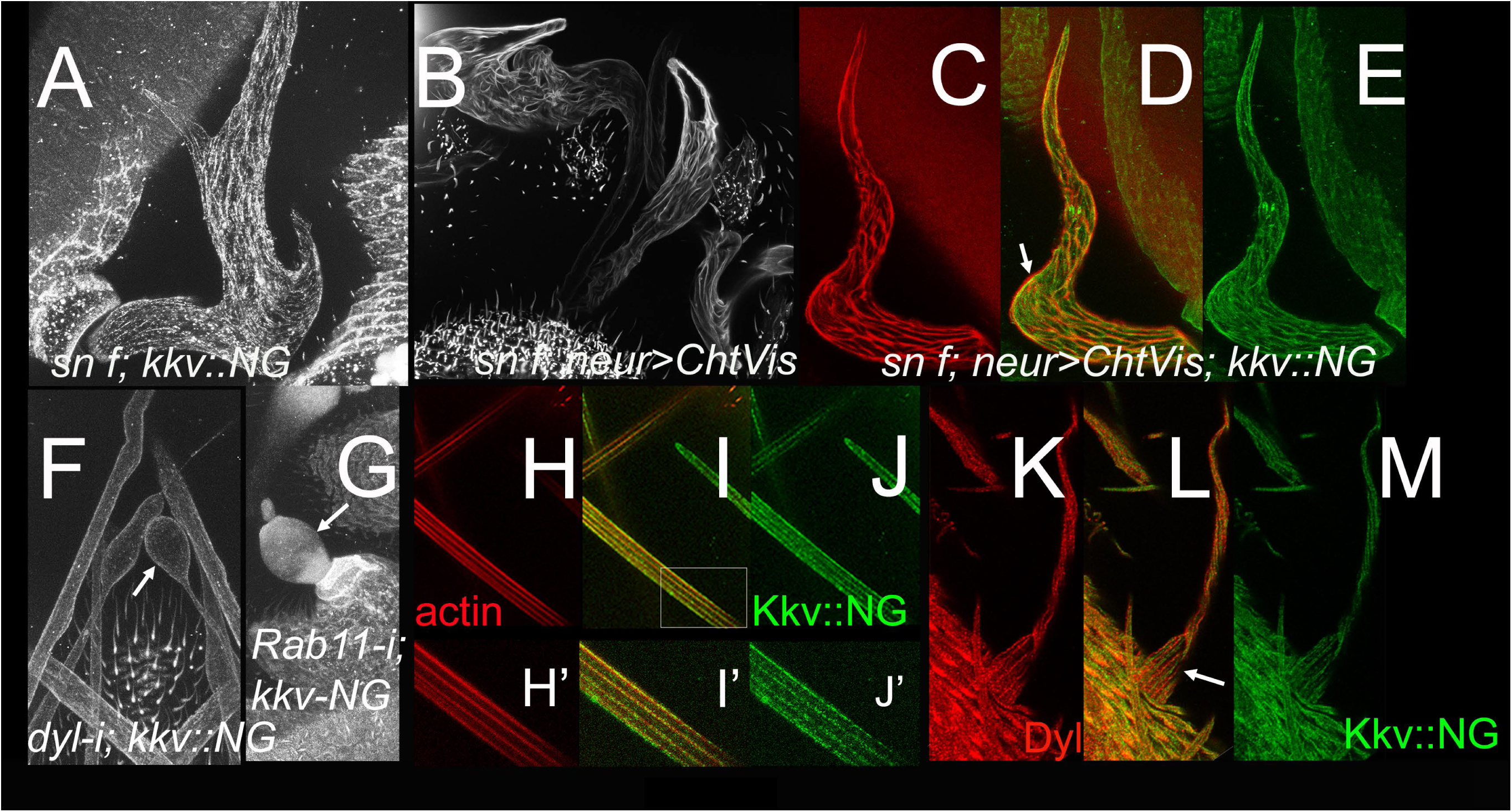
Factors that mediate the localization of Kkv in stripes in bristles. A. A *sn f; kkv::NG* bristle by in vivo imaging. The lack of the large F-actin bundles due to the loss of *sn* and *f* leads to highly abnormal shape and abnormal pattering of Kkv::NG stripes. B. A *sn f;UAS-ChtVis/+; neur-Gal4/+* bristle by in vivo imaging. The lack of the large F-actin bundles due to the loss of *sn* and *f* leads to highly abnormal bristle shape and to abnormal patterning of chitin. C,D,E. A *sn f; UAS-ChtVis/+; neur-Gal4 kkv::NG/kkv::NG* bristle by in vivo imaging. The lack of the large F-actin bundles due to the loss of *sn* and *f* leads to highly abnormal shape and abnormal patterning of both chitin (C - red) and Kkv::NG (E – green) stripes. Note the close association of Kkv and chitin (D- merged image, arrow) even though the pattern as a whole is highly abnormal. F. A *UAS-dyl RNAi; neur-Gal4 kkv::NG/kkv::NG* thorax by in vivo imaging. Note the bulged bristle (arrow) and the lack/great reduction of Kkv stripes. G A *UAS-Rab11-RNAi; neur-Gal4 kkv::NG/kkv::NG* bristle by in vivo imaging. Note the stub bristle phenotype (arrow) and the lack of Kkv::NG stripes. H,I,J. A UAS-Ruby-Lifeact; *neur-Gal4 kkv::NG/kkv::NG* bristle showing the large bundles of F-actin (H – red) and stripes of Kkv::NG (J – green). H’, I’,L’ are higher magnification images where the overlap between the F-actin bundles and Kkv::NG stripes is obvious. KLM. *Kkv::NG* bristles immunostained with anti-Dyl antibody (K – red) and anti-NeonGreen antibody (M – green). In the merged image (L) the interdigitated stripes of red and green can be seen (arrow).

**Figure 6.**
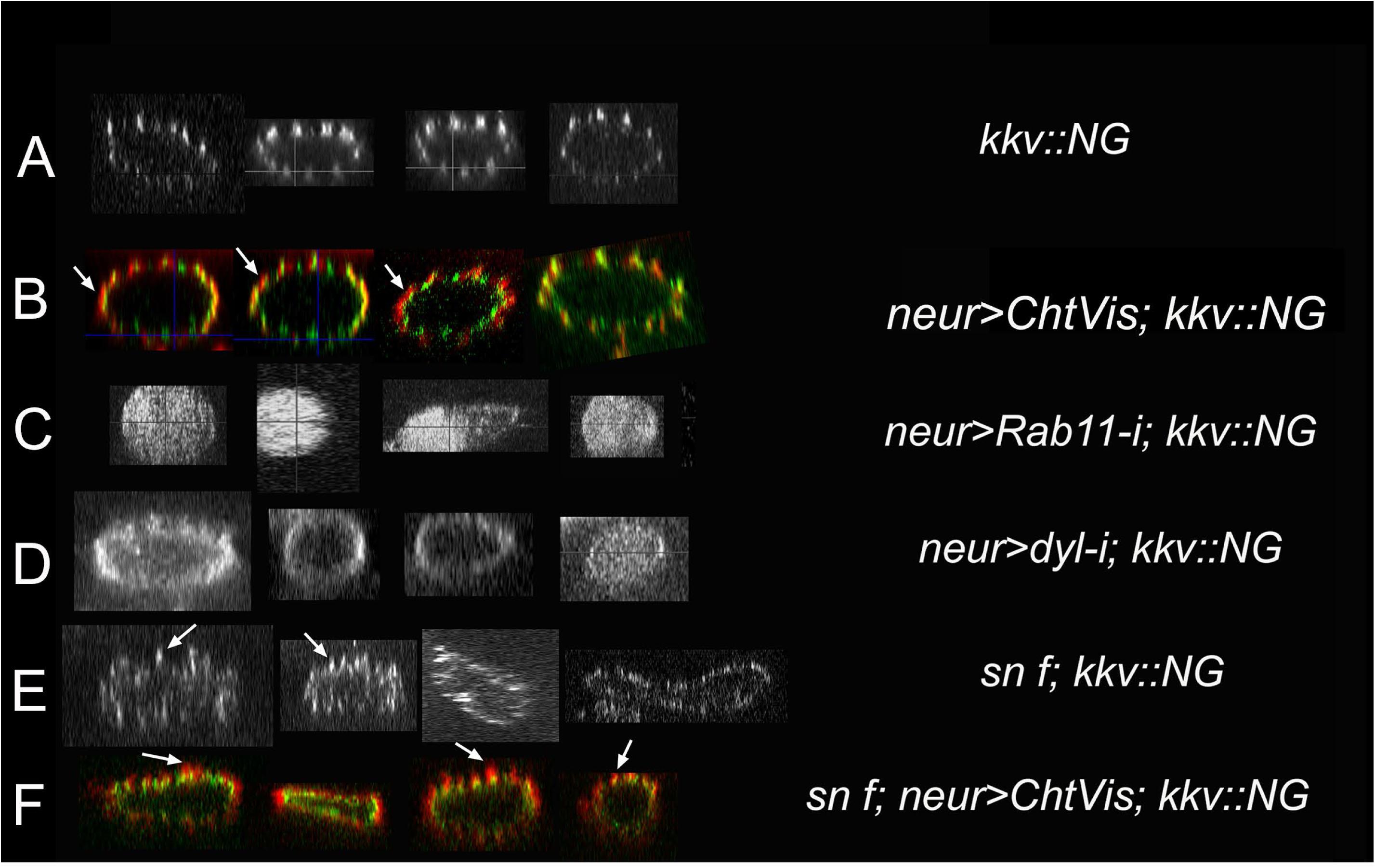
Orthogonal cross sections of bristles of various genotypes. A. *kkv::NG* shows a distinct pattern of stripes. B. *UAS-ChtVis/+; neur-Gal4 kkv::NG/kkv::NG* shows the close association of the chitin bands and the stripes of Kkv::NG. In many cases the chitin is external to Kkv::NG (arrows). C. *UAS-Rab11-RNAi; neur-Gal4 kkv::NG/kkv::NG* bristles show a failure in the localization of Kkv to the plasma membrane. D. *UAS-dyl RNAi; neur-Gal4 kkv::NG/kkv::NG* bristles show much of the Kkv::NG is inserted into the plasma membrane but it is not localized into the strip pattern (e.g. A). E. *sn f; kkv::NG* bristles have an abnormal cross section and evidence for abnormal stripes of Kkv::NG are evident (arrows). F. *sn f; UAS-ChtVis; neur-Gal4 kkv::NG/kkv::NG* bristles show the expected abnormal cross section shape with the irregular banding of Kkv::NG and chitin. Note the close association of Kkv::NG and chitin (arrows) even in these very abnormal bristles.

To investigate the localization of chitin and CS we carried out experiments where we localized both Kkv::NG and our chitin reporter (Cht-Vis) in bristles in living pupae (Sobala et al., 2015). The bands of ChtVis differed from those of Kkv::NG by being smooth rather than punctate (Fig 4C-C”). However, the two patterns were largely co-aligned in stripes. In cross section the ChtVis signal was usually exterior to the Kkv-NG signal as expected for chitin being secreted and CS being in the plama membrane (Fig 6B, arrows). The stripe of Kkv::NG was often also offset a bit from the ChtVis signal, which could be a consequence of ChtVis reporting on chitin (an accumulated product) while the Kkv::NG signal represent protein at a particular instance in time. The different cellular location (plasma membrane vs extracellular) and geometry likely contributes to this (Fig S7).

### The actin cytoskeleton influences the accumulation of Kkv

We also explored the relationship between Kkv::NG accumulation and the large bundles of cross-linked F-actin found in bristles (Tilney et al., 1995). These experiments were complicated by the breakdown of the actin bundles, which starts around 43 hr (Guild et al., 2002), and our inability to reliably immunostain bristles older than about 48 hr awp. In our best experiments we examined pupae that were 48hr or younger. An additional complication is geometric and due to Kkv being in the plasma membrane (and the NG in Kkv::NG being extracellular as described below) while the F-actin extends some distance into the cytoplasm (Tilney et al., 1995) (Fig S7). Using an F-actin reporter (Lifeact-Ruby - (Hatan et al., 2011; Riedl et al., 2008) we observed a close connection between the localization of Kkv::NG and the large bundles of F-actin in *neur>lifeact-Ruby; kkv::NG* pupae. The results varied from the two appearing to co-localize to their being slightly offset (Fig 5H-J, H’-J’) consistent with the geometry considerations (Fig S7).

The large bundles of highly cross-linked actin filaments support the shape and growth of developing bristles and in their absence in *sn*^*3*^ *f*^*36*^ double mutants the resulting bristles are bent, curved, split, shorter and stand more upright than normal (Guild et al., 2002; Tilney et al., 2004; Tilney et al., 1995). In separate experiments, we observed an abnormal distribution of chitin and Kkv::NG in living *sn*^*3*^ *f*^*36*^ double mutants. The robust parallel array of chitin bands and Kkv::NG stripes were severely disrupted (Fig 5AB). Kkv::NG was primarily in the plasma membrane and the chitin appeared to be extracellular so the bundles of F-actin do not appear to be required for the localization of either. To determine the relationship between the abnormal stripes of Kkv::NG and chitin we examined the distribution of both in the same living bristle. We found substantial co-localization of chitin and Kkv::NG conserved in these highly abnormal bristles (Fig 5CDE).

### Proteins required for the proper localization of Kkv

We previously established that Rab11 and exocyst function is required for the deposition of cuticle and the bands of chitin in bristles (Nagaraj and Adler, 2012). Affected bristles become unstable and collapse after the highly cross-linked F-actin bundles in developing bristles begin to depolymerize (Guild et al., 2002; Nagaraj and Adler, 2012). To determine if this was associated with improperly localized CS we examined Kkv::NG in thoracic bristles of living *neur-Gal4 Gal80-ts/Rab11 RNAi; kkv-NG/kkv-NG* pupae shifted to 29.5°C at wpp (white prepupae). These animals developed the extreme stub macrocheatae phenotype (Fig 5G, arrow) described previously (Nagaraj and Adler, 2012). The morphology of the developing bristles depended on pupal age. In the youngest animals examined, the bristles showed the blebbing characteristic of the early stages of the collapse program (Nagaraj and Adler, 2012). In older animals the collapsed stub bristle morphology was seen (Fig 5G, arrow). The stripes of Kkv::NG were lost and in Z sections we found that the protein was cytoplasmic (Fig 6C). Thus, Rab11 function is required for the insertion of Kkv into the plasma membrane and hence the loss of chitin bands in the mutant (Nagaraj and Adler, 2012).

The Zona Pellucida domain Dusky-Like (Dyl) protein acts as a Rab-11 effector for chitin deposition in bristles (Nagaraj and Adler, 2012). We examined living *UAS-dyl-RNAi/+; neur-Gal4 kkv::NG/kkv::NG* pupae and observed the bristle blebbing phenotype was associated with the loss of the robust striping pattern of Kkv::NG accumulation (Fig 5F, arrow). Kkv::NG in such bristles was primarily spread around the plasma membrane (Fig 6D). The distribution was not uniform but was far from the nicely spaced stripes seen in wild type. The data indicate that Dyl is required for the localization of Kkv::NG in stripes but not for its insertion into the plasma membrane. The difference in Kkv::NG localization between the Rab11 and Dyl knockdowns suggests that these two genes and proteins mediate different steps in the localization of Kkv.

We also simultaneously localized Kkv-NG and Dyl in bristles by immunostaining. The stripes of Kkv::NG and Dyl were interdigitated but did not appear to touch (Fig 5KLM, arrow). We further established that the large actin bundles are essential for the accumulation of Dyl in stripes in bristles (Fig S8AB).

### Localization of an inactive mutant Kkv

We next addressed whether the catalytic activity of Kkv might impact its subcellular localization by placing a R896K mutation into *UAS-kkv::NG*. This missense mutation is the cause of the amorphic *kkv*^1^ allele and is in an invariant site that is part of the enzyme’s active site (Dorfmueller et al., 2014; Merzendorfer, 2006; Moussian et al., 2005; Nagahashi et al., 1995). We first tested if this transgene could rescue the wing hair phenotype seen in *kkv::smFP* flies. The wings of *ap-Gal4/+; UAS-kkv-R896K::NG kkv::smFP/kkv::smFP* flies showed no evidence of rescue of the *kkv::smFP* hair phenotype (Fig 2IJ). The failure of this transgene, which encodes a protein that very likely has little or no catalytic activity, provides support for the validity of the *kkv::smFP* rescue assay.

We observed a failure of the normal apical localization of Kkv-R896K::NG in salivary gland cells (compare Fig S4I and J). At higher magnification, we observed Kkv::NG accumulated in vesicles that contained bright puncta (Fig S4I’, arrow). In contrast the mutant protein accumulated either between vesicles or in abnormally shaped vesicles (Fig S4 J’). Kkv-R896K::NG also failed to accumulate in pupal wing hairs (Fig S4A-D) and it did not accumulate in stripes in bristles (Fig S4E-H). As a control we immunostained pupal wings and bristles that expressed Kkv-R896K::NG using both the anti-NG and anti-Kkv-M antibodies, and observed extensive co-localization (Fig S9S-F). Hence, the mislocalization was not due to cleavage of the NG reporter from the mutant protein.

### A Kkv with a catalytic domain mutated to a mosquito catalytic domain is functional

We next attempted a more ambitious test of the *kkv-smFP* rescue assay using a UAS transgene where the con1 domain of *kkv* was replaced by the equivalent region of a mosquito chitin synthase (*UAS-kkv-mos::NG* – see Fig 1B and Methods for details). We generated *ap-GaL4/+; UAS-kkv-mos::NG kkv::smFP/kkv::smFP* flies and found that the mutant wing hair phenotype of *kkv::smFP* to be fully rescued in the dorsal but not ventral wing surface hairs (Fig 2KL) indicating that this “hybrid” protein is active.

### Topology of Kkv

Reagents we generated for other reasons provided us with tools we could use to probe the topology of Kkv. Using a collection of programs to predict the location of transmembrane domains led us to a consensus of 14 predicted transmembrane domains (Fig 7B and Methods). We found both anti-NG and anti-Ollas antibodies stained wing discs expressing the relevant transgene in the absence of permeabilization indicating that the C terminus is exposed to the extracellular space (Fig 7A). All of the topology programs predicted this. In contrast when we used an anti-Kkv polyclonal antibody (anti-Kkv-raised against aa 1097-1246 we did not see any staining in the absence of Triton X-100 permeablization (Fig 7A), arguing this region is intracellular. This disagrees with the consensus prediction. Similar results were obtained when we used antibodies directed against aa 53-66 and also antibodies directed against aa 530-541 – a region located not far from the catalytic domain (Fig 7AB). It is worth noting that no programs predicted a transmembrane domain in the region encompassing aa 1-53, which when combined with our data suggests that the amino terminus of Kkv is located in the cytoplasm.

**Fig 7.**
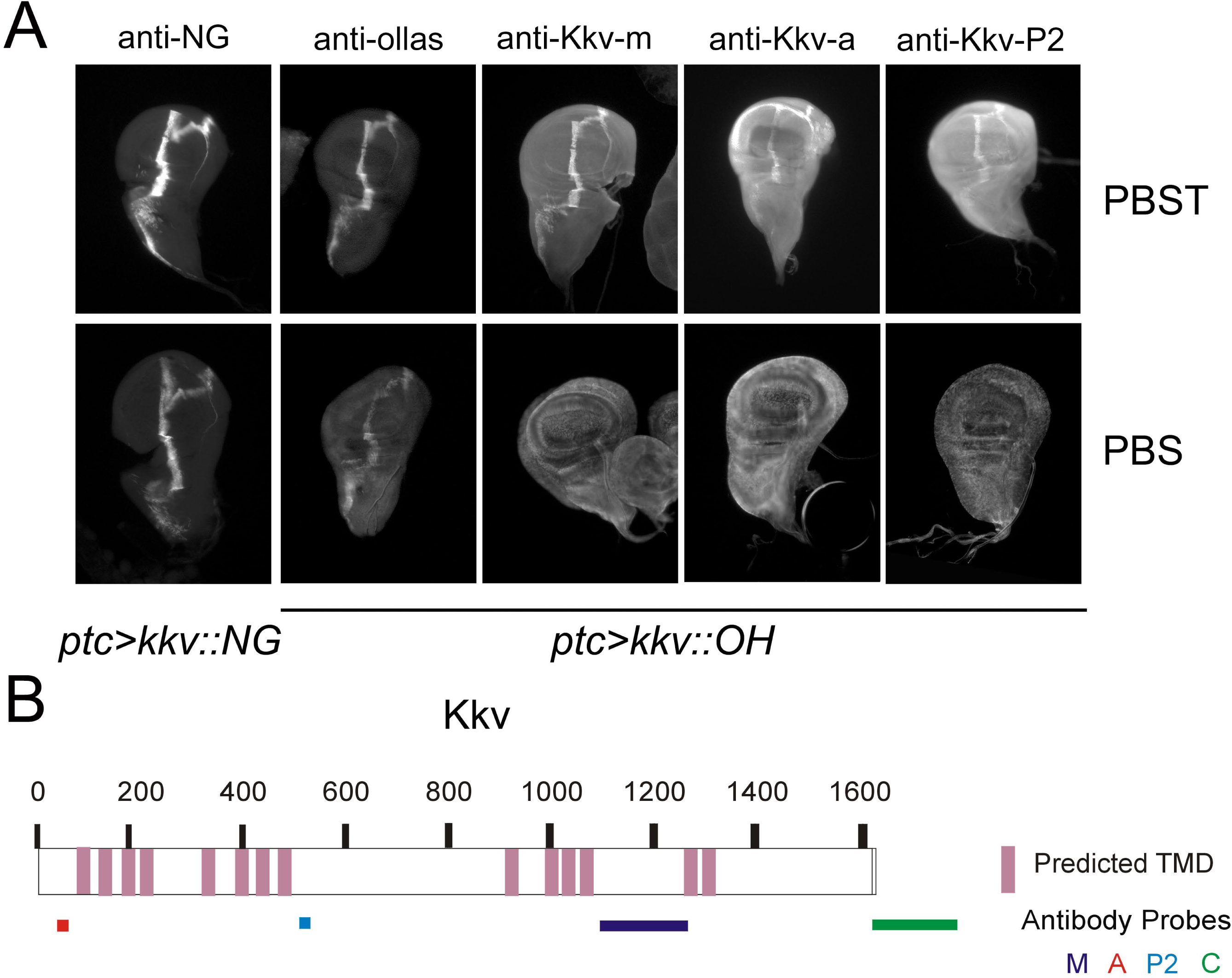
Topology of Kkv. A. *ptc-Gal4 UAS-kkv::NG* (or *UAS-kkv-*OH) wing discs where *kkv* is expressed in a stripe immunostained with various antibodies with or without detergent treatment (PBST vs PBS). B. A cartoon showing the consensus transmembrane domains as described in the Methods. The location of the epitope recognized by each of the antibodies. The data indicate that the amino terminus is cytoplasmic and the carboxy terminus is extracellular.

## Discussion

### The patterning of Chitin deposition in bristles is linked to the localization of Kkv

Our observations establish that the localization of chitin deposition is closely linked to the localization of Kkv (Chitin synthase) in bristles. The most compelling data being that chitin and Kkv remain largely colocalized even when the distribution of both is highly abnormal. A limitation of our observations is that they focused on the bands of chitin seen in developing sensory bristles. The cuticle that covers the sensory bristles differs in two ways from the cuticle that covers much of the fly’s body. First, we have not detected the layering of chitin in TEM studies of bristles although this could simply be due to a higher concentration other components in bristle procuticle interfering with our ability to detect the layering. A second difference is that the prominent undulae seen in most cells synthesizing cuticle were not detected in bristle forming cells (Adler, 2017). Further studies are needed to determine if the linkage between chitin and chitin synthase is a general result for arthropod epithelial cells. The close connection between CS and chitin is similar to that seen in yeast and fungi (Leal-Morales et al., 1994) (Chuang and Schekman, 1996) (Santos and Snyder, 1997) (Kozubowski et al., 2003) (Latge et al., 2005) and is reminiscent of the connection between cellulose and cellulose synthase in plants (Polko and Kieber, 2019).

The localization of Kkv in bristles requires both the intracellular transport of the protein into the plasma membrane and its restriction to stripes. We previously identified *Rab11* and *dyl* as being essential for the normal deposition of chitin in bands in bristles (Nagaraj and Adler, 2012). We established here that in the absence of *Rab11* function Kkv::NG failed to localize to the plasma membrane. In contrast, in the absence of *dyl* function Kkv::NG localized to the plasma membrane but not preferentially accumulate into the appropriate stripes. These results argue that these two genes mediate different steps essential for the localization of Chitin Synthase. Rab11 is also required for the insertion of Dyl into the plasma membrane of developing bristles (Nagaraj and Adler, 2012) and we suspect it has a general role for insertion of proteins into the shaft plasma membrane. The role of *dyl* is of particular interest. Dyl is a ZP (zona pellucida) domain protein and like other ZP domain proteins it can polymerize (Adler et al., 2013; Jovine et al., 2005; Jovine et al., 2002) and it is thought that this allows it to organize the apical extracellular matrix (Chanut-Delalande et al., 2012; Fernandes et al., 2010). The expression of *dyl* is almost entirely restricted to the period of envelop deposition (Sobala and Adler, 2016) and it accumulates in bands along the proximal distal axis of developing bristles (Nagaraj and Adler, 2012). Hence, it localizes in a way that is appropriate for instructing the later accumulation of Kkv in stripes. I observed that the stripes of Kkv::NG and Dyl were interdigitated and did not overlap. I suggest that that Dyl functions as a negative factor to inhibit the accumulation of Kkv::NG from regions of the bristle plasma membrane, but likely does so indirectly as there appears to be space between the interdigitated bands. Future studies will be needed to elucidate the mechanisms involved here (e.g. local Dyl could recruit a factor that removes Kkv::NG from nearby regions of the membrane). In addition to Rab11 and Dyl we also established that the large bundles of cross linked F-actin in bristles were also required for the normal deposition of chitin bands. The localization of both Dyl and Kkv::NG were altered in developing *sn f* bristles. The mislocalization of Dyl provides a mechanism for the mislocalization of Kkv::NG and the subsequent abnormal chitin deposition in *dyl* mutant bristles. A number of other genes have been identified that are required for normal chitin deposition or where a loss of function leads to a *kkv* like wing hair phenotype (Adler et al., 2013; Chaudhari et al., 2011; Moussian et al., 2015; Moussian et al., 2006b). It will be interesting to determine which, if any of these also mediate Kkv localization.

It is possible that Kkv is actively cycled from the plasma membrane to cytoplasmic endosomes/chitosomes and then back to the plasma membrane. There is strong evidence for the recycling of chin synthase in yeast (Hernandez-Gonzalez et al., 2018; Knafler et al., 2019; Sacristan et al., 2013) and Rab11 is a well-established marker for late endosomes (Calero-Cuenca and Sotillos, 2018) (Welz et al., 2014) and the recycling of membrane proteins. In such a model the failure of Kkv::NG localization in Rab11 deficient bristles could be due to a defect in recycling and not in the original localization. This might also explain the failure of the presumptive catalytic defective Kkv-R896K mutant protein to localize to the plasma membrane. It is possible that the inactive protein is more rapidly removed from the membrane and that it is preferentially not recycled back to the plasma membrane or recycled more slowly. This could be a quality control mechanism in the formation of insect cuticle.

### *kkv::smGFP* is useful as a system for structure function studies on CS

The importance of chitin synthase function for insects is demonstrated by the lethality associated even with moderately small clones of *kkv* mutant cells (Ren et al., 2005) (Adler et al., 2013) and knocking down *kkv* function for a restricted period of time in a limited set of epidermal cells (pna - unpublished). The edited *kkv::smFP* allele is the only homozygous viable hypomorpic allele of *kkv* that we are aware of. As we demonstrated the rescue of the wing hair phenotype of *kkv::smFP* is an easy assay for testing the functionality of mutant Kkv proteins. This assay relies on UAS-Gal4 driven expression and this could be misleading as overexpression could prevent distinguishing between mutants with reduced vs completely normal activity. There are however, advantages to this assay compared to editing the endogenous gene. It is important to consider that while CRISPR/Cas9 mediated editing is not difficult it still involves more time and labor than UAS transgenesis and mutations identified as interesting by the UAS-Gal4 system can later be assessed using CRISPR/Cas9 to test for reduced but still significant chitin synthase activity. Further, chitin synthases are known to function as multimers (Merzendorfer, 2011) (Gohlke et al., 2017) and some mutations might be dominant negatives. These could be identified using the UAS-Gal4 system but they would likely fail to be recovered by CRISPR/CAS9 mediated editing (or by classical mutagenesis) as they are likely to be dominant lethals. The UAS/Gal4 system could also be used to identify parts of the Kkv protein that are essential for its localization. The rescue by the Kkv-mos::NG protein indicates that the system should be able to assess at least in part the function of non-Drosophila chitin synthases.

It was not surprising that the *kkv R896K* mutant showed no rescue activity as this missense mutation is considered an amorphic allele in Drosohila (Moussian et al., 2005) and a similar substitution in yeast contained only about 1% of wild type activity (Nagahashi et al., 1995). The failure of Kkv R896K to show rescue activity validates the rescue system for structure function studies on the fly CS. It was surprising that this mutant protein did not localize properly. It is possible that the active site missense mutation disrupts both catalytic activity and normal protein folding and the folding defect leads to a failure to traffic the protein to the apical plasma membrane. As noted above it is also possible that the defect is not in the initial trafficking but is due to the inactive protein being removed more quickly. Further studies will be required to distinguish between these hypotheses.

The *kkv::NG* and *kkv-smFP* edits were in the same location in the genome so it is likely that the greater activity and fluorescence of the *kkv::NG* edit compared to the *kkv-smFP* edit is not due to differences in transcription. Rather, our data suggests the smFP tagged only accumulates to a much lower level than the NG tagged protein. This could be due to a reduced half-life of the smFP tagged protein or to it folding less efficiently. One possible cause of this is the presence of multiple copies of the HA epitope tag in smFP. A study in yeast reported that a 3XHA tag could cause a dramatic decrease in the accumulation of some of the tagged proteins (Saiz-Baggetto et al., 2017). It is possible that a similar phenomenon can explain our results with *kkv-smFP*.

### Kkv is present in the plasma membrane prior to procuticle formation

Previous studies on the transcriptome of pupal wing cells (Ren et al., 2005; Sobala and Adler, 2016) established that *kkv* RNA was present prior to the start of wing blade procuticle deposition. Part of the reason for this is that wing hair chitin is deposited earlier than wing blade chitin. However, this cannot explain the presence of *kkv* RNA 8 hrs prior to the start of hair morphogenesis and 16 hrs prior to the earliest time we can detect hair chitin (Ren et al., 2005) (Adler et al., 2013). Neither can it explain the presence of Kkv protein in the general apical membrane (i.e. not in the hair) more than 12 hrs prior to blade procuticle deposition. These observations suggest the possibility that Kkv has an earlier function in cuticle formation that is not due directly to chitin synthesis (e.g. a structural role for the protein) or that the synthesis of unstable chitin could be important prior to the time when it begins to accumulate. Both of these hypotheses suggest it might be possible to detect abnormalities at early stages of cuticle formation in *kkv* mutant cells.

### Similarities and differences in bristle and tracheal chitin deposition

Chitin deposition in bristles and the adult cuticle shows both similarities and differences from that described in trachea. In trachea the distribution of chitin changes during development. Starting out as a thick filament that largely fills the lumen it transforms into a thin zig zag shaped filament. The thin filament is eventually lost and during the this period chitin becomes concentrated over the distinctive tracheal taenidial folds (Devine et al., 2005; Ozturk-Colak et al., 2016; Tonning et al., 2005). It is not clear whether there is a complete loss of the chitin fibrils found in the central filament or if there is a reorganization of those fibrils into the taenidial fold chitin. In both tissues the disruption of the actin cytoskeleton results in an abnormal pattern of chitin; however in tracheal development a lack of chitin lead to an abnormal actin cytoskeleton while we did not see that in wing cells that lacked Kkv (Adler et al., 2013). In trachea Kkv puncta were seen more frequently over the taenidial folds than in the inter-fold region (Ozturk-Colak et al., 2016) but the patterning was less distinctive than we have seen in bristles. Some of the differences between these results could be due to the use of UAS-Gal4 to drive the expression of Kkv in trachea as in our hands using UAS-Gal4 to express *kkv* in bristles resulted in a “messier” pattern than was observed using the edited *kkv* gene. This is presumably due to UAS-Gal4 leading to overexpression.

## Methods and Materials

### Fly Stocks and Genetics

Flies were grown on standard fly food. They were routinely raised at 25°C, but in some experiments, they were raised at 21°C to slow development. In other experiments we used a temperature sensitive Gal80 to limit UAS transgene expression (McGuire et al., 2004). In these experiments, the animals were grown at 21 °C or 18 °C and then at the desired stage transferred to 29.5°C to inactivate the Gal80 and induce the expression of the UAS transgene. The various RNAi inducing transgenes came either from the VDRC (Dietzl et al., 2007) or TRiP collections (Perkins et al., 2015). The VDRC lines were obtained from the VDRC (http://stockcenter.vdrc.at/control/main). The TRiP lines were obtained from the Bloomington Drosophila Stock Center (http://flystocks.bio.indiana.edu/) (NIH P40OD018537) as were many other lines used in the research (e.g. Gal4 lines, Df stocks, *kkv*^*1*^ carrying stock). Flies that carried a *y w sn*^*3*^ *f*^*36a*^ X chromosome were kindly provided by G. Guild. Other stocks were made by the author in his lab.

### Constructs for generating transgenic lines

#### UAS constructs

The UAS constructs were in the pUAST-attb vector (Bischof et al., 2007). There are 3 *kkv* mRNA isoforms that encode two distinct *kkv* proteins (Thurmond et al., 2018). All of our experiments and analyses were done with the A isoform unless stated otherwise. The C protein isoform is identical to the A isoform and both contain 1615 aa. The D isoform also contains 1615 aa but it differs from the other two proteins by 14 aa due to its mRNA containing an alternative coding exon. The 14 amino acids are found in the region bounded by aa 1277 and 1322 of the A isoform. A comparison of the sequence of the genomic *kkv* gene and the longest *kkv* cDNA (RE32455) from the Drosophila genome project revealed two putative single base pair deletions in RE32455. A comparison of conceptual translation with those of other chitin synthases showed that the genomic sequence was correct. The two single base pair deletions were repaired by site directed mutagenesis to correspond to the genomic sequence. The cDNA was amplified and fused to the coding region for Neon Green kkv by Gibson assembly (NEB-E2611). This fusion gene was inserted into pUAST-attb using added Xho1 and Xba1 sites present in pUAST-attb and added to *kkv::NG* during construction using PCR and oligos containing the sites. This plasmid is referred to as *UAS-kkv::NG* (Addgene-138953). A similar strategy was used for the construct where the Neon Green tag was replaced by the ollas-his_6_ (OH) tag (Addgene - 138956). The nucleic acid sequences are provided in supplementary files S1 and S2 and the sequences of the tagged Kkv proteins are provided in files S3 and S4. *The UAS-kkv-R896K::NG* plasmid (Addgene – 138957) used for transgenesis was made by site directed mutagenesis of *UAS-kkv::NG*. Although an R to K substitution is generally considered a conservative substitution R896 is conserved in all chitin synthases and is thought to be at the catalytic site. In addition, the R to K change is found in the amorphic *kkv*^*1*^ mutation in Drosophila (Moussian et al., 2005). The same R to K mutation in yeast Chs2 resulted in a reduction to ∼ 1% of normal Chs2 enzyme activity (Nagahashi et al., 1995). The Kkv-mos::NG protein differs from Kkv::NG by a series of mutations that lead to 8 amino acid changes (in the CS-C domain (pfam 03142)) that are found in several mosquito species (e.g Aedes aegypti, Aedes albopictus, Anopheles gambiae str. PEST, Culex pipiens pallens, Anopheles quadrimaculatus, Anopheles sinensis). In Kkv-mos::NG the sequence from aa 702-909 is identical to the mosquito Chitin Synthase 1 proteins. The CS-C domain is from aa 722 to aa 904 in *kkv* and is slightly larger than a region of ScCHS2 that was shown to contain chitin synthase catalytic activity. The *UAS-kkv-mos-NG* plasmid (Addgene - 138958) was constructed from *UAS-kkv::NG* by synthesis of the relevant region and by it being placed into *kkv-NG* by Gibson assembly (this and several other DNA manipulations were done by EpochLifeSciences). The nucleic acid sequence of *kkv-mos::NG* is provided in sequence file S5 and the protein sequence in S6.

#### The HDR repair constructs

The upstream, middle and downstream repair regions were synthesized by assembly of oligonucleotides by EpochLifeSciences. The segments that comprised Neon Green and smGFP-HA were obtained by PCR from plasmids obtained from Allelebiotech and Addgene (#63166) respectively, added in the correct position by Gibson Assembly. The synthesized segment included several silent mutations to prevent re-cutting by Crispr/Cas9. The repair segments were subcloned into pHD-DsRed vector (Addgene plasmid #51434) resulting in pHD-DsRed-kkv-3 (Addgene – 138960 for the NG repair construct). The sequences of the plasmids that contain the HDR repair constructs are provided in files S7 and S8. The sequences of the Kkv proteins encoded by the two edited genes are provided in File S9 and S10. The construction of the edited genes resulted in a two amino acid linker (AG) between the C terminal aa of kkv and the first amino acid of NG (or smFP). A carton showing the strategy is provided in Fig 1.

##### gRNA constructs

Two plasmids that express the needed gRNAs were made by inserting oligonucleotides (files S11) into the pCFD3-dU6:gRNA plasmid where they would be expressed from the pU6-3 promoter (Addgene plasmid – 45946). These two plasmids are pCFD-pUG-DH1 (Addgene 138963) and pCFD-dUG-DO1 (Addgene - 138962).

##### Transgenic Lines

Injections of DNA into embryos were done by Rainbow Transgenics. The UAS transgenes were injected into embryos that contained the VK00033 attp landing site (cytol location 65B2; 3L:6,442,676..6,442,676). The transgenes were marked by a *w*^*+*^ gene and Go flies were crossed to w^1118^; TM3/TM2 flies and the progeny screened for eye color. G1 male flies with eye color were crossed to w; TM3/TM6 female flies and stocks were established by crossing siblings.

The HDR construct and the gRNA constructs were both injected into *nos-Cas9* expressing embryos (injections by RainbowTransgenics). The Go flies were crossed to w; TM3/TM2 flies and the G1 flies were screened for candidate edits by the expression of DsRed. Numerous putative edits were obtained by screening for RFP expression from the PhD-Ds-red vector used for HDR. Putative edit containing flies were crossed to w; TM2/TM3 flies and stocks established by crossing siblings that contained the TM3 balancer. The Ds-Red expression was monitored and proved to be useful in later stock constructions. We also generated fly stocks where the DsRed was removed by crossing edited male flies to *hs-cre; TM3/TM2* females and then crossing *hs-cre; kkv::NG + DS-Red/TM3* males to *w; TM3/TM2* females. The progeny from this cross were screened for *TM3* (and non-*TM2*) flies that did not express Ds-Red. Stocks were established from such single male flies and characterized by PCR to insure they carried the edited Kkv-NG gene but lacked Ds-red sequences. No phenotypic differences were observed between edited flies that carried or did not carry Ds-Red. The presence of Ds-Red expression was convenient for following the edited gene in crosses and it was used for some experiments where we were not imaging an alternative red fluorescent protein or stain.

##### Characterization of *kkv* edits

Six independent lines were established for both types of edits. DNA was isolated from these and assayed for the correct DNA changes by PCR followed by sequencing (the oligos used for these experiments are in Table S1). Most of the lines appeared to be as designed and resulted in the in frame fusion of the C terminus of Kkv and the fluorescent protein with the designed two amino acid linker. Three *kkv-NG* and two *kkv-smGFP-HA* lines were retained and further characterized. No differences were seen between the 3 NG edits and between the 2 smGFP edits. One line of each was chosen as the standard for routine use. Both of these are available at the Bloomington Drosophila stock center.

#### Confocal Microscopy

Immunostaining of fixed pupal epidermal cells during the deposition of cuticle is complicated by the inability of the antibodies to penetrate cuticle after the early stages of its development. Thus, most of the imaging experiments we carried out on Kkv in pupae were done by in vivo imaging of Kkv::NG. In a small number of experiments we examined Kkv-NG in fixed tissue. In some we simply used the inherent fluorescence of the neon green tag (sometimes combined with phalloidin staining of actin). In others we used anti-NG immunostaining. Such tissue was only weakly fixed and we did not use animals that were older than around 48 hr after white prepupae (awp). Otherwise, immunostaining of pupal and larval tissues were done as described previously (Nagaraj and Adler, 2012). Imaging of live Kkv::NG containing pupae was done on a Zeiss 780 confocal microscope in the Keck Center for Cellular Imaging. Stained samples were examined on the same microscope.

#### Comparison of *kkv::NG* and *kkv::smFP*

We estimated the brightness difference between the products of the *kkv::NG* and *kkv::smFP* edited genes by live imaging both in the same confocal session using the same microscope conditions. We measured the brightness of maximal projections of both types of animals and then subtracted the background brightness. The ratio of brightness for *kkv::NG* and *kkv::smFP* was 14.7. To estimate the relative amount of the two proteins present we needed to correct for the relative brightness of the two fluorescent protein tags. We were unable to find a value for the brightness of *smFP* but we were able to find a value for the progenitor of smFP, superfolder GFP and Neon-Green (Lambert and Thorn, 2019). The relative brightness was 1.7, which gave an estimate that Kkv::NG was present in 8.65 fold higher concentration than Kkv::smFP. Assuming Kkv::smFP has the same specific activity as wild type Kkv we estimate the *kkv::smFP* cells only contain about 11% of the normal Kkv enzyme activity.

#### Kkv topology experiments

We obtained predictions for the number and locations of transmembrane domains from the following programs: TMHMM2.0, TMPRED, Uniprot, PHDhtm and CCTOP. The CCTOP site returned predictions for HMMTOP, Memsat, Octopus, Philius, Phobius, Pro, Prodiv, Scampi, ScampiMsa as well as CCTOP. 14 putative transmembrane domains were predicted by 13 or 14 of these 14 programs. These “consensus sites are shown in Fig 6 and the specific TMHMM2.0 predictions are provided in Table S2. The specific amino acids predicted to be in each transmembrane domain were often shifted by a few amino acids by different programs but the putative transmembrane domains substantially overlapped.

To examine the topology of Kkv we drove the expression of *UAS-kkv-OH* or *UAS-kkv::NG* by *ptc-GAL4*. This results in a stripe of expression along the anterior/posterior compartment boundary in wing discs. Wing discs were dissected in PBS and fixed in the cold for 15’. The discs were then manually cut or punctured to ensure the apical surface of the epithelial cells was exposed to antibody. They were then incubated for 30’ in PBS supplemented with 10% Sheep Serum. The discs were then stained overnight at 4°C in PBS, 10% sheep serum plus the desired antibody. The discs were then rinsed 4 times in PBS and then stained with secondary antibody for 3 hrs at room temperature in PBS, 10% sheep serum plus secondary antibody. After 4 rinses in PBS the discs were washed with PBST (PBS plus .3% triton X100) followed by three additional washes in PBS. Finally the discs were mounted in ProLong Diamond. As a control several of the fixed and cut discs had PBST substituted for PBS in all steps in the experiment. The wing discs were examined on a Zeiss Axioskop II and photographed on a Spot Digital Camera (National Diagnostics).

## Supporting information

all supplemental files

## Acknowledgements

This research was supported by funds provided by the W. R. Kenan Chair to the author and limited personal funds of the author. The author thanks H.S. Tzu for helpful conversations. “We acquired confocal images using the Keck Center Zeiss 780 Confocal microscopy system (NIH OD016446). We acquired Scanning Electron Microscope images at the Advanced Microscopy Facility at the University of Virginia. The images were obtained on a Zeiss VP HD SEM field purchased with a grant from the NIH (NIH 1S10OD011966). The author is retiring and closing his laboratory in May 2020. Hence, any requests for reagents or additional information should be made prior to then.

