## Supplementary material for "The localization of chitin synthase mediates the patterned deposition of chitin in developing Drosophila bristles": all supplemental files

**Supplemental Figure Legends.**

Fig S1. The melanotic spot phenotype. A. No melanotic spot phenotype is seen in hypomorphic *kkv::smFP/Df* flies. B. The over expression of the catalytically inactive Kkv R896K mutant driven by *ap-Gal4* results in the melanotic spot (arrow) phenotype. C. The over expression of Kkv::NG driven by *ap-Gal4* results in the melanotic spot (arrow) phenotype.

Fig S2. SEM examination of wing cuticle of edited flies. A. Flies homozygous for *kkv::smFP* show the thin bent and curved wing hair phenotype. B. *kkv::smFP/Df* flies show a slightly enhanced wing hair phenotype. C. Flies homozygous for *kkv::NG* have normal wing hairs. D. *Kkv::NG/DF* flies also have normal wing hairs. E. Oregon R wild type flies have normal wing hairs. F. Ore-R/Df flies also have normal wing hairs showing that *kkv* is not haplo-insufficient.

Fig S3. *kkv::NG* is much brighter than *kkv::smFP.* A. Image of live *kkv::NG* thorax (maximal projection) as obtained from the confocal with no modifications. The image was obtained with a lower laser power than we typically used to increase the range of brightness. B image of live *kkv::smFP* as obtained from the confocal using the same settings as in A. C. The image from A brightened in Photoshop by +150. D. The image in B brightened in Photoshop by +150. E. The image in C brightened by a further +150. F. The image from D brightened by a further +150. G. The image from E with both the brightness and contrast increased by 150. H. The image from E with both the brightness and contrast increased by 150. Hairs on the notum are finally visible and a bristle is barely visible. The asterisks mark pupal cuticle. The *kkv::NG* animal was a few hrs younger (48 hr awp) than the *kkv::smFP* animal (52 hr awp). Since 52 hr animals show slightly higher fluorescence this figure slightly underestimates the difference in brightness.

Fig S4. Comparison of the subcellular localization of Kkv::NG and Kkv R896K::NG. A. Image of live 44hr ap>*kkv::NG* pupal wing. B. Image of live 44hr ap>*kkv-R896K::NG* pupal wing. C. Image of a 36hr ap>*kkv::NG* pupal wing fixed and stained with both anti-NG antibody (green) and Alexa 568 phalloidin (red) to shoe F-actin. D. Image of a 36hr ap>*kkv R896K::NG* pupal wing fixed and stained with both anti-NG antibody (green) and Alexa 568 phalloidin (red) to shoe F-actin. E. Image of a live 52 hr ap>*kkv::NG* pupal wing showing a bristle with Kkv stripes. The arrows point to shed fluorescent puncta. F. Image of a live 52 hr ap>*kkv R896K::NG* pupal wing showing a bristle with abnormal accumulation of Kkv. G. Orthogonal cross sections of bristles from *ap>kkv::NG* live pupae. H. Orthogonal cross sections of bristles from *ap>kkv R896K::NG* live pupae. Note the stripes in G and the mislocalized Kkv in H. I. A live dissected salivary gland from a *ptc-Gal4/+; UAS-kkv::NG* third instar larva. The Kkv::NG preferentially accumulates on the apical surface of the gland (arrow) next to the lumen (asterisk). I' is a blow up of a single optical section of a *ptc-Gal4/+; UAS-kkv::NG* salivary gland. Note the many round vesicles with puncta obvious in the membrane of these vesicles (arrow). J. A live dissected salivary gland from a *ptc-Gal4/+; UAS-kkv R896K::NG* third instar larva. The Kkv R896K::NG does not preferentially accumulate on the apical surface of the gland next to the lumen (asterisk). Rather, it is rather evenly distributed across the cytoplasm. J'. The blow up shows a different morphology in how the protein accumulates compared to Kkv::NG (I').

Fig S5. Specificity of Kkv staining. A. A *ptc-Gal4/UAS-ChtVis; UAS-kkv::NG* wing disc shows the accumulation of Kkv::NG in the *ptc* domain (arrow). The secreted ChtVis (red) diffuses all over the disc in the extracellular domin that separates the disc epithelium and the peripodial membrane. B. Kkv::NG preferentially accumulates on the apical surface of disc cells (arrow). C-F. Comparisons of *kkv::NG* (CE) and Ore-R (DF) 63 hr awp pupae by live imaging. C and D were taken at the same microscope settings as were E and F. Note the bristle, hair and cell membrane fluorescence of Kkv::NG is specific and not seen in Ore-R. The arrows in C and D point to auto-flourescent pupal cuticle. E and F are higher magnification images that show the stripes of Kkv::NG in bristles and the lack of such fluorescence in Ore-R.

Fig S6. Kkv::NG fluorescence as a function of developmental time since white prepuae formation. All images were taken at the same microscope setting.

Fig S7. How geometry affects optical sectioning of bristles. A cartoon of a bristle (B) in cross section is shown with intracellular bands of red and extracellular bands of green. The arrows indicate different planes of sectioning and the images the arrow point to show the expected resulting images and how these can differ depending on the location and orientation of the optical sections.

Fig S8. The accumulation of Dyl in stripes in bristles is altered in sn f double mutants. A. A *sn f* bristle immunostained with anti-Dyl antibodies. Note the parallel lines of Dyl staining (Nagaraj and Adler, 2012) are disrupted. B. Cross sections of several *sn f* bristles immunostained for Dyl. Note the disruption in staining and the retention of lines of Dyl is variable from bristle to bristle.

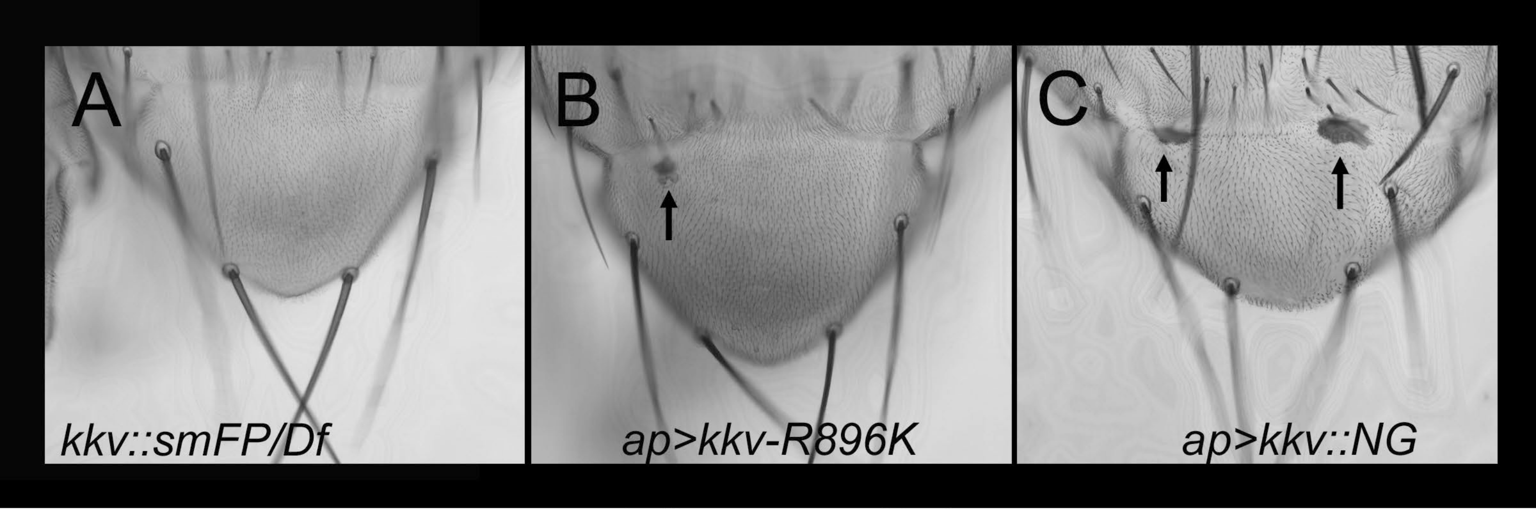


Figure S1

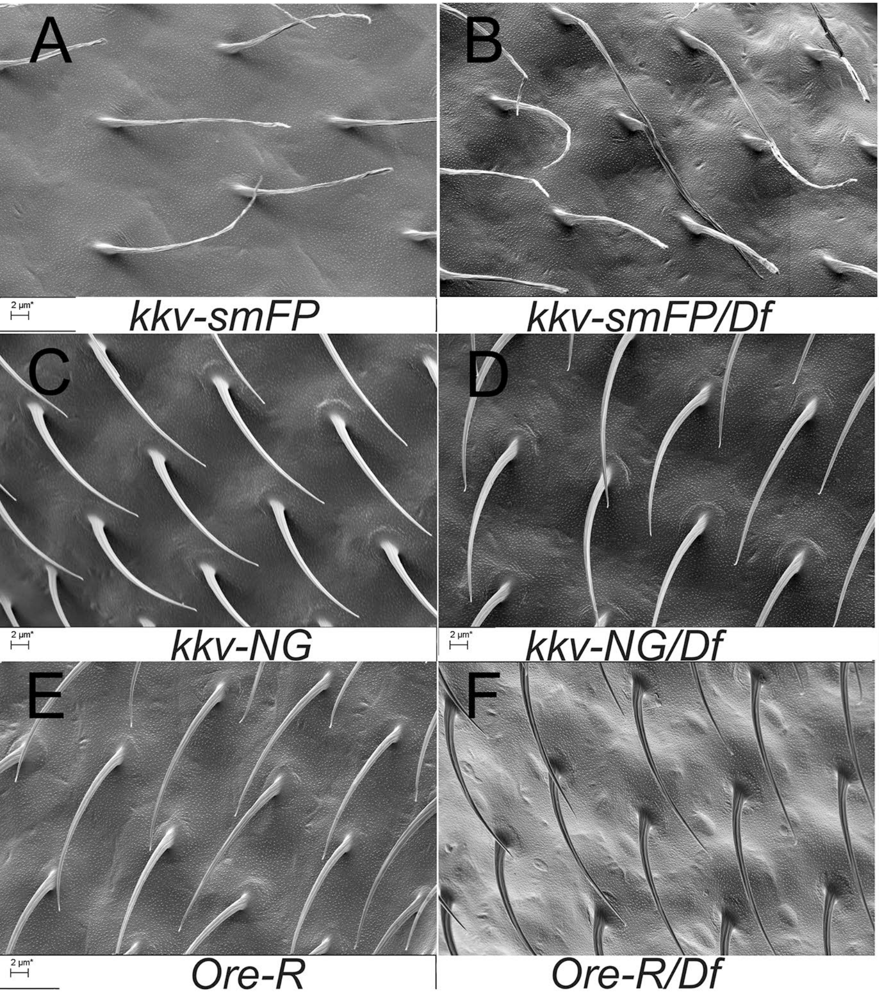
A
B
2 µm*
kkv-smFP
kkv-smFP/Df
C
D
2 µm*
kkv-NG
2 µm*
kkv-NG/Df
E
F
2 µm*
Ore-R
Ore-R/Df

Figure S2

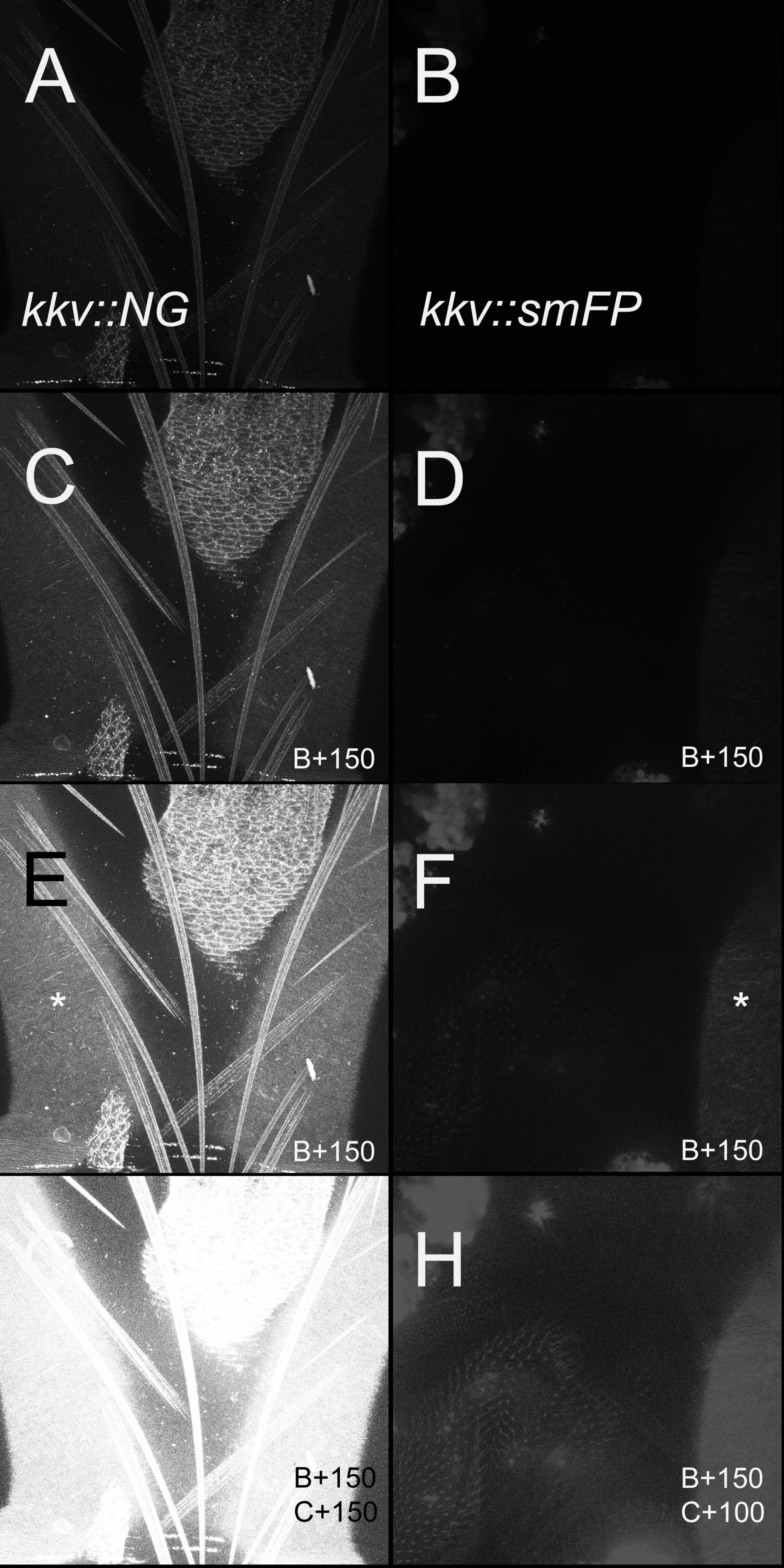
A
kkv::NG
B
kkv::smFP
C
B+150
D
B+150
E
*
B+150
F
*
B+150
H
B+150
C+150
B+150
C+100

Figure S3

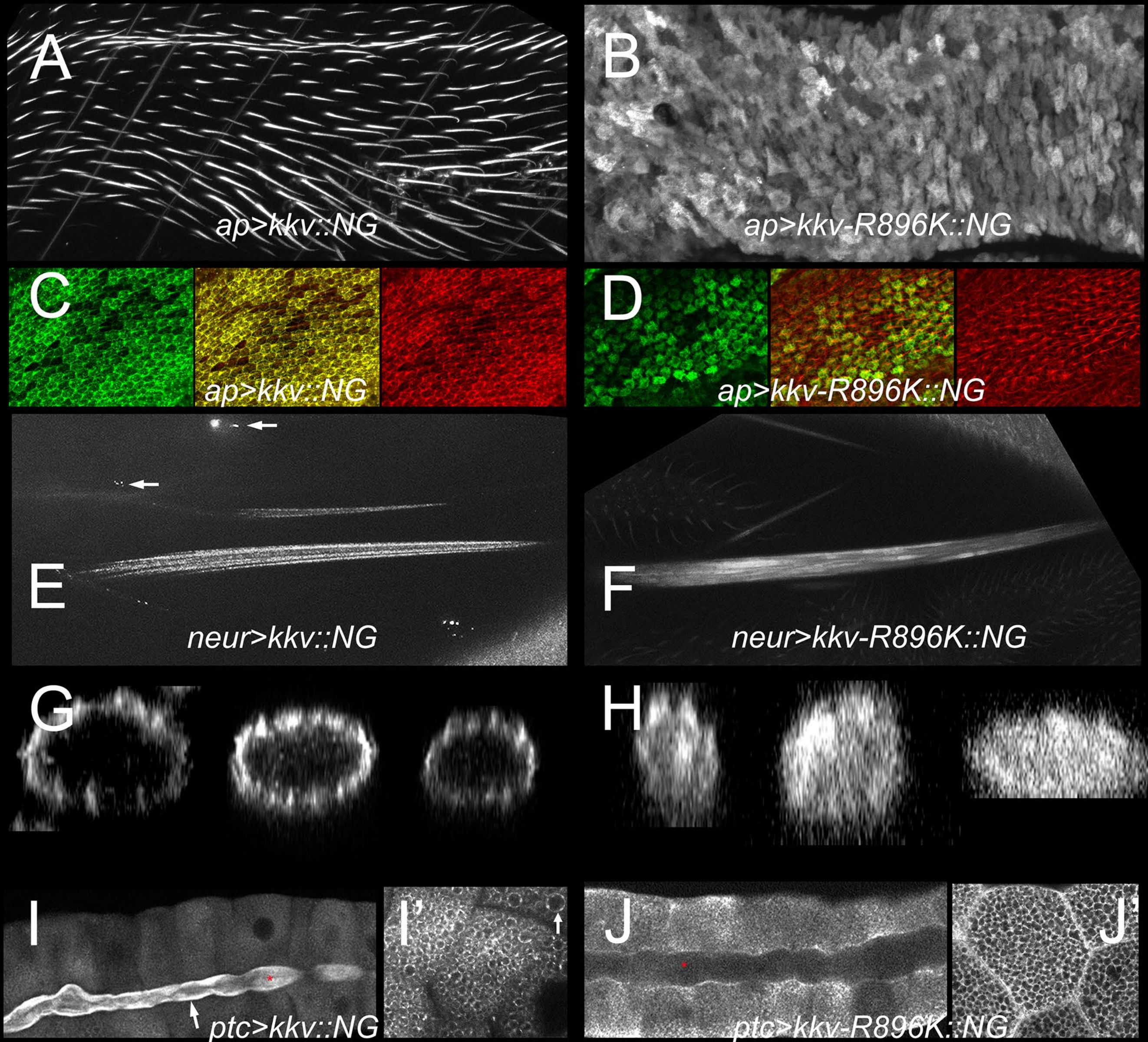
A
ap>kkv::NG
B
ap>kkv-R896K::NG
C
ap>kkv::NG
D
ap>kkv-R896K::NG
E
neur>kkv::NG
F
neur>kkv-R896K::NG
G
H
I
I'
ptc>kkv::NG
J
J'
ptc>kkv-R896K::NG

Figure S4

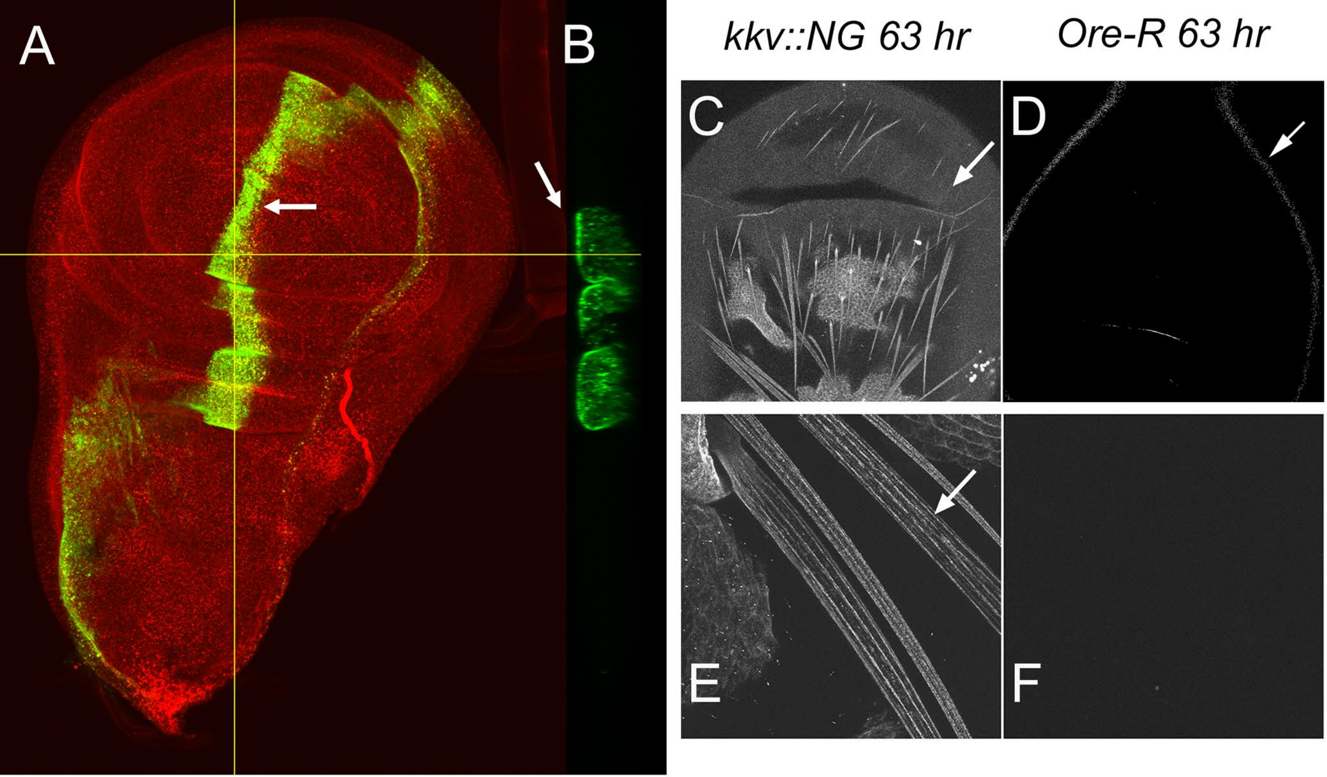


Figure S5

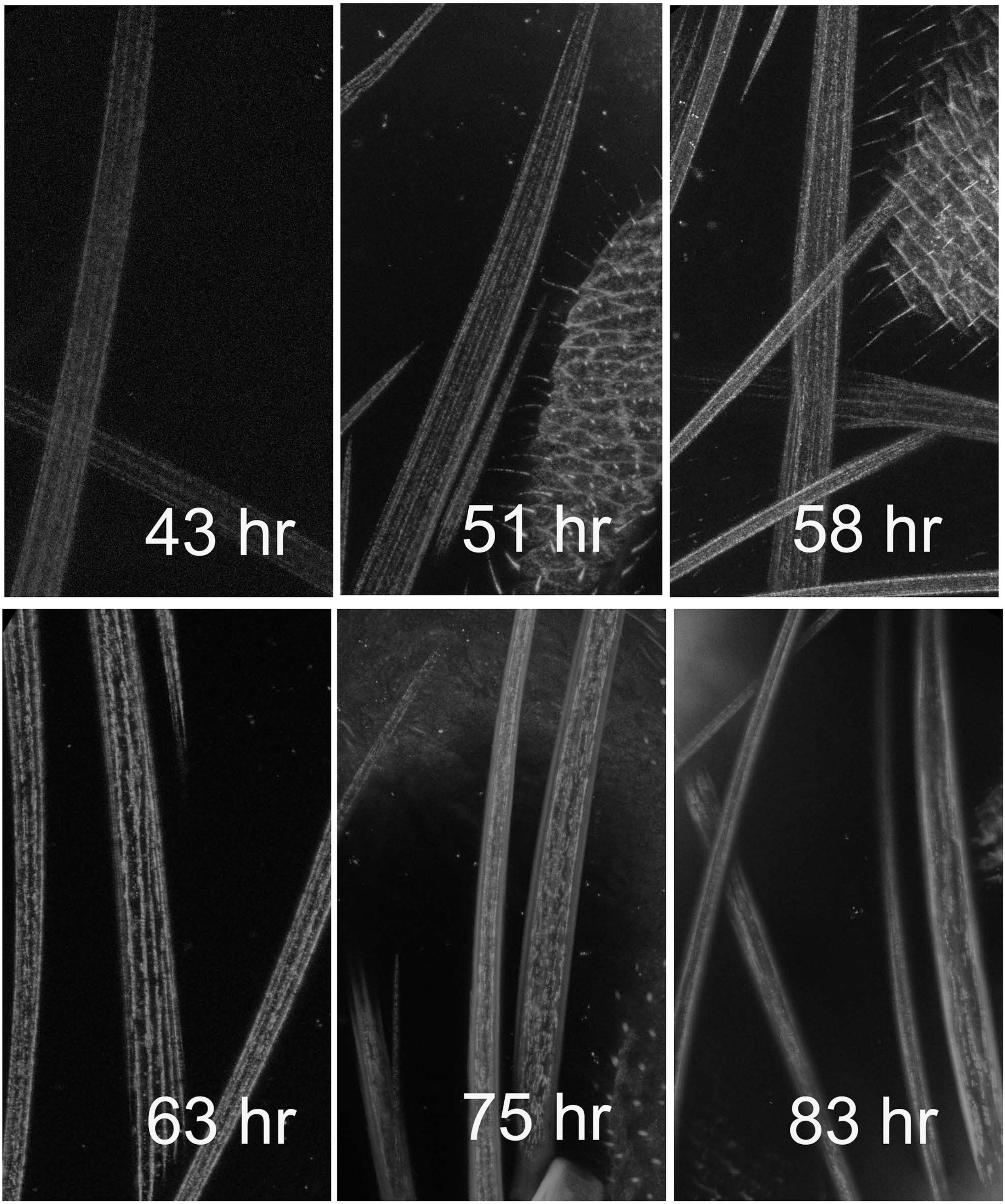


Figure s6

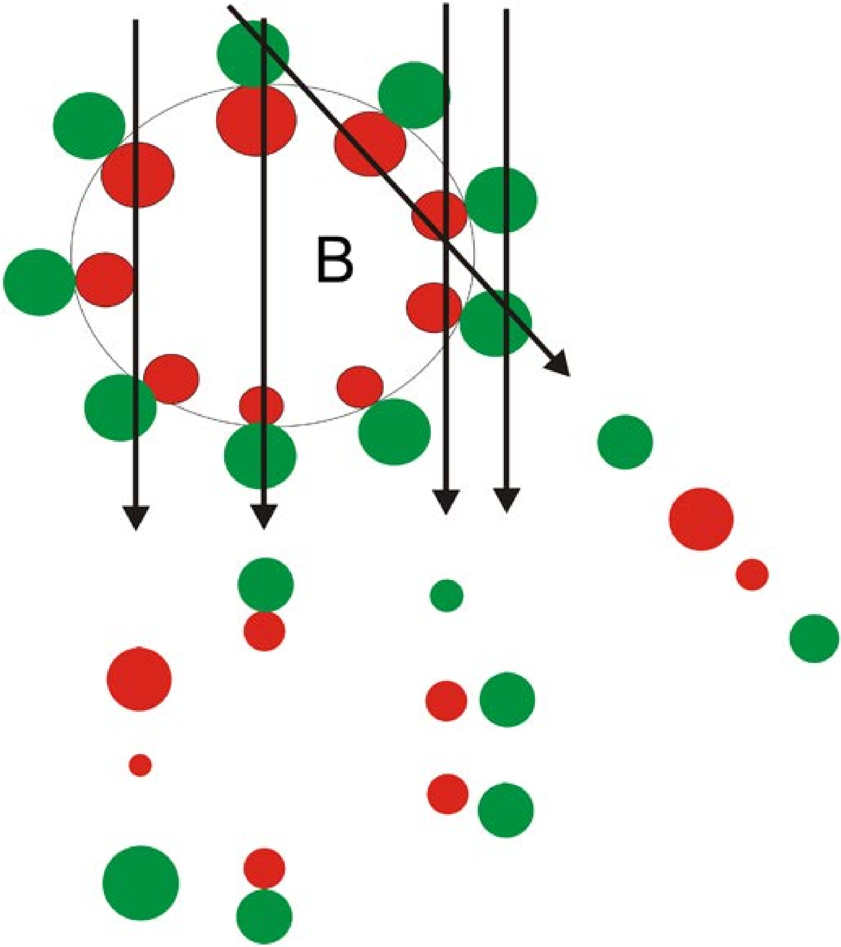


Fig s7

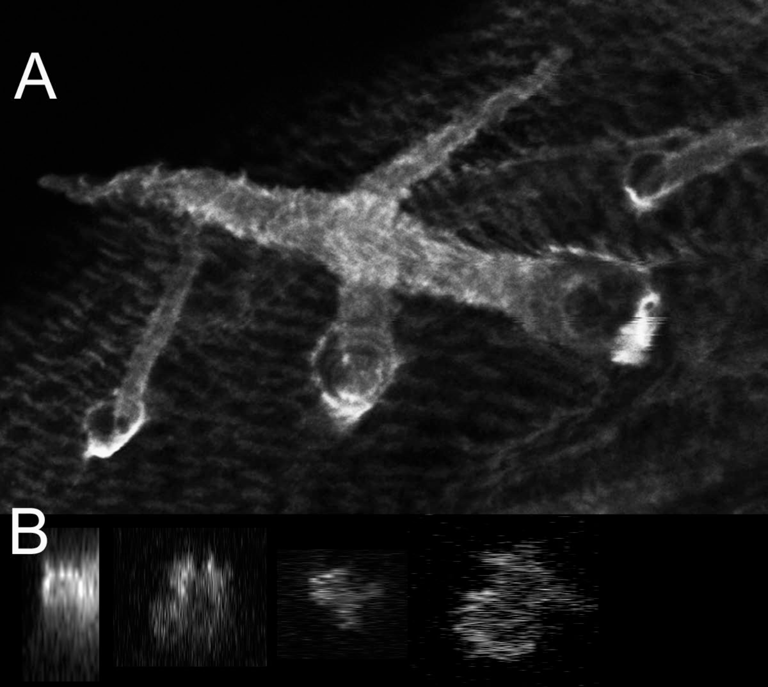


Fig s8

Table S1

| name | sequence | Location | Comment |
|---|---|---|---|
| kkv_seq1 | AACTGGAGCACCGGTGAGATTA | 3R 5379079-5379058 | in 9th exon of kkv |
| kkv_seq2 | TAAGCCAATACCTAGAATTG | 3R 5378544-5378525 | in 11th exon of kkv - 3'UTR |
| kkv_seq3 | GCCGGAGTATAAATAGAGG | 4068-4086/3729-3747 | hdr-DsRed-smFP/hdr-ng, in vector |
| kkv_seq4 | CACAAGGCCCTGAAGCTGAAGG | 4696-4717/4357-4378 | pHDR-DsRed-smFP/pHDR-DsRed-ng, in vector |
| kkv_HA_seq1 | CGTATGACGTGCCAGACTACGCGG | 2349-2372 | pHDR-DsRed-smFP, in one of the HA coding regions |
| kkv_HA_seq2 | GCCATAATGTGTACATTACG | 2817-2836 | pHDR-DsRed-smFP, in superfolder GFP coding region |
| kkv neongreen_seq | TCTGGTCCCTCATATCGGGTAT | 2467-2488 | pHDR-DsRed-NG - in neongreen coding |
| pcr rev | AACCTCTACAAATGTGGTATGGCTG | 4938-4914/4599-4575 | pHDR-dsRed-smFP/pHDR-DsRed-NG - downstream of DsRed coding seq in vector |
| pcr fwd | CGTATTATCATCCCCCATAGGTGC | 5379711-5379687 | kkv genomic - in intron just upstream of 8th exon |

Table S2

| Predicted TMD | aa start | aa end | Prediction | antibody | comment |
|---|---|---|---|---|---|
| | 1 | 89 | inside | A - by experiment inside | amino terminus |
| 1 | 89 | 111 | | | |
| | 112 | 144 | outside | | |
| 2 | 145 | 164 | | | |
| | 165 | 182 | inside | | |
| 3 | 183 | 205 | | | |
| | 206 | 209 | outside | | |
| 4 | 210 | 232 | | | |
| | 233 | 315 | inside | | |
| 5 | 316 | 338 | | | |
| | 339 | 385 | outside | | |
| 6 | 386 | 408 | | | |
| | 409 | 420 | inside | | |
| 7 | 421 | 443 | | | |
| | 444 | 476 | outside | | |
| 8 | 477 | 496 | | | |
| | 497 | 923 | inside | P2 - by experiment inside | |
| 9 | 924 | 946 | | | |
| | 947 | 960 | outside | | |
| 10 | 961 | 980 | | | |
| | 981 | 986 | inside | | |
| 11 | 987 | 1006 | | | |
| | 1007 | 1015 | outside | | |
| 12 | 1016 | 1038 | | | |
| | 1039 | 1038 | inside | | |
| 13 | 1043 | 1065 | | | |
| | 1066 | 1287 | outside | M - by experiment inside | |
| 14 | 1288 | 1305 | | | |
| | 1306 | 1344 | inside | | |
| 15 | 1345 | 1367 | | | |
| | 1368 | 1615 | outside | anti-NG by experiment outside | C terminus |
| | | | | anti-ollas by experiment outside | |

Supplemental Text File 1

When expressed using *ap-Gal4* as a driver only modest phenotypes were seen were seen with *UAS-kkv::NG* or *UAS-kkv-OH* and these were also seen at a lower frequency in *ap-Gal4* flies that did not contain a *UAS-kkv* transgene. The most distinctive were small regions of melanized cuticle seen primarily near the suture connecting the scutum to the scutellum on the notum (Fig S1). This phenotype is very rare in *ap-Gal4* flies. The basis for this phenotype is unclear as it does not appear to be a straight forward gain of function as an equivalent phenotype is seen *in ap-Gal4/+; Df(3R)ED5156/+* flies (Df(3R)ED5156) deletes the *kkv* gene) (Dos Santos et al., 2015; Ryder et al., 2007). Further, the phenotype associated with *ap-Gal4* driven expression of *kkv::NG* was neither enhanced nor suppressed when a copy of *kkv* was deleted. We also observed the loss (or less frequently deformation) of one or more scutellar or dorso-central bristles in these flies and a malformed scutum, but these were also seen to a lesser extent in *ap-Gal4* flies and to a similar extent in *ap>kkv::NG* and *ap-Gal4/+; Df(3R)ED5156)/+* flies. It is not surprising that our UAS transgenes do not show dramatic gain of function phenotypes as Moussian and colleagues (Moussian et al., 2015a) have established that Kkv requires a second protein (MH2 domain containing - either Reb or Exp) for the accumulation of chitin.

File S1
pUAST-kkv::NG NA sequence.

ctgacgcgccctgtagcggcgcattaagcgcggcgggtgtggtggttacgcgcagcgtgaccgctacacttgccagcgcc
ctagcgcccgctcctttcgctttcttcccttcctttctcgccacgttcgccggctttccccgtcaagctctaaatcgggg
gctccctttagggttccgatttagtgctttacggcacctcgaccccaaaaaacttgattagggtgatggttcacgtagtg
ggccatcgccctgatagacggtttttcgccctttgacgttggagtccacgttctttaatagtggactcttgttccaaact
ggaacaacactcaaccctatctcggtctattcttttgatttataagggattttgccgatttcggcctattggttaaaaaa
tgagctgatttaacaaaaatttaacgcgaattttaacaaaatattaacgcttacaatttccattcgccattcaggctgcg
caactgttgggaagggcgatcggtgcgggcctcttcgctattacgccagctggcgaaagggggatgtgctgcaaggcgat
taagttgggtaacgccagggttttcccagtcacgacgttgtaaaacgacggccagtgaattgtaatacgactcactatag
ggcgaattgggtacgtaccgggcccctagtatgtatgtaagttaataaaacccatttttgcggaaagtagataaaaaaaa
catttttttttttttactgcactggatatcattgaacttatctgatcagttttaaatttacttcgatccaagggtatttga
tgtaccaggttctttcgattacctctcactcaaaatgacattccactcaaagtcagcgctgtttgcctccttctctgtcc
acagaaatatcgccgtctctttcgccgctgcgtccgctatctctttcgccaccgtttgtagcgttacgtagcgtcaatgt
ccgccttcagttgcattttgtcagcggtttcgtgacgaagctccaagcggtttacgccatcaattaaacacaaagtgctg
tgccaaaactcctctcgcttcttatttttgtttgtttttttgagtgattggggtggtgattggttttgggtgggtaagcag
gggaaagtgtgaaaaatcccggcaatgggccaagaggatcaggagctattaattcgcggaggcagcaaacacccatctgc
cgagcatctgaacaatgtgagtagtacatgtgcatacatcttaagttcacttgatctataggaactgcgattgcaacatc
aaattgtctgcggcgtgagaactgcgacccacaaaaatcccaaaccgcaattgcacaaacaaatagtgacacgaaacaga
ttattctggtagctgttctcgctatataagacaattttttgagatcatatcatgatcaagacatctaaaggcattcatttt
cgactatattctttttttacaaaaaatataacaaccagatattttaagctgatcctagatgcacaaaaaataaataaaagt
ataaacctacttcgtaggatacttcggggtactttttgttcggggttagatgagcataacgcttgtagttgatatttgag
atcccctatcattgcagggtgacagcggagcggcttcgcagagctgcattaaccagggcttcgggcaggccaaaaactac
ggcacgctccggccacccagtccgccggaggactccggttcagggagcggccaactagccgagaacctcacctatgcctg
gcacaatatggacatctttggggcggtcaatcagccgggctccggatggcggcagctggtcaaccggacacgcggactat
tctgcaacgagcgacacataccggcgcccaggaaacatttgctcaagaacggtgagtttctattcgcagtcggctgatct
gtgtgaaatcttaataaagggtccaattaccaatttgaaactcagtttgcggcgtggcctatccgggcgaacttttggcc
gtgatgggcagttccggtgccggaaagacgaccctgctgaatgcccttgcctttcgatcgccgcagggcatccaagtatc
gccatccgggatgcgactgctcaatggccaacctgtggacgccaaggagatgcaggccaggtgcgcctatgtccagcagg
atgacctctttatcggctccctaacggccagggaacacctgattttccaggccatggtgcggatgccacgacatctgacc
tatcggcagcgagtggcccgcgtggatcaggtgatccaggagctttcgctcagcaaatgtcagcacacgatcatcggtgt
gcccggcagggtgaaaggtctgtccggcggagaaaggaagcgtctggcattcgcctccgaggcactaaccgatccgccgc
ttctgatctgcgatgagcccacctccggactggactcatttaccgcccacagcgtcgtccaggtgctgaagaagctgtcg
cagaagggcaagaccgtcatcctgaccattcatcagccgtcttccgagctgtttgagctctttgacaagatccttctgat
ggccgagggcagggtagctttcttgggcactcccagcgaagccgtcgacttcttttcctagtgagttcgatgtgtttatt
aagggtatctagcattacattacatctcaactcctatccagcgtgggtgcccagtgtcctaccaactacaatccggcgga
cttttacgtacaggtgttggccgttgtgcccggacgggagatcgagtcccgtgatcggatcgccaagatatgcgacaatt
ttgctattagcaaagtagcccgggatatggagcagttgttggccaccaaaaatttggagaagccactggagcagccggag
aatgggtacacctacaaggccacctggttcatgcagttccgggcggtcctgtggcgatcctggctgtcggtgctcaagga
accactcctcgtaaaagtgcgacttattcagacaacggtgagtggttccagtggaaacaaatgatataacgcttacaatt
cttggaaacaaattcgctagattttagttagaattgcctgattccacacccttcttagtttttttcaatgagatgtatag
tttatagttttgcagaaaataaataaatttcatttaactcgcgaacatgttgaagatatgaatattaatgagatgcgagt
aacattttaatttgcagatggttgccatcttgattggcctcatctttttgggccaacaactcacgcaagtgggcgtgatg
aatatcaacggagccatcttcctcttcctgaccaacatgacctttcaaaacgtctttgccacgataaatgtaagtcttgt
ttagaatacatttgcatattaataatttactaactttctaatgaatcgattcgatttaggtgttcacctcagagctgcca
gtttttatgagggaggcccgaagtcgactttatcgctgtgacacatactttctgggcaaaacgattgccgaattaccgct
ttttctcacagtgccactggtcttcacggcgattgcctatccgatgatcggactgcgggccggagtgctgcacttcttca
actgcctggcgctggtcactctggtggccaatgtgtcaacgtccttcggatatctaatatcctgcgccagctcctcgacc
tcgatggcgctgtctgtgggtccgccggttatcataccattcctgctctttggcggcttcttcttgaactcgggctcggt
gccagtatacctcaaatggttgtcgtacctctcatggttccgttacgccaacgagggtctgctgattaaccaatgggcgg
acgtggagccgggcgaaattagctgcacatcgtcgaacaccacgtgccccagttcgggcaaggtcatcctggagacgctt
aacttctccgccgccgatctgccgctggactacgtgggtctggccattctcatcgtgagcttccgggtgctcgcatatct
ggctctaagacttcggcccgacgcaaggagtagccgacatatatccgaaataactgcttgtttttttttttttaccatta
ttaccatcgtgtttactgtttattgccccctcaaaaagctaatgtaattatatttgtgccaataaaaacaagatatgacc
tatagaatacaagtatttccccttcgaacatccccacaagtagactttggatttgtcttctaaccaaaagacttacacac
ctgcataccttacatcaaaaactcgtttatcgctacataaaacaccgggatatatttttatatacatacttttcaaatc
gcgcgccctcttcataattcacctccaccacaccacgtttcgtagttgctctttcgctgtctcccacccgctctccgcaa
cacattcaccttttgttcgacgaccttggagcgactgtcgttagttccgcgcgattcggttcgctcaaatggttccgagt
ggttcatttcgtctcaatagaaattagtaataaatatttgtatgtacaatttatttgctccaatatatttgtatatattt
ccctcacagctatatttattctaatttaatattatgactttttaaggtaatttttttgtgacctgttcggagtgattagcg
ttacaatttgaactgaaagtgacatccagtgtttgttccttgtgtagatgcatctcaaaaaaatggtgggcataatagtg
ttgtttatatatatcaaaaataacaactataataataagaatacatttaatttagaaaatgcttggatttcactggaact

aggctagcataacttcgtataatgtatgctatacgaagttatgctagcggatccaagcttgcatgcctgcaggtcggagt
actgtcctccgagcggagtactgtcctccgagcggagtactgtcctccgagcggagtactgtcctccgagcggagtactg
tcctccgagcggagactctagcgagcgccggagtataaatagaggcgcttcgtctacggagcgacaattcaattcaaaca
agcaaagtgaacacgtcgctaagcgaaagctaagcaaataaacaagcgcagctgaacaagctaaacaatctgcagtaaag
tgcaagttaaagtgaatcaattaaaagtaaccagcaaccaagtaaatcaactgcaactactgaaatctgccaagaagtaa
ttattgaatacaagaagagaactctgaatagggaattgggaattcgttaacagatctgcggccgcggctcgagatgtctg
cgatgcggcatcgcccgatggcccctccgggccaaggaccaggagcgggaaccgctggggagcacgtcgacagcgatgac
aacaactttaccgacgacgagagctcgcccctcacccacgacatttacggcggcagccaacgcaccatacaagagacgaa
gggttgggatgttttccgggatccgccaatcaaaattgagaccggatcgacggcgaaccaggaatgcttagaattaaccg
tgaagatcttgaagatcttcgcctatgtgattacgtttataatagttttaaccggtggcgtcatcgccaagggcacaatg
ctcttcatgacttcgcaggtgcgcaaggataagaagatggagtactgcaacaaagacttgggtcgggacaaaagcttcgt
agtccgacttccggaggaggagcgagtggcctggatctgggctctgctcatagcatatgcccttcccgagatcggagcct
tgatccgatccgcccgcatctgcttcttcaagacctttaaggtgccgaagacgggccacttcctcttcgtctggctgatg
gagagcttgagtgccgtgggcatggcgctgctgatgttcgtggttctgccccagatcgacgccatccagggcgccatgct
aaccaactgcctgtgcgtcgtgccgggcatcttcggccttctatcgcgcacctccaaggagggcaagcgctttgtaaagg
tgatcatcgatctggcggcggtggcggcccaggtgacgggcctggtgatctggccgctgctggagaaccgccgtgagctg
tgggtcatcccggtggcctgtgtcatgatctcgtgcggctggtgggagaattacgtgtcgccacagtccccgctcggcct
ggttcgagccctgggtcgcatcaaggaggagatgaagtacactcgctacttctgccacatattcctatccatctggaaga
tcctgcttttcttcaccgtcacactgctcatctactgggcgcaaggcgaggaacccggcaatctgtttgccatgtacgga
gatgcctttggaccccacaagatcatagtctacgaactgccagcgggactgggcggtgtgcttccagataccctggaatc
ggccaacattgacaccgtggatgtggatgccgcctacaatacggtggtgtatgtactgcttctacagattttcggcgcct
acttgtgctacatcttcggaaagttcgcctgcaagatcctcatccagggattcagctacgcgtttcccgtcagtctaact
gttccgctctctgttacgttcctgatcgcagcctgtggaatccggatcgacgatccctgcttcttccatgacaccattcc
ggactatctgttcttcacgagcccctcgaacttccggttcaacaactttgtcaccgagcaaatggcgtgggcctggatcc
tttggctcctcagtcagacctggatagcactgcacatctggactccgaagtgcgagcgtctggccaccaccgagaagttg
tttgtccagcccatgtactcctctctactgatcgatcagtcgatggccctcaaccgaaggcgcgatgaccaggcagatgt
gaaaacagaggatctctcggagatcgaaaaggaaaagggcgatgagtactacgagaccatttctgtgcacacggatcgct
cctcggcacccaacaagccatcgattaagtcctccgacaacatcacacgaatctactcctgcgcgaccatgtggcatgag
acgaaggacgagatgatagagttcttgaagagcatcatgcgaatggacgaggaccagtgtgctcgtcgcgtggctcaaaa
gtatctgagggttctcgatccggactattatgaatttgagacccacatcttcttcgatgacgccttcgaaatctccgatc
acagcgacgatgatatccagtgcaatcgattcgtcaagctcttgatagccaccatggacgaggctgcctccgagatccat
cagaccacgatcagactgcgtccgcccaagaagtatcccactccgtacggaggccgcctggtgtggacccttccagggaa
gaccaaattcattacgcacttaaaggacaaggaccgcattcgtcacaggaagcgttggtctcaggtcatgtacatgtatt
acctgctgggacatcgtctaatggagctgcccatttcggtggatcgcaaggatgtaatggcggagaacacctaccttttg
accctggacggagatattgacttcaacccgagtgcggtgactctactggtggatctgatgaagaagaacaagaacctggg
tgctgcctgcggtcgcattcatcccatcggatcgggacccatggtgtggtaccaaaaattcgaatacgccatcggccatt
ggctgcaaaaggccacggagcacatgattggctgcgtgctctgttcacctggatgcttctccctgttcaggggcaagggc
ctcatggatgacaatgtgatgaggaaatacacgacacggtcggatgaggctcgtcactacgtgcagtacgatcagggcga
ggatcgttggctgtgcacattgctcctccagaggggataccgcgtggagtactcggctgccagtgatgcgtacacccact
gccccgagggattcaatgagttctacaaccagcggcgcagatgggtgccctcgaccatagccaacatcatggatcttctg
gcggatgcgaagcgcacgatcaagatcaatgacaacatttcgcttctgtacatcttctaccagatgatgttgatgggcgg
caccatcctgggacccggaaccattttccttatgttggtgggtgcattcgtggcagcctttcgcattgacaactggacct
ccttccactacaacatcgtgccgatcctggccttcatgtttatctgcttcacctgtaagtcgaacatccagttgttcgtg
gcccaagtcctttcgacggcatatgccctgatcatgatggcggtgattgtgggtacggccttgcagttgggagaggacgg
aataggctccccttcggccattttcctgatatccatggtgggctcattcttcatagccgcatgtttgcatccacaagagt
tctggtgcattacatgcggcctaatctatttgctgtccatcccgtccatgtacctgctgctcatcttgtactctatcatc
aacctaaacgtcgtctcctggggcacccgcgaggtggtggctaagaagaccaagaaagaactggaggcggagaagaaggc
cgccgaggaggccaaaaagagggtgaagcagaagagcatgctgagcttccttcagagtggaatcggggacaatggcgacg
aggagggctccgtggagttctctctcgctggactcttccgctgcatattctgcacccacggaaaaacatccgacgagaag
cagcagctgacctccatcgccgaatcgctggacacgatcaaacatcgaatggacaccatcgagagtgctgtggatcccca
tggtcaccatgcctcccgacatggcaggaggagaaccacatccagtggctccaaggatcaccacctgctgacctcggtgg
cggagaagtctggcgatgaatcggacgagtcggactcagacacttcggcggaaccaaagcaggagcgagactttctaacc
aatccctactggatcgaggaccccgatgtacgcaagggcgaggtcgatttctctccagcaccgagatccagttctggaa
ggatctcatcgaccagtatctctatcccatcgacaatgatcccgtggagcaggcccgcatagccaaagatctgaaggagc
tgcgcgactcgtcggtgttcgcgttcttcatgatcaatgcactgttcgtgctgatcgtgttcctgctgcagctgaacaag
gacaacatccacgtgaagtggcccttcggagtgcggacgaacatcacctacgacgagtccacccaggaggtgcacatctc
caaggagtaccttcagctggagcccatcggattggtgttcgtattcttctttgctcttattttgataatccagttcacgg
ccatgctgttccatcgatttggaaccatttcgcacatcctggcctcgactgaactcaacttctgcaagaagaagtctgag
gatctgacacaggaccaactaatagataagcatgctgtggagatcgtcaagaacttgcagaggctgcagggaatcgatgg
cgactacgacaatgactccggttccgggccagatcgcatagccaggcggaagaccattcaaaatctggagaaggcgcgac
agccacgtcgccagatcggcaccctggatgtggcgttcaagaagcgcttcctgaaactcaccgccgatgcggagaacaat
ccggccacgcccattctcacccgccgcctgacaatgcgggcggagaccatccgggcactggaggtgcgtaagaattcggt
gatggccgaaaggcggaagtccgcgatgcaaactctgggagcaaagaacgagtacggaatcacaactggagcaccgatta

acaacaatggagctctgccgaatcagcgaagcggacgtgtctcgaatgcaggcatcagcatcaaggatgtattcaatgtg
aacggtggtggtgccgagcagatttacggctcgaatgggggcggcactatcaaccagggttacgagcatgtgatcgacga
ggacggcgacggcaactccctgcggctgaccaccaggaatccccatccccacccgcaccaccaggtctcctggggccaga
acaccaacggcggcggcggtaatggcacaggtcgcctgatggtgagcaagggcgaggaggataacatggcctctctccca
gcgacacatgagttacacatctttggctccatcaacggtgtggactttgacatggtgggtcagggcaccggcaatccaaa
tgatggttatgaggagttaaacctgaagtccaccaagggtgacctccagttctccccctggattctggtccctcatatcg
ggtatggcttccatcagtacctgccctaccctgacgggatgtcgcctttccaggccgccatggtagatggctccggatac
caagtccatcgcacaatgcagtttgaagatggtgcctcccttactgttaactaccgctacacctacgagggaagccacat
caaaggagaggcccaggtgaaggggactggtttccctgctgacggtcctgtgatgaccaactcgctgaccgctgcggact
ggtgcaggtcgaagaagacttaccccaacgacaaaaccatcatcagtacctttaagtggagttacaccactggaaatggc
aagcgctaccggagcactgcgcggaccacctacacctttgccaagccaatggcggctaactatctgaagaaccagccgat
gtacgtgttccgtaagacggagctcaagcactccaagaccgagctcaacttcaaggagtggcaaaaggcctttaccgatg
tgatgggcatggacgagctgtacaagtaatagtctagaggatctttgtgaaggaaccttacttctgtggtgtgacataat
tggacaaactacctacagagatttaaagctctaaggtaaatataaaatttttaagtgtataatgtgttaaactactgatt
ctaattgtttgtgtattttagattccaacctatggaactgatgaatgggagcagtggtggaatgcctttaatgaggaaaa
cctgttttgctcagaagaaatgccatctagtgatgatgaggctactgctgactctcaacattctactcctccaaaaaaga
agagaaaggtagaagaccccaaggactttccttcagaattgctaagttttttgagtcatgctgtgtttagtaatagaact
cttgcttgctttgctatttacaccacaaaggaaaaagctgcactgctatacaagaaaattatggaaaaatatttgatgta
tagtgccttgactagagatcataatcagccataccacatttgtagaggttttacttgctttaaaaaacctcccacacctc
cccctgaacctgaaacataaaatgaatgcaattgttgttgttaacttgtttattgcagcttataatggttacaaataaag
caatagcatcacaaatttcacaaataaagcatttttttcactgcattctagttgtggtttgtccaaactcatcaatgtat
cttatcatgtctggatccactagtgtcgacgatgtaggtcacggtctcgaagccgcggtgcgggtgccagggcgtgccct
tgggctccccgggcgcgtactccacctcacccatctggtccatcatgatgaacgggtcgaggtggcggtagttgatcccg
gcgaacgcgcggcgcaccgggaagccctcgccctcgaaaccgctgggcgcggtggtcacggtgagcacgggacgtgcgac
ggcgtcggcgggtgcggatacgcggggcagcgtcagcgggttctcgacggtcacggcgggcatgtcgacactagttctag
ccagcttttgttccctttagtgagggttaatttcgagcttggcgtaatcatggtcatagctgtttcctgtgtgaaattgt
tatccgctcacaattccacacaacatacgagccggaagcataaagtgtaaagcctggggtgcctaatgagtgagctaact
cacattaattgcgttgcgctcactgcccgctttccagtcgggaaacctgtcgtgccagctgcattaatgaatcggccaac
gcgcggggagaggcggtttgcgtattgggcgctcttccgcttcctcgctcactgactcgctgcgctcggtcgttcggctg
cggcgagcggtatcagctcactcaaaggcggtaatacggttatccacagaatcaggggataacgcaggaaagaacatgtg
agcaaaaggccagcaaaaggccaggaaccgtaaaaaggccgcgttgctggcgtttttccataggctccgcccccctgacg
agcatcacaaaaatcgacgctcaagtcagaggtggcgaaacccgacaggactataaagataccaggcgtttccccctgga
agctccctcgtgcgctctcctgttccgaccctgccgcttaccggatacctgtccgcctttctcccttcgggaagcgtggc
gctttctcatagctcacgctgtaggtatctcagttcggtgtaggtcgttcgctccaagctgggctgtgtgcacgaacccc
ccgttcagcccgaccgctgcgccttatccggtaactatcgtcttgagtccaacccggtaagacacgacttatcgccactg
gcagcagccactggtaacaggattagcagagcgaggtatgtaggcggtgctacagagttcttgaagtggtggcctaacta
cggctacactagaaggacagtatttggtatctgcgctctgctgaagccagttaccttcggaaaaagagttggtagctctt
gatccggcaaacaaaccaccgctggtagcggtggtttttttgtttgcaagcagcagattacgcgcagaaaaaaaggatct
caagaagatcctttgatcttttctacggggtctgacgctcagtggaacgaaaactcacgttaagggattttggtcatgag
attatcaaaaaggatcttcacctagatccttttaaattaaaaatgaagttttaaatcaatctaaagtatatatgagtaaa
cttggtctgacagttaccaatgcttaatcagtgaggcacctatctcagcgatctgtctatttcgttcatccatagttgcc
tgactccccgtcgtgtagataactacgatacgggagggcttaccatctggccccagtgctgcaatgataccgcgagaccc
acgctcaccggctccagatttatcagcaataaaccagccagccggaagggccgagcgcagaagtggtcctgcaactttat
ccgcctccatccagtctattaattgttgccgggaagctagagtaagtagttcgccagttaatagtttgcgcaacgttgtt
gccattgctacaggcatcgtggtgtcacgctcgtcgtttggtatggcttcattcagctccggttcccaacgatcaaggcg
agttacatgatcccccatgttgtgcaaaaaagcggttagctccttcggtcctccgatcgttgtcagaagtaagttggccg
cagtgttatcactcatggttatggcagcactgcataattctcttactgtcatgccatccgtaagatgcttttctgtgact
ggtgagtactcaaccaagtcattctgagaatagtgtatgcggcgaccgagttgctcttgcccggcgtcaatacgggataa
taccgcgccacatagcagaactttaaaagtgctcatcattggaaaacgttcttcggggcgaaaactctcaaggatcttac
cgctgttgagatccagttcgatgtaacccactcgtgcacccaactgatcttcagcatcttttactttcaccagcgtttct
gggtgagcaaaaacaggaaggcaaaatgccgcaaaaaagggaataagggcgacacggaaatgttgaatactcatactctt
cctttttcaatattattgaagcatttatcagggttattgtctcatgagcggatacatatttgaatgtatttagaaaaata
aacaaataggggttccgcgcacatttccccgaaaagtgccac

File S2
> pUAST-attb-KKV-OH 13394 bp

ctgacgcgccctgtagcggcgcattaagcgcggcgggtgtggtggttacgcgcagcgtgaccgctacacttgccagcgcc
ctagcgcccgctcctttcgctttcttcccttcctttctcgccacgttcgccggctttccccgtcaagctctaaatcgggg
gctccctttagggttccgatttagtgctttacggcacctcgaccccaaaaaacttgattagggtgatggttcacgtagtg
ggccatcgccctgatagacggtttttcgccctttgacgttggagtccacgttctttaatagtggactcttgttccaaact
ggaacaacactcaaccctatctcggtctattcttttgatttataagggattttgccgatttcggcctattggttaaaaaa
tgagctgatttaacaaaaatttaacgcgaattttaacaaaatattaacgcttacaatttccattcgccattcaggctgcg
caactgttgggaagggcgatcggtgcgggcctcttcgctattacgccagctggcgaaagggggatgtgctgcaaggcgat
taagttgggtaacgccagggttttcccagtcacgacgttgtaaaacgacggccagtgaattgtaatacgactcactatag
ggcgaattgggtacgtaccgggcccctagtatgtatgtaagttaataaaacccatttttgcggaaagtagataaaaaaaa
catttttttttttttactgcactggatatcattgaacttatctgatcagttttaaatttacttcgatccaagggtatttga
tgtaccaggttctttcgattacctctcactcaaaatgacattccactcaaagtcagcgctgtttgcctccttctctgtcc
acagaaatatcgccgtctctttcgccgctgcgtccgctatctctttcgccaccgtttgtagcgttacgtagcgtcaatgt
ccgccttcagttgcattttgtcagcggtttcgtgacgaagctccaagcggtttacgccatcaattaaacacaaagtgctg
tgccaaaactcctctcgcttcttatttttgtttgtttttttgagtgattggggtggtgattggttttgggtgggtaagcag
gggaaagtgtgaaaaatcccggcaatgggccaagaggatcaggagctattaattcgcggaggcagcaaacacccatctgc
cgagcatctgaacaatgtgagtagtacatgtgcatacatcttaagttcacttgatctataggaactgcgattgcaacatc
aaattgtctgcggcgtgagaactgcgacccacaaaaatcccaaaccgcaattgcacaaacaaatagtgacacgaaacaga
ttattctggtagctgttctcgctatataagacaattttttgagatcatatcatgatcaagacatctaaaggcattcatttt
cgactatattctttttttacaaaaaatataacaaccagatattttaagctgatcctagatgcacaaaaaataaataaaagt
ataaacctacttcgtaggatacttcggggtacttttttgttcggggttagatgagcataacgcttgtagttgatatttgag
atcccctatcattgcagggtgacagcggagcggcttcgcagagctgcattaaccagggcttcgggcaggccaaaaactac
ggcacgctccggccacccagtccgccggaggactccggttcagggagcggccaactagccgagaacctcacctatgcctg
gcacaatatggacatctttggggcggtcaatcagccgggctccggatggcggcagctggtcaaccggacacgcggactat
tctgcaacgagcgacacataccggcgcccaggaaacatttgctcaagaacggtgagtttctattcgcagtcggctgatct
gtgtgaaatcttaataaagggtccaattaccaatttgaaactcagtttgcggcgtggcctatccgggcgaacttttggcc
gtgatgggcagttccggtgccggaaagacgaccctgctgaatgcccttgcctttcgatcgccgcagggcatccaagtatc
gccatccgggatgcgactgctcaatggccaacctgtggacgccaaggagatgcaggccaggtgcgcctatgtccagcagg
atgacctctttatcggctccctaacggccagggaacacctgattttccaggccatggtgcggatgccacgacatctgacc
tatcggcagcgagtggcccgcgtggatcaggtgatccaggagctttcgctcagcaaatgtcagcacacgatcatcggtgt
gcccggcagggtgaaaggtctgtccggcggagaaaggaagcgtctggcattcgcctccgaggcactaaccgatccgccgc
ttctgatctgcgatgagcccacctccggactggactcatttaccgcccacagcgtcgtccaggtgctgaagaagctgtcg
cagaagggcaagaccgtcatcctgaccattcatcagccgtcttccgagctgtttgagctctttgacaagatccttctgat
ggccgagggcagggtagctttcttgggcactcccagcgaagccgtcgacttcttttcctagtgagttcgatgtgtttatt
aagggtatctagcattacattacatctcaactcctatccagcgtgggtgcccagtgtcctaccaactacaatccggcgga
cttttacgtacaggtgttggccgttgtgcccggacgggagatcgagtcccgtgatcggatcgccaagatatgcgacaatt
ttgctattagcaaagtagcccgggatatggagcagttgttggccaccaaaaatttggagaagccactggagcagccggag
aatgggtacacctacaaggccacctggttcatgcagttccgggcggtcctgtggcgatcctggctgtcggtgctcaagga
accactcctcgtaaaagtgcgacttattcagacaacggtgagtggttccagtggaaacaaatgatataacgcttacaatt
cttggaaacaaattcgctagattttagttagaattgcctgattccacacccttcttagtttttttcaatgagatgtatag
tttatagttttgcagaaaataaataaatttcatttaactcgcgaacatgttgaagatatgaatattaatgagatgcgagt
aacattttaatttgcagatggttgccatcttgattggcctcatctttttgggccaacaactcacgcaagtgggcgtgatg
aatatcaacggagccatcttcctcttcctgaccaacatgacctttcaaaacgtctttgccacgataaatgtaagtcttgt
ttagaatacatttgcatattaataatttactaactttctaatgaatcgattcgatttaggtgttcacctcagagctgcca
gtttttatgagggaggcccgaagtcgactttatcgctgtgacacatactttctgggcaaaacgattgccgaattaccgct
ttttctcacagtgccactggtcttcacggcgattgcctatccgatgatcggactgcgggccggagtgctgcacttcttca
actgcctggcgctggtcactctggtggccaatgtgtcaacgtccttcggatatctaatatcctgcgccagctcctcgacc
tcgatggcgctgtctgtgggtccgccggttatcataccattcctgctctttggcggcttcttcttgaactcgggctcggt
gccagtatacctcaaatggttgtcgtacctctcatggttccgttacgccaacgagggtctgctgattaaccaatgggcgg
acgtggagccgggcgaaattagctgcacatcgtcgaacaccacgtgccccagttcgggcaaggtcatcctggagacgctt
aacttctccgccgccgatctgccgctggactacgtgggtctggccattctcatcgtgagcttccgggtgctcgcatatct
ggctctaagacttcgggcccgacgcaaggagtagccgacatatatccgaaataactgcttgtttttttttttttaccatta
ttaccatcgtgtttactgtttattgccccctcaaaaagctaatgtaattatatttgtgccaataaaaacaagatatgacc
tatagaatacaagtatttccccttcgaacatccccacaagtagactttggatttgtcttctaaccaaaagacttacacac
ctgcataccttacatcaaaaactcgtttatcgctacataaaacaccgggatatattttttatatacatacttttcaaatc
gcgcgccctcttcataattcacctccaccacaccacgtttcgtagttgctctttcgctgtctcccacccgctctccgcaa
cacattcacctttgttcgacgaccttggagcgactgtcgttagttccgcgcgattcggttcgctcaaatggttccgagt
ggttcatttcgtctcaatagaaattagtaataaatatttgtatgtacaatttatttgctccaatatatttgtatatattt
ccctcacagctatatttattctaatttaatattatgacttttaaggtaattttttgtgacctgttcggagtgattagcg
ttacaatttgaactgaaagtgacatccagtgtttgttccttgtgtagatgcatctcaaaaaaatggtgggcataatagtg
ttgtttatatatatcaaaaataacaactataataataagaatacatttaatttagaaaatgcttggatttcactggaact

aggctagcataacttcgtataatgtatgctatacgaagttatgctagcggatccaagcttgcatgcctgcaggtcggagt
actgtcctccgagcggagtactgtcctccgagcggagtactgtcctccgagcggagtactgtcctccgagcggagtactg
tcctccgagcggagactctagcgagcgccggagtataaatagaggcgcttcgtctacggagcgacaattcaattcaaaca
agcaaagtgaacacgtcgctaagcgaaagctaagcaaataaacaagcgcagctgaacaagctaaacaatctgcagtaaag
tgcaagttaaagtgaatcaattaaaagtaaccagcaaccaagtaaatcaactgcaactactgaaatctgccaagaagtaa
ttattgaatacaagaagagaactctgaatagggaattgggaattcgttaacagatctgcggccgcggctcgagatgtctg
cgatgcggcatcgcccgatggcccctccgggccaaggaccaggagcgggaaccgctggggagcacgtcgacagcgatgac
aacaactttaccgacgacgagagctcgcccctcacccacgacatttacggcggcagccaacgcaccatacaagagacgaa
gggttgggatgttttccgggatccgccaatcaaaattgagaccggatcgacggcgaaccaggaatgcttagaattaaccg
tgaagatcttgaagatcttcgcctatgtgattacgtttataatagttttaaccggtggcgtcatcgccaagggcacaatg
ctcttcatgacttcgcaggtgcgcaaggataagaagatggagtactgcaacaaagacttgggtcgggacaaaagcttcgt
agtccgacttccggaggaggagcgagtggcctggatctgggctctgctcatagcatatgcccttcccgagatcggagcct
tgatccgatccgcccgcatctgcttcttcaagacctttaaggtgccgaagacggccacttcctcttcgtctggctgatg
gagagcttgagtgccgtgggcatggcgctgctgatgttcgtggttctgccccagatcgacgccatccagggcgccatgct
aaccaactgcctgtgcgtcgtgccgggcatcttcggccttctatcgcgcacctccaaggagggcaagcgctttgtaaagg
tgatcatcgatctggcggcggtggcggcccaggtgacgggcctggtgatctggccgctgctggagaaccgccgtgagctg
tgggtcatcccggtggcctgtgtcatgatctcgtgcggctggtgggagaattacgtgtcgccacagtccccgctcggcct
ggttcgagccctgggtcgcatcaaggaggagatgaagtacactcgctacttctgccacatattcctatccatctggaaga
tcctgcttttcttcaccgtcacactgctcatctactgggcgcaaggcgaggaacccggcaatctgtttgccatgtacgga
gatgcctttggaccccacaagatcatagtctacgaactgccagcgggactgggcggtgtgcttccagataccctggaatc
ggccaacattgacaccgtggatgtggatgccgcctacaatacggtggtgtatgtactgcttctacagattttcggcgcct
acttgtgctacatcttcggaaagttcgcctgcaagatcctcatccagggattcagctacgcgtttcccgtcagtctaact
gttccgctctctgttacgttcctgatcgcagcctgtggaatccggatcgacgatccctgcttcttccatgacaccattcc
ggactatctgttcttcacgagcccctcgaacttccggttcaacaactttgtcaccgagcaaatggcgtgggcctggatcc
tttggctcctcagtcagacctggatagcactgcacatctggactccgaagtgcgagcgtctggccaccaccgagaagttg
tttgtccagcccatgtactcctctctactgatcgatcagtcgatggccctcaaccgaaggcgcgatgaccaggcagatgt
gaaaacagaggatctctcggagatcgaaaaggaaaagggcgatgagtactacgagaccatttctgtgcacacggatcgct
cctcggcacccaacaagccatcgattaagtcctccgacaacatcacacgaatctactcctgcgcgaccatgtggcatgag
acgaaggacgagatgatagagttcttgaagagcatcatgcgaatggacgaggaccagtgtgctcgtcgcgtggctcaaaa
gtatctgagggttctcgatccggactattatgaatttgagacccacatcttcttcgatgacgccttcgaaatctccgatc
acagcgacgatgatatccagtgcaatcgattcgtcaagctcttgatagccaccatggacgaggctgcctccgagatccat
cagaccacgatcagactgcgtccgcccaagaagtatcccactccgtacggaggccgcctggtgtggacccttccagggaa
gaccaaattcattacgcacttaaaggacaaggaccgcattcgtcacaggaagcgttggtctcaggtcatgtacatgtatt
acctgctgggacatcgtctaatggagctgcccatttcggtggatcgcaaggatgcaattgcggagaacacctacctttttg
accctggacggagatattgacttcaagccgaatgcggtgactctactggtggatctgatgaagaagaacaagaacctggg
tgctgcctgcggtcgcattcatcccgtgggatcgggacccatggtgtggtaccaacttttcgaatacgccatcggccatt
ggctgcaaaaggccacggagcacatgattggctgcgtgctctgttcacctggatgcttctccctgttcaggggcaaggcc
ctcatggatgacaatgtgatgaagaaatacacgacacggtcggatgaggctcgtcactacgtgcagtacgatcagggcga
ggatcgttggctgtgcacattgctcctccagaggggataccgcgtggagtactcggctgccagtgatgcgtacacccact
gccccgagggattcaatgagttctacaaccagcggcgcagatgggtgccctcgaccatagccaacatcatggatcttctg
gcggatgcgaagcgcacgatcaagatcaatgacaacatttcgcttctgtacatcttctaccagatgatgttgatgggcgg
caccatcctgggacccggaaccattttccttatgttggtgggtgcattcgtggcagcctttcgcattgacaactggacct
ccttccactacaacatcgtgccgatcctggccttcatgtttatctgcttcacctgtaagtcgaacatccagttgttcgtg
gcccaagtcctttcgacggcatatgccctgatcatgatggcggtgattgtgggtacggccttgcagttgggagaggacgg
aataggctccccttcggccatttcctgatatccatggtgggctcattcttcatagccgcatgtttgcatccacaagagt
tctggtgcattacatgcggcctaatctatttgctgtccatcccgtccatgtacctgctgctcatcttgtactctatcatc
aacctaaacgtcgtctcctggggcacccgcgaggtggtggctaagaagaccaagaaagaactggaggcggagaagaaggc
cgccgaggaggccaaaaagagggtgaagcagaagagcatgctgagcttccttcagagtggaatcggggacaatggcgacg
aggagggctccgtggagttctctctcgctggactcttccgctgcatattctgcacccacggaaaaacatccgacgagaag
cagcagctgacctccatcgccgaatcgctggacacgatcaaacatcgaatggacaccatcgagagtgctgtggatcccca
tggtcaccatgcctcccgacatggcaggaggagaaccacatccagtggctccaaggatcaccacctgctgacctcggtgg
cggagaagtctggcgatgaatcggacgagtcggactcagacacttcggcggaaccaaagcaggagcgagactttctaacc
aatccctactggatcgaggaccccgatgtacgcaagggcgaggtcgattttctctccagcaccgagatccagttctggaa
ggatctcatcgaccagtatctctatcccatcgacaatgatcccgtggagcaggcccgcatagccaaagatctgaaggagc
tgcgcgactcgtcggtgttcgcgttcttcatgatcaatgcactgttcgtgctgatcgtgttcctgctgcagctgaacaag
gacaacatccacgtgaagtggcccttcggagtgcggacgaacatcacctacgacgagtccacccaggaggtgcacatctc
caaggagtaccttcagctggagcccatcggattggtgttcgtattcttctttgctcttattttgataatccagttcacgg
ccatgctgttccatcgatttggaaccatttcgcacatcctggcctcgactgaactcaacttctgcaagaagaagtctgag
gatctgacacaggaccaactaatagataagcatgctgtggagatcgtcaagaacttgcagaggctgcagggaatcgatgg
cgactacgacaatgactccggttccgggccagatcgcatagccaggcggaagaccattcaaaatctggagaaggcgcgac
agccacgtcgccagatcggcaccctggatgtggcgttcaagaagcgcttcctgaaactcaccgccgatgcggagaacaat
ccggccacgcccattctcacccgccgcctgacaatgcgggcggagaccatccgggcactggaggtgcgtaagaattcggt
gatggccgaaaggcggaagtccgcgatgcaaactctgggagcaaagaacgagtacggaatcacaactggagcaccgatta

acaacaatggagctctgccgaatcagcgaagcggacgtgtctcgaatgcaggcatcagcatcaaggatgtattcaatgtg
aacggtggtggtgccgagcagatttacggctcgaatgggggcggcactatcaaccagggttacgagcatgtgatcgacga
ggacggcgacggcaactccctgcggctgaccaccaggaatccccatccccacccgcaccaccaggtctcctggggccaga
acaccaacggcggcggcggtaatggcacaggtcgcctgtccggcttcgccaacgagctgggcccccgcctgatgggcaag
catcaccatcatcaccactaatagtctagaggatctttgtgaaggaaccttacttctgtggtgtgacataattggacaaa
ctacctacagagatttaaagctctaaggtaaatataaaatttttaagtgtataatgtgttaaactactgattctaattgt
ttgtgtattttagattccaacctatggaactgatgaatgggagcagtggtggaatgcctttaatgaggaaaacctgtttt
gctcagaagaaatgccatctagtgatgatgaggctactgctgactctcaacattctactcctccaaaaaagaagagaaag
gtagaagaccccaaggactttccttcagaattgctaagttttttgagtcatgctgtgtttagtaatagaactcttgcttg
ctttgctatttacaccacaaaggaaaaagctgcactgctatacaagaaaattatggaaaaatatttgatgtatagtgcct
tgactagagatcataatcagccataccacatttgtagaggttttacttgctttaaaaaacctcccacacctccccctgaa
cctgaaacataaaatgaatgcaattgttgttgttaacttgtttattgcagcttataatggttacaaataaagcaatagca
tcacaaatttcacaaataaagcatttttttcactgcattctagttgtggtttgtccaaactcatcaatgtatcttatcat
gtctggatccactagtgtcgacgatgtaggtcacggtctcgaagccgcggtgcgggtgccagggcgtgcccttgggctcc
ccgggcgcgtactccacctcacccatctggtccatcatgatgaacgggtcgaggtggcggtagttgatcccggcgaacgc
gcggcgcaccgggaagccctcgccctcgaaaccgctgggcgcggtggtcacggtgagcacgggacgtgcgacggcgtcgg
cgggtgcggatacgcggggcagcgtcagcgggttctcgacggtcacggcgggcatgtcgacactagttctagccagcttt
tgttccctttagtgagggttaatttcgagcttggcgtaatcatggtcatagctgtttcctgtgtgaaattgttatccgct
cacaattccacacaacatacgagccggaagcataaagtgtaaagcctggggtgcctaatgagtgagctaactcacattaa
ttgcgttgcgctcactgcccgctttccagtcgggaaacctgtcgtgccagctgcattaatgaatcggccaacgcgcgggg
agaggcggtttgcgtattgggcgctcttccgcttcctcgctcactgactcgctgcgctcggtcgttcggctgcggcgagc
ggtatcagctcactcaaaggcggtaatacggttatccacagaatcaggggataacgcaggaaagaacatgtgagcaaaag
gccagcaaaaggccaggaaccgtaaaaaggccgcgttgctggcgtttttccataggctccgcccccctgacgagcatcac
aaaaatcgacgctcaagtcagaggtggcgaaacccgacaggactataaagataccaggcgtttccccctggaagctccct
cgtgcgctctcctgttccgaccctgccgcttaccggatacctgtccgcctttctcccttcgggaagcgtggcgctttctc
atagctcacgctgtaggtatctcagttcggtgtaggtcgttcgctccaagctgggctgtgtgcacgaaccccccgttcag
cccgaccgctgcgccttatccggtaactatcgtcttgagtccaacccggtaagacacgacttatcgccactggcagcagc
cactggtaacaggattagcagagcgaggtatgtaggcggtgctacagagttcttgaagtggtggcctaactacggctaca
ctagaaggacagtatttggtatctgcgctctgctgaagccagttaccttcggaaaaagagttggtagctcttgatccggc
aaacaaaccaccgctggtagcggtggtttttttgtttgcaagcagcagattacgcgcagaaaaaaaggatctcaagaaga
tcctttgatcttttctacggggtctgacgctcagtggaacgaaaactcacgttaagggattttggtcatgagattatcaa
aaaggatcttcacctagatccttttaaattaaaaatgaagttttaaatcaatctaaagtatatatgagtaaacttggtct
gacagttaccaatgcttaatcagtgaggcacctatctcagcgatctgtctatttcgttcatccatagttgcctgactccc
cgtcgtgtagataactacgatacgggagggcttaccatctggccccagtgctgcaatgataccgcgagacccacgctcac
cggctccagatttatcagcaataaaccagccagccggaagggccgagcgcagaagtggtcctgcaactttatccgcctcc
atccagtctattaattgttgccgggaagctagagtaagtagttcgccagttaatagtttgcgcaacgttgttgccattgc
tacaggcatcgtggtgtcacgctcgtcgtttggtatggcttcattcagctccggttcccaacgatcaaggcgagttacat
gatcccccatgttgtgcaaaaaagcggttagctccttcggtcctccgatcgttgtcagaagtaagttggccgcagtgtta
tcactcatggttatggcagcactgcataattctcttactgtcatgccatccgtaagatgcttttctgtgactggtgagta
ctcaaccaagtcattctgagaatagtgtatgcggcgaccgagttgctcttgcccggcgtcaatacgggataataccgcgc
cacatagcagaactttaaaagtgctcatcattggaaaacgttcttcggggcgaaaactctcaaggatcttaccgctgttg
agatccagttcgatgtaacccactcgtgcacccaactgatcttcagcatcttttactttcaccagcgtttctgggtgagc
aaaaacaggaaggcaaaatgccgcaaaaaagggaataagggcgacacggaaatgttgaatactcatactcttcctttttc
aatattattgaagcatttatcagggttattgtctcatgagcggatacatatttgaatgtatttagaaaaataaacaaata
ggggttccgcgcacatttccccgaaaagtgccac

File S3
> [Kkv::NG] – protein

MSAMRHRPMAPPGQGPGAGTAGEHVDSDDNNFTDDESSPLTHDIYGGSQRTIQETKGWDVFRDPPIKIETGSTANQECLE
LTVKILKIFAYVITFIIVLTGGVIAKGTMLFMTSQVRKDKKMEYCNKDLGRDKSFVVRLPEEERVAWIWALLIAYALPEI
GALIRSARICFFKTFKVPKTGHFLFVWLMESLSAVGMALLMFVVLPQIDAIQGAMLTNCLCVVPGIFGLLSRTSKEGKRF
VKVIIDLAAVAAQVTGLVIWPLLENRRELWVIPVACVMISCGWWENYVSPQSPLGLVRALGRIKEEMKYTRYFCHIFLSI
WKILLFFTVTLLIYWAQGEEPGNLFAMYGDAFGPHKIIVYELPAGLGGVLPDTLESANIDTVDVDAAYNTVVYVLLLQIF
GAYLCYIFGKFACKILIQGFSYAFPVSLTVPLSVTFLIAACGIRIDDPCFFHDTIPDYLFFTSPSNFRFNNFVTEQMAWA
WILWLLSQTWIALHIWTPKCERLATTEKLFVQPMYSSLLIDQSMALNRRRDDQADVKTEDLSEIEKEKGDEYYETISVHT
DRSSAPNKPSIKSSDNITRIYSCATMWHETKDEMIEFLKSIMRMDEDQCARRVAQKYLRVLDPDYYEFETHIFFDDAFEI
SDHSDDDIQCNRFVKLLIATMDEAASEIHQTTIRLRPPKKYPTPYGGRLVWTLPGKTKFITHLKDKDRIRHRKRWSQVMY
MYYLLGHRLMELPISVDRKDAIAENTYLLTLDGDIDFKPNAVTLLVDLMKKNKNLGAACGRIHPVGSGPMVWYQLFEYAI
GHWLQKATEHMIGCVLCSPGCFSLFRGKALMDDNVMKKYTTRSDEARHYVQYDQGEDRWLCTLLLQRGYRVEYSAASDAY
THCPEGFNEFYNQRRRWVPSTIANIMDLLADAKRTIKINDNISLLYIFYQMMLMGGTILGPGTIFLMLVGAFVAAFRIDN
WTSFHYNIVPILAFMFICFTCKSNIQLFVAQVLSTAYALIMMAVIVGTALQLGEDGIGSPSAIFLISMVGSFFIAACLHP
QEFWCITCGLIYLLSIPSMYLLLILYSIINLNVVSWGTREVVAKKTKKELEAEKKAAEEAKKRVKQKSMLSFLQSGIGDN
GDEEGSVEFSLAGLFRCIFCTHGKTSDEKQQLTSIAESLDTIKHRMDTIESAVDPHGHHASRHGRRRTTSSGSKDHHLLT
SVAEKSGDESDESDSDTSAEPKQERDFLTNPYWIEDPDVRKGEVDFLSSTEIQFWKDLIDQYLYPIDNDPVEQARIAKDL
KELRDSSVFAFFMINALFVLIVFLLQLNKDNIHVKWPFGVRTNITYDESTQEVHISKEYLQLEPIGLVFVFFFALILIIQ
FTAMLFHRFGTISHILASTELNFCKKKSEDLTQDQLIDKHAVEIVKNLQRLQGIDGDYDNDSGSGPDRIARRKTIQNLEK
ARQPRRQIGTLDVAFKKRFLKLTADAENNPATPILTRRLTMRAETIRALEVRKNSVMAERRKSAMQTLGAKNEYGITTGA
PINNNGALPNQRSGRVSNAGISIKDVFNVNGGGAEQIYGSNGGGTINQGYEHVIDEDGDGNSLRLTTRNPHPHPHHQVSW
GQNTNGGGGNGTGRLMVSKGEEDNMASLPATHELHIFGSINGVDFDMVGQGTGNPNDGYEELNLKSTKGDLQFSPWILVP
HIGYGFHQYLPYPDGMSPFQAAMVDGSGYQVHRTMQFEDGASLTVNYRYTYEGSHIKGEAQVKGTGFPADGPVMTNSLTA
ADWCRSKKTYPNDKTIISTFKWSYTTGNGKRYRSTARTTYTFAKPMAANYLKNQPMYVFRKTELKHSKTELNFKEWQKAF
TDVMGMDELYK

File S4

```
> [Kkv::ollas-his] 1635 bp

MSAMRHRPMAPPGQGPGAGTAGEHVDSDDNNFTDDESSPLTHDIYGGSQRTIQETKGWDVFRDPPIKIETGSTANQECLE
LTVKILKIFAYVITFIIVLTGGVIAKGTMLFMTSQVRKDKKMEYCNKDLGRDKSFVVRLPEEERVAWIWALLIAYALPEI
GALIRSARICFFKTFKVPKTGHFLFVWLMESLSAVGMALLMFVVLPQIDAIQGAMLTNCLCVVPGIFGLLSRTSKEGKRF
VKVIIDLAAVAAQVTGLVIWPLLENRRELWVIPVACVMISCGWWENYVSPQSPLGLVRALGRIKEEMKYTRYFCHIFLSI
WKILLFFTVTLLIYWAQGEEPGNLFAMYGDAFGPHKIIVYELPAGLGGVLPDTLESANIDTVDVDAAYNTVVYVLLLQIF
GAYLCYIFGKFACKILIQGFSYAFPVSLTVPLSVTFLIAACGIRIDDPCFFHDTIPDYLFFTSPSNFRFNNFVTEQMAWA
WILWLLSQTWIALHIWTPKCERLATTEKLFVQPMYSSLLIDQSMALNRRRDDQADVKTEDLSEIEKEKGDEYYETISVHT
DRSSAPNKPSIKSSDNITRIYSCATMWHETKDEMIEFLKSIMRMDEDQCARRVAQKYLRVLDPDYYEFETHIFFDDAFEI
SDHSDDDIQCNRFVKLLIATMDEAASEIHQTTIRLRPPKKYPTPYGGRLVWTLPGKTKFITHLKDKDRIRHRKRWSQVMY
MYYLLGHRLMELPISVDRKDAIAENTYLLTLDGDIDFKPNAVTLLVDLMKKNKNLGAACGRIHPVGSGPMVWYQLFEYAI
GHWLQKATEHMIGCVLCSPGCFSLFRGKALMDDNVMKKYTTRSDEARHYVQYDQGEDRWLCTLLLQRGYRVEYSAASDAY
THCPEGFNEFYNQRRRWVPSTIANIMDLLADAKRTIKINDNISLLYIFYQMMLMGGTILGPGTIFLMLVGAFVAAFRIDN
WTSFHYNIVPILAFMFICFTCKSNIQLFVAQVLSTAYALIMMAVIVGTALQLGEDGIGSPSAIFLISMVGSFFIAACLHP
QEFWCITCGLIYLLSIPSMYLLLILYSIINLNVVSWGTREVVAKKTKKELEAEKKAAEEAKKRVKQKSMLSFLQSGIGDN
GDEEGSVEFSLAGLFRCIFCTHGKTSDEKQQLTSIAESLDTIKHRMDTIESAVDPHGHHASRHGRRRTTSSGSKDHHLLT
SVAEKSGDESDESDSDTSAEPKQERDFLTNPYWIEDPDVRKGEVDFLSSTEIQFWKDLIDQYLYPIDNDPVEQARIAKDL
KELRDSSVFAFFMINALFVLIVFLLQLNKDNIHVKWPFGVRTNITYDESTQEVHISKEYLQLEPIGLVFVFFFALILIIQ
FTAMLFHRFGTISHILASTELNFCKKKSEDLTQDQLIDKHAVEIVKNLQRLQGIDGDYDNDSGSGPDRIARRKTIQNLEK
ARQPRRQIGTLDVAFKKRFLKLTADAENNPATPILTRRLTMRAETIRALEVRKNSVMAERRKSAMQTLGAKNEYGITTGA
PINNNGALPNQRSGRVSNAGISIKDVFNVNGGGAEQIYGSNGGGTINQGYEHVIDEDGDGNSLRLTTRNPHPHPHHQVSW
GQNTNGGGGNGTGRLSGFANELGPRLMGKHHHHHH
```

File S5
pUAST-kkv-mos::NG

aGctgacgcgccctgtagcggcgcattaagcgcggcgggtgtggtggttacgcgcagcgtgaccgctacacttgccagcg
ccctagcgcccgctcctttcgctttcttcccttcctttctcgccacgttcgccggctttccccgtcaagctctaaatcgg
gggctccctttagggttccgatttagtgctttacggcacctcgaccccaaaaaacttgattagggtgatggttcacgtag
tgggccatcgccctgatagacggtttttcgccctttgacgttggagtccacgttctttaatagtggactcttgttccaaa
ctggaacaacactcaaccctatctcggtctattcttttgatttataagggattttgccgatttcggcctattggttaaaa
aatgagctgatttaacaaaaatttaacgcgaattttaacaaaatattaacgcttacaatttccattcgccattcaggctg
cgcaactgttgggaagggcgatcggtgcgggcctcttcgctattacgccagctggcgaaagggggatgtgctgcaaggcg
attaagttgggtaacgccagggttttcccagtcacgacgttgtaaaacgacggccagtgaattgtaatacgactcactat
agggcgaattgggtacgtaccgggcccctagtatgtatgtaagttaataaaacccattttgcggaaagtagataaaaaa
aacatttttttttttttactgcactggatatcattgaacttatctgatcagttttaaatttacttcgatccaagggtattt
gatgtaccaggttctttcgattacctctcactcaaaatgacattccactcaaagtcagcgctgtttgcctccttctctgt
ccacagaaatatcgccgtctctttcgccgctgcgtccgctatctctttcgccaccgtttgtagcgttacgtagcgtcaat
gtccgccttcagttgcattttgtcagcggtttcgtgacgaagctccaagcggtttacgccatcaattaaacacaaagtgc
tgtgccaaaactcctctcgcttcttatttttgtttgtttttttgagtgattggggtggtgattggttttgggtgggtaagc
aggggaaagtgtgaaaaatcccggcaatgggccaagaggatcaggagctattaattcgcggaggcagcaaacacccatct
gccgagcatctgaacaatgtgagtagtacatgtgcatacatcttaagttcacttgatctataggaactgcgattgcaaca
tcaaattgtctgcggcgtgagaactgcgacccacaaaaatcccaaaccgcaattgcacaaacaaatagtgacacgaaaca
gattattctggtagctgttctcgctatataagacaattttgagatcatatcatgatcaagacatctaaaggcattcatt
ttcgactatattctttttttacaaaaaatataacaaccagatattttaagctgatcctagatgcacaaaaaataaataaaa
gtataaacctacttcgtaggatacttcggggtactttttgttcggggttagatgagcataacgcttgtagttgatatttg
agatcccctatcattgcagggtgacagcggagcggcttcgcagagctgcattaaccagggcttcgggcaggccaaaaact
acggcacgctccggccacccagtccgccggaggactccggttcagggagcggccaactagccgagaacctcacctatgcc
tggcacaatatggacatctttggggcggtcaatcagccgggctccggatggcggcagctggtcaaccggacacgcggact
attctgcaacgagcgacacataccggcgcccaggaaacatttgctcaagaacggtgagtttctattcgcagtcggctgat
ctgtgtgaaatcttaataaagggtccaattaccaatttgaaactcagtttgcggcgtggcctatccgggcgaacttttgg
ccgtgatgggcagttccggtgccggaaagacgaccctgctgaatgcccttgcctttcgatcgccgcagggcatccaagta
tcgccatccgggatgcgactgctcaatggccaacctgtggacgccaaggagatgcaggccaggtgcgcctatgtccagca
ggatgacctctttatcggctccctaacggccagggaacacctgattttccaggccatggtgcggatgccacgacatctga
cctatcggcagcgagtggcccgcgtggatcaggtgatccaggagctttcgctcagcaaatgtcagcacacgatcatcggt
gtgcccggcagggtgaaaggtctgtccggcggagaaaggaagcgtctggcattcgcctccgaggcactaaccgatccgcc
gcttctgatctgcgatgagcccacctccggactggactcatttaccgcccacagcgtcgtccaggtgctgaagaagctgt
cgcagaagggcaagaccgtcatcctgaccattcatcagccgtcttccgagctgtttgagctctttgacaagatccttctg
atggccgagggcagggtagctttcttgggcactcccagcgaagccgtcgacttcttttcctagtgagttcgatgtgttta
ttaagggtatctagcattacattacatctcaactcctatccagcgtgggtgcccagtgtcctaccaactacaatccggcg
gacttttacgtacaggtgttggccgttgtgcccggacgggagatcgagtcccgtgatcggatcgccaagatatgcgacaa
ttttgctattagcaaagtagcccgggatatggagcagttgttggccaccaaaaatttggagaagccactggagcagccgg
agaatgggtacacctacaaggccacctggttcatgcagttccgggcggtcctgtggcgatcctggctgtcggtgctcaag
gaaccactcctcgtaaaagtgcgacttattcagacaacggtgagtggttccagtggaaacaaatgatataacgcttacaa
ttcttggaaacaaattcgctagattttagttagaattgcctgattccacacccttcttagtttttttcaatgagatgtat
agtttatagttttgcagaaaataaataaatttcatttaactcgcgaacatgttgaagatatgaatattaatgagatgcga
gtaacattttaatttgcagatggttgccatcttgattggcctcatctttttgggccaacaactcacgcaagtgggcgtga
tgaatatcaacggagccatcttcctcttcctgaccaacatgacctttcaaaacgtctttgccacgataaatgtaagtctt
gtttagaatacatttgcatattaataatttactaactttctaatgaatcgattcgatttaggtgttcacctcagagctgc
cagtttttatgagggaggcccgaagtcgactttatcgctgtgacacatactttctgggcaaaacgattgccgaattaccg
ctttttctcacagtgccactggtcttcacggcgattgcctatccgatgatcggactgcgggccggagtgctgcacttctt
caactgcctggcgctggtcactctggtggccaatgtgtcaacgtccttcggatatctaatatcctgcgccagctcctcga
cctcgatggcgctgtctgtgggtccgccggttatcataccattcctgctctttggcggcttcttcttgaactcgggctcg
gtgccagtatacctcaaatggttgtcgtacctctcatggttccgttacgccaacgagggtctgctgattaaccaatgggc
ggacgtggagccgggcgaaattagctgcacatcgtcgaacaccacgtgccccagttcgggcaaggtcatcctggagacgc
ttaacttctccgccgccgatctgccgctggactacgtgggtctggccattctcatcgtgagcttccgggtgctcgcatat
ctggctctaagacttcgggcccgacgcaaggagtagccgacatatatccgaaataactgcttgtttttttttttttaccat
tattaccatcgtgtttactgtttattgcccctcaaaaagctaatgtaattatatttgtgccaataaaaacaagatatga
cctatagaatacaagtatttccccttcgaacatccccacaagtagactttggatttgtcttctaaccaaaagacttacac
acctgcataccttacatcaaaaactcgtttatcgctacataaaacaccgggatatattttttatatacatacttttcaaa
tcgcgcgccctcttcataattcacctccaccacaccacgtttcgtagttgctctttcgctgtctcccacccgctctccgc
aacacattcaccttttgttcgacgaccttggagcgactgtcgttagttccgcgcgattcggttcgctcaaatggttccga
gtggttcatttcgtctcaatagaaattagtaataaatatttgtatgtacaatttatttgctccaatatatttgtatatat
ttccctcacagctatatttattctaatttaatattatgacttttaaggtaatttttgtgacctgttcggagtgattag
cgttacaatttgaactgaaagtgacatccagtgtttgttccttgtgtagatgcatctcaaaaaaatggtgggcataatag
tgttgtttatatatatcaaaaataacaactataataataagaatacatttaatttagaaaatgcttggatttcactggaa

ctaggctagcataacttcgtataatgtatgctatacgaagttatgctagcggatccaagcttgcatgcctgcaggtcgga
gtactgtcctccgagcggagtactgtcctccgagcggagtactgtcctccgagcggagtactgtcctccgagcggagtac
tgtcctccgagcggagactctagcgagcgccggagtataaatagaggcgcttcgtctacggagcgacaattcaattcaaa
caagcaaagtgaacacgtcgctaagcgaaagctaagcaaataaacaagcgcagctgaacaagctaaacaatctgcagtaa
agtgcaagttaaagtgaatcaattaaaagtaaccagcaaccaagtaaatcaactgcaactactgaaatctgccaagaagt
aattattgaatacaagaagagaactctgaatagggaattgggaattcgttaacagatctgcggccgcggctcgagatgtc
tgcgatgcggcatcgcccgatggcccctccgggccaaggaccaggagcgggaaccgctggggagcacgtcgacagcgatg
acaacaactttaccgacgacgagagctcgcccctcacccacgacatttacggcggcagccaacgcaccatacaagagacg
aagggttgggatgttttccgggatccgccaatcaaaattgagaccggatcgacggcgaaccaggaatgcttagaattaac
cgtgaagatcttgaagatcttcgcctatgtgattacgtttataatagttttaaccggtggcgtcatcgccaagggcacaa
tgctcttcatgacttcgcaggtgcgcaaggataagaagatggagtactgcaacaaagacttgggtcgggacaaaagcttc
gtagtccgacttccggaggaggagcgagtggcctggatctgggctctgctcatagcatatgcccttcccgagatcggagc
cttgatccgatccgcccgcatctgcttcttcaagacctttaaggtgccgaagacgggccacttcctcttcgtctggctga
tggagagcttgagtgccgtgggcatggcgctgctgatgttcgtggttctgccccagatcgacgccatccagggcgccatg
ctaaccaactgcctgtgcgtcgtgccgggcatcttcggccttctatcgcgcacctccaaggagggcaagcgctttgtaaa
ggtgatcatcgatctggcggcggtggcggcccaggtgacgggcctggtgatctggccgctgctggagaaccgccgtgagc
tgtgggtcatcccggtggcctgtgtcatgatctcgtgcggctggtgggagaattacgtgtcgccacagtccccgctcggc
ctggttcgagccctgggtcgcatcaaggaggagatgaagtacactcgctacttctgccacatattcctatccatctggaa
gatcctgcttttcttcaccgtcacactgctcatctactgggcgcaaggcgaggaacccggcaatctgtttgccatgtacg
gagatgcctttggaccccacaagatcatagtctacgaactgccagcgggactgggcggtgtgcttccagataccctggaa
tcggccaacattgacaccgtggatgtggatgccgcctacaatacggtggtgtatgtactgcttctacagattttcggcgc
ctacttgtgctacatcttcggaaagttcgcctgcaagatcctcatccagggattcagctacgcgtttcccgtcagtctaa
ctgttccgctctctgttacgttcctgatcgcagcctgtggaatccggatcgacgatccctgcttcttccatgacaccatt
ccggactatctgttcttcacgagcccctcgaacttccggttcaacaactttgtcaccgagcaaatggcgtgggcctggat
cctttggctcctcagtcagacctggatagcactgcacatctggactccgaagtgcgagcgtctggccaccaccgagaagt
tgtttgtccagcccatgtactcctctctactgatcgatcagtcgatggccctcaaccgaaggcgcgatgaccaggcagat
gtgaaaacagaggatctctcggagatcgaaaaggaaaagggcgatgagtactacgagaccatttctgtgcacacggatcg
ctcctcggcacccaacaagccatcgattaagtcctccgacaacatcacacgaatctactcctgcgcgaccatgtggcatg
agacgaaggacgagatgatagagttcttgaagagcatcatgcgaatggacgaggaccagtgtgctcgtcgcgtggctcaa
aagtatctgagggttctcgatccggactattatgaatttgagacccacatcttcttcgatgacgccttcgaaatctccga
tcacagcgacgatgatatccagtgcaatcgattcgtcaagctcttgatagccaccatggacgaggctgcctccgagatcc
atcagaccacgatcagactgcgtccgcccaagaagtatcccactccgtacggaggccgcctggtgtggacccttccaggg
aagaccaaattcattacgcacttaaaggacaaggaccgcattcgtcacaggaagcgttggtctcaggtcatgtacatgta
ttacctgctgggacatcgtctaatggagctgcccatttcggtggatcgcaaggatgtaatggcggagaacacctaccttt
tgaccctggacggagatattgacttcaacccgagtgcggtgactctactggtggatctgatgaagaagaacaagaacctg
ggtgctgcctgcggtcgcattcatcccatcggatcgggacccatggtgtggtaccaaaaattcgaatacgccatcggcca
ttggctgcaaaaggccacggagcacatgattggctgcgtgctctgttcacctggatgcttctccctgttcaggggcaagg
gcctcatggatgacaatgtgatgaggaaatacacgacacggtcggatgaggctcgtcactacgtgcagtacgatcagggc
gaggatcgttggctgtgcacattgctcctccagaggggataccgcgtggagtactcggctgccagtgatgcgtacaccca
ctgccccgagggattcaatgagttctacaaccagcggcgcagatgggtgccctcgaccatagccaacatcatggatcttc
tggcggatgcgaagcgcacgatcaagatcaatgacaacatttcgcttctgtacatcttctaccagatgatgttgatgggc
ggcaccatcctgggacccggaaccattttccttatgttggtgggtgcattcgtggcagcctttcgcattgacaactggac
ctccttccactacaacatcgtgccgatcctggccttcatgtttatctgcttcacctgtaagtcgaacatccagttgttcg
tggcccaagtcctttcgacggcatatgccctgatcatgatggcggtgattgtgggtacggccttgcagttgggagaggac
ggaataggctccccttcggccattttcctgatatccatggtgggctcattcttcatagccgcatgtttgcatccacaaga
gttctggtgcattacatgcggcctaatctatttgctgtccatcccgtccatgtacctgctgctcatcttgtactctatca
tcaacctaaacgtcgtctcctggggcacccgcgaggtggtggctaagaagaccaagaaagaactggaggcggagaagaag
gccgccgaggaggccaaaaagagggtgaagcagaagagcatgctgagcttccttcagagtggaatcggggacaatggcga
cgaggagggctccgtggagttctctctcgctggactcttccgctgcatattctgcacccacggaaaaacatccgacgaga
agcagcagctgacctccatcgccgaatcgctggacacgatcaaacatcgaatggacaccatcgagagtgctgtggatccc
catggtcaccatgcctcccgacatggcaggaggagaaccacatccagtggctccaaggatcaccacctgctgacctcggt
ggcggagaagtctggcgatgaatcggacgagtcggactcagacacttcggcggaaccaaagcaggagcgagactttctaa
ccaatccctactggatcgaggaccccgatgtacgcaagggcgaggtcgatttctctccagcaccgagatccagttctgg
aaggatctcatcgaccagtatctctatcccatcgacaatgatcccgtggagcaggcccgcatagccaaagatctgaagga
gctgcgcgactcgtcggtgttcgcgttcttcatgatcaatgcactgttcgtgctgatcgtgttcctgctgcagctgaaca
aggacaacatccacgtgaagtggcccttcggagtgcggacgaacatcacctacgacgagtccacccaggaggtgcacatc
tccaaggagtaccttcagctggagcccatcggattggtgttcgtattcttctttgctcttattttgataatccagttcac
ggccatgctgttccatcgatttggaaccatttcgcacatcctggcctcgactgaactcaacttctgcaagaagaagtctg
aggatctgacacaggaccaactaatagataagcatgctgtggagatcgtcaagaacttgcagaggctgcagggaatcgat
ggcgactacgacaatgactccggttccgggccagatcgcatagccaggcggaagaccattcaaaatctggagaaggcgcg
acagccacgtcgccagatcggcaccctggatgtggcgttcaagaagcgcttcctgaaactcaccgccgatgcggagaaca
atccggccacgcccattctcacccgccgcctgacaatgcgggcggagaccatccgggcactggaggtgcgtaagaattcg
gtgatggccgaaaggcggaagtccgcgatgcaaactctgggagcaaagaacgagtacggaatcacaactggagcaccgat

taacaacaatggagctctgccgaatcagcgaagcggacgtgtctcgaatgcaggcatcagcatcaaggatgtattcaatg
tgaacggtggtggtgccgagcagatttacggctcgaatgggggcggcactatcaaccagggttacgagcatgtgatcgac
gaggacggcgacggcaactccctgcggctgaccaccaggaatccccatccccacccgcaccaccaggtctcctggggcca
gaacaccaacggcggcggcggtaatggcacaggtcgcctgatggtgagcaagggcgaggaggataacatggcctctctcc
cagcgacacatgagttacacatctttggctccatcaacggtgtggactttgacatggtgggtcagggcaccggcaatcca
aatgatggttatgaggagttaaacctgaagtccaccaagggtgacctccagttctcccccctggattctggtccctcatat
cgggtatggcttccatcagtacctgccctaccctgacgggatgtcgcctttccaggccgccatggtagatggctccggat
accaagtccatcgcacaatgcagtttgaagatggtgcctcccttactgttaactaccgctacacctacgagggaagccac
atcaaaggagaggcccaggtgaaggggactggtttccctgctgacggtcctgtgatgaccaactcgctgaccgctgcgga
ctggtgcaggtcgaagaagacttaccccaacgacaaaaccatcatcagtacctttaagtggagttacaccactggaaatg
gcaagcgctaccggagcactgcgcggaccacctacacctttgccaagccaatggcggctaactatctgaagaaccagccg
atgtacgtgttccgtaagacggagctcaagcactccaagaccgagctcaacttcaaggagtggcaaaaggcctttaccga
tgtgatgggcatggacgagctgtacaagtaatagtctagaggatctttgtgaaggaaccttacttctgtggtgtgacata
attggacaaactacctacagagatttaaagctctaaggtaaatataaaatttttaagtgtataatgtgttaaactactga
ttctaattgtttgtgtattttagattccaacctatggaactgatgaatgggagcagtggtggaatgcctttaatgaggaa
aacctgttttgctcagaagaaatgccatctagtgatgatgaggctactgctgactctcaacattctactcctccaaaaaa
gaagagaaaggtagaagaccccaaggactttccttcagaattgctaagtttttgagtcatgctgtgtttagtaatagaa
ctcttgcttgctttgctatttacaccacaaaggaaaaagctgcactgctatacaagaaaattatggaaaaatatttgatg
tatagtgccttgactagagatcataatcagccataccacatttgtagaggttttacttgctttaaaaaacctcccacacc
tccccctgaacctgaaacataaaatgaatgcaattgttgttgttaacttgtttattgcagcttataatggttacaaataa
agcaatagcatcacaaatttcacaaataaagcatttttttcactgcattctagttgtggtttgtccaaactcatcaatgt
atcttatcatgtctggatccactagtgtcgacgatgtaggtcacggtctcgaagccgcggtgcgggtgccagggcgtgcc
cttgggctccccgggcgcgtactccacctcacccatctggtccatcatgatgaacgggtcgaggtggcggtagttgatcc
cggcgaacgcgcggcgcaccgggaagccctcgccctcgaaaccgctgggcgcggtggtcacggtgagcacgggacgtgcg
acggcgtcggcgggtgcggatacgcggggcagcgtcagcgggttctcgacggtcacggcgggcatgtcgacactagttct
agccagcttttgttccctttagtgagggttaatttcgagcttggcgtaatcatggtcatagctgtttcctgtgtgaaatt
gttatccgctcacaattccacacaacatacgagccggaagcataaagtgtaaagcctggggtgcctaatgagtgagctaa
ctcacattaattgcgttgcgctcactgcccgctttccagtcgggaaacctgtcgtgccagctgcattaatgaatcggcca
acgcgcggggagaggcggtttgcgtattgggcgctcttccgcttcctcgctcactgactcgctgcgctcggtcgttcggc
tgcggcgagcggtatcagctcactcaaaggcggtaatacggttatccacagaatcaggggataacgcaggaaagaacatg
tgagcaaaaggccagcaaaaggccaggaaccgtaaaaaggccgcgttgctggcgtttttccataggctccgcccccctga
cgagcatcacaaaaatcgacgctcaagtcagaggtggcgaaacccgacaggactataaagataccaggcgtttccccctg
gaagctccctcgtgcgctctcctgttccgaccctgccgcttaccggatacctgtccgcctttctcccttcgggaagcgtg
gcgctttctcatagctcacgctgtaggtatctcagttcggtgtaggtcgttcgctccaagctgggctgtgtgcacgaacc
ccccgttcagcccgaccgctgcgccttatccggtaactatcgtcttgagtccaacccggtaagacacgacttatcgccac
tggcagcagccactggtaacaggattagcagagcgaggtatgtaggcggtgctacagagttcttgaagtggtggcctaac
tacggctacactagaaggacagtatttggtatctgcgctctgctgaagccagttaccttcggaaaaagagttggtagctc
ttgatccggcaaacaaaccaccgctggtagcggtggtttttttgtttgcaagcagcagattacgcgcagaaaaaaaggat
ctcaagaagatcctttgatcttttctacggggtctgacgctcagtggaacgaaaactcacgttaagggattttggtcatg
agattatcaaaaaggatcttcacctagatccttttaaattaaaaatgaagttttaaatcaatctaaagtatatatgagta
aacttggtctgacagttaccaatgcttaatcagtgaggcacctatctcagcgatctgtctatttcgttcatccatagttg
cctgactccccgtcgtgtagataactacgatacgggagggcttaccatctggccccagtgctgcaatgataccgcgagac
ccacgctcaccggctccagatttatcagcaataaaccagccagccggaagggccgagcgcagaagtggtcctgcaacttt
atccgcctccatccagtctattaattgttgccgggaagctagagtaagtagttcgccagttaatagtttgcgcaacgttg
ttgccattgctacaggcatcgtggtgtcacgctcgtcgtttggtatggcttcattcagctccggttcccaacgatcaagg
cgagttacatgatcccccatgttgtgcaaaaaagcggttagctccttcggtcctccgatcgttgtcagaagtaagttggc
cgcagtgttatcactcatggttatggcagcactgcataattctcttactgtcatgccatccgtaagatgcttttctgtga
ctggtgagtactcaaccaagtcattctgagaatagtgtatgcggcgaccgagttgctcttgcccggcgtcaatacgggat
aataccgcgccacatagcagaactttaaaagtgctcatcattggaaaacgttcttcggggcgaaaactctcaaggatctt
accgctgttgagatccagttcgatgtaacccactcgtgcacccaactgatcttcagcatcttttactttcaccagcgttt
ctgggtgagcaaaaacaggaaggcaaaatgccgcaaaaaagggaataagggcgacacggaaatgttgaatactcatactc
ttcctttttcaatattattgaagcatttatcagggttattgtctcatgagcggatacatatttgaatgtatttagaaaaa
taaacaaataggggttccgcgcacatttccccgaaaagtgccacCCCCGAGCCGTAtAACccCcaacGAtgCacGACtat
taAtGACGatTaGgGgCcggAGaCTtACTCtAgcCTaAAcGtCTAgTGCTCgccGtTGttGaCttGCaGtCctgGCttgT
GACaAgCCaTtAtAATgacGTcTAaGaTGCcctGCtaAGGttAGaaacTActTtCtCAagTggAAGGTCGgCCGgCcaAT
gggCtCcTCaAGtCCTTCctaTggGCtaCCTTAACTgtcTgTGcGctACgATAAccGtcCGtCGcgCTgGCaGaTCAact
GgGAgttTcgatTAcAgcAAgAcCACgGcgTAgcTaaAAGcGtgTTTGGtcgtTCctGAgCAcatAggGtaGGCCacAAA
gaCacgcCTGCTCCaAaTgtcgATgCTacatGGaGTgAGaTTcagccAgCcAaagattTagcTGcgcaTTcCAGTGATct
tcTAGatgCaTcaTGgataGcCCagatgGGACGTCCCGGTACCGGGCACAACGTTGTGCCACGTGAACAACGTGCGTGCG
CACGCACGTTGTTGCACAACGTTGTGCAACGCACGTGCGTTGACGTCGAATTGGCCAATTCCACGTGAACACGTGCGCAC
GTGTTGCATGCGAACTCCGGAGTTCCAGCTGCCAGGCTGCAGCCTGGCCAGGCCTGGCCGGGCCCGGGCCCGGCCGGGCC
CGGAACGTGCACGTTCCGGCCGGACGTACGTACGTCCGGCCGGAACGGCGTGCGCACGCCGTTGCACGGCGTTAACGCCG
TGCCGATGCGCATCGCGATCTGCGCAGATCGTGATCATGATCTCATGAGATCATGATCGATCAAACCTGCGTGCGCACGC

AGGTTGCACCTGCGTTAACGCAGGTGCGCCGGCAACGTGCCGGCACGTTAACTGGGGTGCGCACCCCAGTTGCACTGGGG
TTAACCCCAGTGCGACGTCGAGCTCCCGGTCTCGAGACCGGGATATCGATCATCGATGATCGATCGATCCCGGCGCCGGA
CCTCCGGAGGTCCGGCCGGCCGGAACGTGCAGCTGCACGTTGATCCGGATCGATCATGATCCCGGCCGGAACTGGGTGCG
CACCCAGTTGCACTGGGTTAACCCAGTGCAACAACCGGTGCGTGCGCACGCACCGGTTGTTGCACAACCGGTTGTGCAAC
GCACCGGTGCGTTGCATGCGCACGTTCGAACGTGCCCATGGCCAGGCCTGGCCTGGCCAGGCAAGTGGCCACTTGTTTAA
ACGTGCGCACGTTTAAAGCACGTTTAAACGTGCCCAGGCCTGGCCTAGGCCAGGCCGGCCTGGCCGGGCCTGGCCAGGCC
CGGGATCGATCGATCTCGAGATCGAGATCGATCTCGATCGATCGTAACGTGCGCACGTTACGCACGTTAACGTGCGATCG
ATCGCGCTAACTATAACGGTCCTAAGGTAGCGAATTCGCTACCTTAGGACCGTTATAGTTATAGGGATAACAGGGTAATA
TTACCCTGTTATCCCTAGTTAACGTGCGCACGTTAACAACAACGCGTGCGTGCGCACGCACGCGTTGTTGCACAACGCGT
TGTGCAACGCACGCGTGCGTTTCCAACGTGCGCACGTTGGACATGAACAGTGCGCACTGTTGTTTAAACGTGCGCACGTT
TAAACCCGGCGCCGGCGCCGGCCGGCCGGCCGGGATCGCGATCAACAACATGTGCGTGCGCACGCACATGTTGTTGCACA
ACATGTTGTGCAACGCACATGTGCGTTGAATTCGACGTCCCATGGCCAGGTGGCCACCTGGTGGCAAACAGCTATTATGG
GTATTATGGGTACCCATAATACCCATAATAGCTGTTTGCCAATCTATGTCGGGTGCGGAGAAAGAGGTAATGAAATGGCA
TGCCATTTCATTACCTCTTTCTCCGCACCCGACATAGATAACGCACGTGCGTTGAACCTCGAGGTTCCCAGGCCGGCCTG
GCCGGGCCTGGCCAGGCCCGGCCAGGCCTGGTTATAACGTGCGCACGTTATAAGCACGTTATAACGTGCGGCCGGCCAAC
ACTAGTGCGCACTAGTGTTAACTAGTGTGCGCACACTAGTTAACAACTAGTGCGTGCGCACGCACTAGTTGTTGCACAAC
TAGTTGTGCAACGCACTAGTGCGTTAACAACGTGCGTGCGCACGCACGTTGTTTTAACGTGCGCACGTTAACACTCCGGA
GTGCACGATCCGGATCGTGGATCTAGATCCCATGGATAACGGTCCTAAGGTAGCGAATTCGCTACCTTAGGACCGTTATG
CACCCAGTTAACTGGGTGCTAACTATGACTCTCTTAAGGTAGCCAAATATTTGGCTACCTTAAGAGAGTCATAGTTACCG
GTACCGGTACCGGGAAGTACGTACTTCCGAGTTCATAGCTGGGCCCAACGTTGGGCCCAGCTATGAACTCGAAGCTTGAA
CTACGTAGTTCGAGATCGAGGTCGGCGCGCCGCGATCGCCCGGCCGGCCGGCCGGGACGTCCCGGGATCGATCGATCCGG
ATCCGGATCGATCATGATCATGATCCCTGGCCGGCCAGGCCGGCCGGCCGGCCAGGCCTAGGCCTGGGGCCGGCCTGCGC
AGATCGATCGATCCCTGGCCATGGCCAGGGTTTAAACGCGGCCGCGATCGCGATCGCGATCTTAATTAACTCGAGCCTGC
AGGCCCGGGATTTAAATGATCTAGATCTAGATCCGCGCGCGCGCCGGCGCGTCGACGGCCCGGGCCCTCGAGGCACCTGC
GCAGGTGCCTCAGCGCTGAGGGCTAGCCCTAGCGCTAGGGATCGATATCGATCGGTACCCCTAGGCCCGGGGCCCCGGGC
GGCCGCCTGGAGCTCCAGGGGCCGGTGATCGATCACCGATCGATCGATCGCTCTTCGAAGAGCGAAGATCGATCTTCCCC
GGGTCCGGATCCGGAGGCCACCGGTGGGCCCGACGTCGTACGGTACCAACGTTCTGAAGCTTCAGAGCGCTCTTAAGACG
TACCGGTCACGTCGACGTGGGCCGTGCACGGGCCCAATTATTAATCCGGCCTAGGGGATCCGGCCGGCCCACGAGCTCGT
GGAAGACGTCTTCGTATCCGGATACTGATCAACTAGTACTGGGCCCAGTCTAGATGCATACCTGCGCAGGTAGATCTGGC
CGCTAGCCTGGAGCTCCAGCTTGAGCTCAAGGGTCTCGAGACCACGTGCGCCCGGCCGGGCAATGCATTGCGAGGAGCTC
CTCCCCAGCGCTGGGGTGCAGCTGCACCGCGCGTACGGAATGCGCATTCCGTCTCGAGACGTTCGAAGGCCTGTACAGCG
CTCGCGAATCGATTCCGGATCATGAGGCCCCGCTCGAGCGGCCGGGCGCGCGTATACGATCCCGGGCGATGCATCGCGCA
GTGCACTGCGCGCGGCCCCGCGGTTTAAACGGCCGCTCTTCGAAGAGGGCGGATCCGCCGAGCTCGATATCTACGTACCG
GAGGCCTCTGAGCTCAGCACGTGCAGCAGCTGCTGGAATTCGGCGCCTGCGCAGTACAAGCTTGTTAACTCGAGTACCAA
TTGTGGCCAACGCGTTCCACGTGGACGCGGCCGGCCCATGGCATATGCATGGCATGCCTCGAGACATGTTTATAACTGCA
GCGATCGCAGCTGGTCGACAGTACTCCCGGGCTAGAATATTGATCGATCGATCGTACTACCATGGTATCTAGAGCCGAGC
TCGGCGACCGACACCCATCGGTCTGGGTGGGATCGATCCGTCTCGAGACGCGCGGCCGCAGCGCTGCCCATCGATGGACG
GCGCCGTCCGGCCGGCCGGGGATGCATCCCTCAGCTGAGACTGGCCAGTGGGACGTCCCGGCCGACGCGCGTCCTAGCCC
GCGCGGGGCGCGATCGGTGATCACCTCGACGCGACGTGCATCGATGCGAAGATCTTCGAGTCGACTCGTACGATCGCGCC
ATGCCTTCGAAGGATGAATTCATACGGATCCGTGTACCAGTGCACTGCCGCGCGGAGCTCTAGGATCCGCGGGCCGTACC
ATGGCGCCCGGTGCAACGTCCTCGAGGTTAATCGAAATTGCATCGCACAAaAAACTaCagaCatGaTtaAtGaCCCcCGt
GatGaaGgCgAGctcTAGGtCGTtCTAtcctCtcCctAcGacACcgttAAGaTgAGaTTcAaaGCTgGtAGataGctTaa
aTCTGctaTTggctTcatTgCaTatTcCTgAGaCTcCagtAtTaGATattaGcctgccaatatTAtacttatctatctat
ctatctacatattacattacatattacatAtactattaatctactccctTaatggtagcttaaaattcatatctactccc
tTaatggtagcttaaaattcatatctactccctTaatggtagcttaaaattcatatctactccctTaatggtagcttaaa
attcatatctactccctTaatggtagcttaaaattcatatctactccctTaatggtagcttaaaattcatatctactccc
tTaatggtagcttaaaattcatatctaactaggcatctattgcctTaatggtagcttaaactttGTaattactaggcata
gatatctaccgtctattcctTaatggtagcattaaaattccgtaatatctatactagTAaTctatttcctTaatggtagc
ttaaaatttactagTAaTttatatctaCatcatcctcctatttagtgcctTaatggtagcttaaaattCatcatcctcta
tatctaGgctattgcctTaatggtagcaacttaaaattGgatatctaattcctTaatggtagcCttaaaattaatatgta
tttatatctaCctattCcctTaatggtagcttaaaattCatatctaaatgatAgacctTaatggtagcaacttaaaatta
atgaAgaAgaatatctatactagTAaTctatttagtgcctTaatggtagcttaaaatttactagTAaTttatatctaccg
tctattagtgcctTaatggtagcattaaaattccgtaatatctaTacttacattttcctTaatggtagcttaaaattTac
ttacattacattatatctattcacactctatttagtgcctTaatggtagcttaaaattttcacacttatatctagccCgc
tattgagtgcctTaatggtagcttaaactttTaattgccCggatatctactactaccacctattCagtgcctTaatggta
gccccccttaaaattctactaccacaAaCatatctaAtatctatttagtgcctTaatggtagcttaaaattAtatatatct
aTctattagtgcctTaatggtagcattaaaattTatatctactattcctTaatggtagcattaaaattatatctactatt
agtgcctTaatggtagcattaaaattatatctaTctattcctTaatggtagcattaaaattTatatctaCgctattgagt
gcctTaatggtagcttaaactttTaattCgatatctactattgagtgcctTaatggtagcttaaactttTaattatatct
aacaactattCagtgcctTaatggtagcttaaaattacaaatatctaacctattcgagtgcctTaatggtagcaacttaa
aattacatatctaActattCagtgcctTaatggtagcttaaaattAatatctaatActattAgaagtgcctTaatggtag
caacttaaaattatAatatctactattccagtgcctTaatggtagcaacttaaaattatatctaAtctatttagtgcctT
aatggtagcttaaaattAatatctaactctatttagtgcctTaatggtagcttaaaattacatatctatttcctTaatgg

tagcCttaaaatttatatctaActattcctTaatggtagcattaaaattAatatctaActattcctTaatggtagcatta
aaattAatatctaActattcctTaatggtagcattaaaattAatatctaActattcctTaatggtagcattaaaattAat
atctaActattcctTaatggtagcattaaaattAatatctaactatctattCagtgcctTaatggtagcccttaaaatta
ctatatattaataaCtatcaatctgTattTttcTCtaCtaCatCatatatTatcTatCtaCGaatCttCtt

File S6
pUAST-kkv-mos::NG protein

MSAMRHRPMAPPGQGPGAGTAGEHVDSDDNNFTDDESSPLTHDIYGGSQRTIQETKGWDVFRDPPIKIETGSTANQECLE
LTVKILKIFAYVITFIIVLTGGVIAKGTMLFMTSQVRKDKKMEYCNKDLGRDKSFVVRLPEEERVAWIWALLIAYALPEI
GALIRSARICFFKTFKVPKTGHFLFVWLMESLSAVGMALLMFVVLPQIDAIQGAMLTNCLCVVPGIFGLLSRTSKEGKRF
VKVIIDLAAVAAQVTGLVIWPLLENRRELWVIPVACVMISCGWWENYVSPQSPLGLVRALGRIKEEMKYTRYFCHIFLSI
WKILLFFTVTLLIYWAQGEEPGNLFAMYGDAFGPHKIIVYELPAGLGGVLPDTLESANIDTVDVDAAYNTVVYVLLLQIF
GAYLCYIFGKFACKILIQGFSYAFPVSLTVPLSVTFLIAACGIRIDDPCFFHDTIPDYLFFTSPSNFRFNNFVTEQMAWA
WILWLLSQTWIALHIWTPKCERLATTEKLFVQPMYSSLLIDQSMALNRRRDDQADVKTEDLSEIEKEKGDEYYETISVHT
DRSSAPNKPSIKSSDNITRIYSCATMWHETKDEMIEFLKSIMRMDEDQCARRVAQKYLRVLDPDYYEFETHIFFDDAFEI
SDHSDDDIQCNRFVKLLIATMDEAASEIHQTTIRLRPPKKYPTPYGGRLVWTLPGKTKFITHLKDKDRIRHRKRWSQVMY
MYYLLGHRLMELPISVDRKDVMAENTYLLTLDGDIDFNPSAVTLLVDLMKKNKNLGAACGRIHPIGSGPMVWYQKFEYAI
GHWLQKATEHMIGCVLCSPGCFSLFRGKGLMDDNVMRKYTTRSDEARHYVQYDQGEDRWLCTLLLQRGYRVEYSAASDAY
THCPEGFNEFYNQRRRWVPSTIANIMDLLADAKRTIKINDNISLLYIFYQMMLMGGTILGPGTIFLMLVGAFVAAFRIDN
WTSFHYNIVPILAFMFICFTCKSNIQLFVAQVLSTAYALIMMAVIVGTALQLGEDGIGSPSAIFLISMVGSFFIAACLHP
QEFWCITCGLIYLLSIPSMYLLLILYSIINLNVVSWGTREVVAKKTKKELEAEKKAAEEAKKRVKQKSMLSFLQSGIGDN
GDEEGSVEFSLAGLFRCIFCTHGKTSDEKQQLTSIAESLDTIKHRMDTIESAVDPHGHHASRHGRRRTTSSGSKDHHLLT
SVAEKSGDESDESDSDTSAEPKQERDFLTNPYWIEDPDVRKGEVDFLSSTEIQFWKDLIDQYLYPIDNDPVEQARIAKDL
KELRDSSVFAFFMINALFVLIVFLLQLNKDNIHVKWPFGVRTNITYDESTQEVHISKEYLQLEPIGLVFVFFFALILIIQ
FTAMLFHRFGTISHILASTELNFCKKKSEDLTQDQLIDKHAVEIVKNLQRLQGIDGDYDNDSGSGPDRIARRKTIQNLEK
ARQPRRQIGTLDVAFKKRFLKLTADAENNPATPILTRRLTMRAETIRALEVRKNSVMAERRKSAMQTLGAKNEYGITTGA
PINNNGALPNQRSGRVSNAGISIKDVFNVNGGGAEQIYGSNGGGTINQGYEHVIDEDGDGNSLRLTTRNPHPHPHHQVSW
GQNTNGGGGNGTGRLMVSKGEEDNMASLPATHELHIFGSINGVDFDMVGQGTGNPNDGYEELNLKSTKGDLQFSPWILVP
HIGYGFHQYLPYPDGMSPFQAAMVDGSGYQVHRTMQFEDGASLTVNYRYTYEGSHIKGEAQVKGTGFPADGPVMTNSLTA
ADWCRSKKTYPNDKTIISTFKWSYTTGNGKRYRSTARTTYTFAKPMAANYLKNQPMYVFRKTELKHSKTELNFKEWQKAF
TDVMGMDELYK

File S7
HDR-kkv3::smGFP

accaatgcttaatcagtgaggcacctatctcagcgatctgtctatttcgttcatccatagttgcctgactccccgtcgtg
tagataactacgatacgggagggcttaccatctggccccagcgctgcgatgataccgcgagaaccacgctcaccggctcc
ggatttatcagcaataaaccagccagccggaagggccgagcgcagaagtggtcctgcaactttatccgcctccatccagt
ctattaattgttgccgggaagctagagtaagtagttcgccagttaatagtttgcgcaacgttgttgccatcgctacaggc
atcgtggtgtcacgctcgtcgtttggtatggcttcattcagctccggttcccaacgatcaaggcgagttacatgatcccc
catgttgtgcaaaaaagcggttagctccttcggtcctccgatcgttgtcagaagtaagttggccgcagtgttatcactca
tggttatggcagcactgcataattctcttactgtcatgccatccgtaagatgcttttctgtgactggtgagtactcaacc
aagtcattctgagaatagtgtatgcggcgaccgagttgctcttgcccggcgtcaatacgggataataccgcgccacatag
cagaactttaaaagtgctcatcattggaaaacgttcttcggggcgaaaactctcaaggatcttaccgctgttgagatcca
gttcgatgtaacccactcgtgcacccaactgatcttcagcatcttttactttcaccagcgtttctgggtgagcaaaaaca
ggaaggcaaaatgccgcaaaaaagggaataagggcgacacggaaatgttgaatactcatattcttcctttttcaatatta
ttgaagcatttatcagggttattgtctcatgagcggatacatatttgaatgtatttagaaaaataaacaaataggggtca
gtgttacaaccaattaaccaattctgaacattatcgcgagcccatttatacctgaatatggctcataacaccccttgttt
gcctggcggcagtagcgcggtggtcccacctgaccccatgccgaactcagaagtgaaacgccgtagcgccgatggtagtg
tggggactccccatgcgagagtagggaactgccaggcatcaaataaaacgaaaggctcagtcgaaagactgggcctttcg
cccgggcgtattactcatccccataggtgcacatctccaaggagtaccttcagctggagcccatcggattggtgttcgt
attcttctttgctcttattttgataatccagttcacggccatgctgttccatcgatttggaaccatttcgcacatcctgg
cctcgactgaactcaacttctgcaagaagaagtctgaggatctgacacaggaccaactaatagataaggtaggtggcccg
taattgacattaatacaaataataaaattatgatattatttagcatgctgtggagatcgtcaagaacttgcagaggctgc
agggaatcgatggcgactacgacaatgactccggttccgggccagatcgcatagccaggcggaagaccattcaaaatctg
gagaaggcgcgacagccacgtcgccagatcggcaccctggatgtggcgttcaagaagcgcttcctgaaactcaccgccga
tgcggagaacaatccggccacgcccattctcacccgccgcctgacaatgcgggcggagaccatccgggcactggaggtgc
gtaagaattcggtgatggccgaaaggcggaagtccgcgatgcaaactctgggagcaaagaacgagtacggaatcacaact
ggagcaccggtgagattagaaaagtttttggagaaatttagatacaggatcataacttctttaatgcatatatgtagat
taacaacaatggagctctgccgaatcagcgaagcggacgtgtctcgaatgcaggcatcagcatcaaggatgtattcaatg
tgaacggtggtggtgccgaggtaaaggccaatttataatcataaaatatgtgtattctcatggaaatattttgttacgtc
cgcttacacttgcagcagatttacggctcgaatgggggcggcactatcaaccagggttacgagcatgtgatcgacgagga
cggcgacggcaactccctgcggctgaccaccaggaatccgcacccacacccgcaccaccaggtctcctggggccagaaca
ccaacggcggcggcggtaatggcacaggtcgcctggccggcatggtgagcaagggcgaggaggataacatggcctctctc
ccagcgacacatgagttacacatctttggctccatcaacggtgtggactttgacatggtgggtcagggcaccggcaatcc
aaatgatggttatgaggagttaaacctgaagtccaccaagggtgacctccagttctccccctggattctggtccctcata
tcgggtatggcttccatcagtacctgccctaccctgacgggatgtcgcctttccaggccgccatggtagatggctccgga
taccaagtccatcgcacaatgcagtttgaagatggtgcctcccttactgttaactaccgctacacctacgagggaagcca
catcaaaggagaggcccaggtgaaggggactggtttccctgctgacggtcctgtgatgaccaactcgctgaccgctgcgg
actggtgcaggtcgaagaagacttaccccaacgacaaaaccatcatcagtacctttaagtggagttacaccactggaaat
ggcaagcgctaccggagcactgcgcggaccacctacacctttgccaagccaatggcggctaactatctgaagaaccagcc
gatgtacgtgttccgtaagacggagctcaagcactccaagaccgagctcaacttcaaggagtggcaaaaggcctttaccg
atgtgatgggcatggacgagctgtacaagtaagtttaaactgaaggacccgcccatcctagacgtagagctaagttagac
ctaggtaacttgttaacgaattccgcctctttccatcaactgcctaaacttaagactaagccaatacctagaattgtagt
cagaaggaaggattgtgttgtatagtatgtatagggacgcgagtgagagccaggacactctggctcagaaggatcaggtc
gttcataaggcttcatggcaacagaacgaacattatgcttagctaatcacactgccagtgtcgatgtgtgccaataattg
ggtggctgatcgcccactgtggcagcaacattcatgcgatgtactatactatcatttgttaggttatctttaagtcaata
cccgcgtggatcatccgaaacacgcggccagtgcatgctgacacaaaatactcaacgtggagctgaaccgcagtggccta
tttgtacactcgacgacatatcgaggatagttctttttaaatgaataatcattgatgatttgatggattgtaaaactttt
taaaacaataaatagaagcatcaaacagatattcgatatttctcactcatcaagctacgcggccgcggacatatgcacac
ctgcgatcataacttcgtataatgtatgctatacgaagttatcgtacgggatctaattcaattagagactaattcaatta
gagctaattcaattaggatccaagcttatcgatttcgaaccctcgaccgccggagtataaatagaggcgcttcgtctacg
gagcgacaattcaattcaaacaagcaaagtgaacacgtcgctaagcgaaagctaagcaaataaacaagcgcagctgaaca
agctaaacaatcggctcgaagccggtcgccaccatggcctcctccgaggacgtcatcaaggagttcatgcgcttcaaggt
gcgcatggagggctccgtgaacggccacgagttcgagatcgagggcgagggcgagggccgcccctacgagggcacccaga
ccgccaagctgaaggtgaccaagggcggccccctgcccttcgcctgggacatcctgtcccccagttccagtacggctcc
aaggtgtacgtgaagcaccccgccgacatccccgactacaagaagctgtccttccccgagggcttcaagtgggagcgcgt
gatgaacttcgaggacggcggcgtggtgaccgtgacccaggactcctccctccaggacggctccttcatctacaaggtga
agttcatcggcgtgaacttcccctccgacggccccgtaatgcagaagaagactatgggctgggaggcgtccaccgagcgc
ctgtaccccgcgacggcgtgctgaagggcgagatccacaaggccctgaagctgaaggacggcggccactacctggtgga
gttcaagtccatctacatggccaagaagcccgtgcagctgcccggctactactacgtggactccaagctggacatcacct
cccacaacgaggactacaccatcgtggagcagtacgagcgcgccgagggccgccaccacctgttcctgtaggggccgcga
ctctagatcataatcagccataccacatttgtagaggttttacttgctttaaaaaacctcccacacctccccctgaacct
gaaacataaaatgaatgcaattgttgttgttaacttgtttattgcagcttataatggttacaaataaagcaatagcatca
caaatttcacaaataaagcatttttttcactgcattctagttgtggtttgtccaaactcatcaatgtatcttaaccggta

taacttcgtataatgtatgctatacgaagttatagaagagcactagtaaagatctcgatatttctcactctcatcaagca
tcggtcatcccaaggataagctacatccgaacccaccttcggtagtgggataagctacatccgaacccaccttcggtagt
ggagaccctaccccaagtttagaccaatacccgcatattttcggcttagagccctggtactatacgaataactttatttt
cgggaaagggtagaaacattactagacatcgcttgcgggacaacattgataaccagctatgatagataatagtattagac
acccatatggaaacgtcgtgtatgctcgcgttatcagcgagttaccttagtggaataaaaggttgcccttcttggcggct
atgggcaagcggtcgccctggaggaacgggtttatgaccgcgtcgtccagccgaggatcgggcatcacaggatatccggc
cttcttagccaggctgttaacgttgttcaggctgttctcctggcccatctgcatgatgtactgctccacggcggagttca
ggtcacccttgtcgccgcccacatcagcactatcctgttccgagcagtcgaacatatgctcgttgtcgcaggtacagtcc
tgagcggcctggaactcccggctgaagatggccgtgttttcgaagtcgacaatgcagctgccgccttgcgtctccatgtc
gtagccctcagggcagggattaggagggttgcagtaggcaggcaggccggcgtcgttcttcactatagtgggatacctgt
tgggaccctctcccattccgccggagatgaactggtgaccccacagcgaactgtgctgcagaaactcgtcatcccgcagc
gaaggatgggaggcagctcccgatgccgagcccgattgcaggatgcagtcgcttccgccgtcacacagcctcgcgtactc
cgaccgggagaccaggtccacgttgtccttggagccagcggctgcctcgttgcccacgtcgtaatagcccacctgcatgt
ctttgtccatgcgattcagcagatccgtcatcaagacatcggcaagaatatccttggcgctataggactgcacctggtag
ccactcaaggcgaccacgagcatcgccaccatcatgtacacacctggaaaaacatgggctattctcttacctcgaggctc
ttccgtcaatcgagttcaagggcgacacaaaatttattctaaatgcataataaatactgataacatcttatagtttgtat
tatattttgtattatcgttgacatgtataattttgatatcaaaaactgattttccctttattattttcgagatttatttt
cttaattctctttaacaaactagaaatattgtatatacaaaaaatcataaataatagatgaatagtttaattataggtgt
tcatcaatcgaaaaagcaacgtatcttatttaaagtgcgttgctttttttctcatttataaggttaaataattctcatata
tcaagcaaagtgacaggcgcccttaaatattctgacaaatgctctttccctaaactccccccataaaaaaacccgccgaa
gcgggtttttacgttatttgcggattaacgattactcgttatcagaaccgcccagggggcccgagcttaagactggccgt
cgttttacaacacagaaagagtttgtagaaacgcaaaaaggccatccgtcaggggccttctgcttagtttgatgcctggc
agttccctactctcgccttccgcttcctcgctcactgactcgctgcgctcggtcgttcggctgcggcgagcggtatcagc
tcactcaaaggcggtaatacggttatccacagaatcaggggataacgcaggaaagaacatgtgagcaaaaggccagcaaa
aggccaggaaccgtaaaaaggccgcgttgctggcgtttttccataggctccgcccccctgacgagcatcacaaaaatcga
cgctcaagtcagaggtggcgaaacccgacaggactataaagataccaggcgtttccccctggaagctccctcgtgcgctc
tcctgttccgaccctgccgcttaccggatacctgtccgcctttctcccttcgggaagcgtggcgctttctcatagctcac
gctgtaggtatctcagttcggtgtaggtcgttcgctccaagctgggctgtgtgcacgaaccccccgttcagcccgaccgc
tgcgccttatccggtaactatcgtcttgagtccaacccggtaagacacgacttatcgccactggcagcagccactggtaa
caggattagcagagcgaggtatgtaggcggtgctacagagttcttgaagtggtggcctaactacggctacactagaagaa
cagtatttggtatctgcgctctgctgaagccagttaccttcggaaaaagagttggtagctcttgatccggcaaacaaacc
accgctggtagcggtggtttttttgtttgcaagcagcagattacgcgcagaaaaaaaggatctcaagaagatcctttgat
cttttctacggggtctgacgctcagtggaacgacgcgcgcgtaactcacgttaagggattttggtcatgagcttgcgccg
tcccgtcaagtcagcgtaatgctctgctttt

File S8
HDR-kkv3-NG

ccaatgcttaatcagtgaggcacctatctcagcgatctgtctatttcgttcatccatagttgcctgactccccgtcgtg
tagataactacgatacgggagggcttaccatctggccccagcgctgcgatgataccgcgagaaccacgctcaccggctcc
ggatttatcagcaataaaccagccagccggaagggccgagcgcagaagtggtcctgcaactttatccgcctccatccagt
ctattaattgttgccgggaagctagagtaagtagttcgccagttaatagtttgcgcaacgttgttgccatcgctacaggc
atcgtggtgtcacgctcgtcgtttggtatggcttcattcagctccggttcccaacgatcaaggcgagttacatgatcccc
catgttgtgcaaaaaagcggttagctccttcggtcctccgatcgttgtcagaagtaagttggccgcagtgttatcactca
tggttatggcagcactgcataattctcttactgtcatgccatccgtaagatgcttttctgtgactggtgagtactcaacc
aagtcattctgagaatagtgtatgcggcgaccgagttgctcttgcccggcgtcaatacgggataataccgcgccacatag
cagaactttaaaagtgctcatcattggaaaacgttcttcggggcgaaaactctcaaggatcttaccgctgttgagatcca
gttcgatgtaacccactcgtgcacccaactgatcttcagcatcttttactttcaccagcgtttctgggtgagcaaaaaca
ggaaggcaaaatgccgcaaaaaagggaataagggcgacacggaaatgttgaatactcatattcttcctttttcaatatta
ttgaagcatttatcagggttattgtctcatgagcggatacatatttgaatgtatttagaaaaataaacaaataggggtca
gtgttacaaccaattaaccaattctgaacattatcgcgagcccatttatacctgaatatggctcataacaccccttgttt
gcctggcggcagtagcgcggtggtcccacctgaccccatgccgaactcagaagtgaaacgccgtagcgccgatggtagtg
tggggactccccatgcgagagtagggaactgccaggcatcaaataaaacgaaaggctcagtcgaaagactgggcctttcg
cccgggcgtattactcatcccccataggtgcacatctccaaggagtaccttcagctggagcccatcggattggtgttcgt
attcttctttgctcttattttgataatccagttcacggccatgctgttccatcgatttggaaccatttcgcacatcctgg
cctcgactgaactcaacttctgcaagaagaagtctgaggatctgacacaggaccaactaatagataaggtaggtggcccg
taattgacattaatacaaataataaaattatgatattatttagcatgctgtggagatcgtcaagaacttgcagaggctgc
agggaatcgatggcgactacgacaatgactccggttccgggccagatcgcatagccaggcggaagaccattcaaaatctg
gagaaggcgcgacagccacgtcgccagatcggcaccctggatgtggcgttcaagaagcgcttcctgaaactcaccgccga
tgcggagaacaatccggccacgcccattctcacccgccgcctgacaatgcgggcggagaccatccgggcactggaggtgc
gtaagaattcggtgatggccgaaaggcggaagtccgcgatgcaaactctgggagcaaagaacgagtacggaatcacaact
ggagcaccggtgagattagaaaagtttttggagaaattttagatacaggatcataacttctttaatgcatatatgtagat
taacaacaatggagctctgccgaatcagcgaagcggacgtgtctcgaatgcaggcatcagcatcaaggatgtattcaatg
tgaacggtggtggtgccgaggtaaaggccaatttataatcataaaatatgtgtattctcatggaaatattttgttacgtc
cgcttacacttgcagcagatttacggctcgaatgggggcggcactatcaaccagggttacgagcatgtgatcgacgagga
cggcgacggcaactccctgcggctgaccaccaggaatccgctacgccacccgcaccaccaggtctcctggggccagaaca
ccaacggcggcggcggtaatggcacaggtcgcctggccggcatgtacccttatgatgtgcccgattatgctggctaccct
tatgatgttcctgattacgccggatatccgtatgacgtgccagactacgcgggaggtgtgagcaagggagaggagttgtt
taccggcgtggtccccatactggtggagttggatggcgacgttaatggacataaattctcggtccgcggcgagggagagg
gagacgccaccaatggcaagctgacgcttaagttcatttgcacgactggaaaattgcctgtcccctggccgaccctggtc
accacactgggcggaggagtgcagtgcttctcccgttacccagaccacatgaagcagcacgatttcttcaagagcgcaat
gccggagggatacgtgcaagaacggactatttccttcaaggatgacggaacctataagacccgtgcggaggtcaaattcg
agggtgacaccctggtgaaccgaattgaactcaaaggaatcgatttcaaggaggatggaaatattctgggtcacaagctg
gaatacaacttcaacagccataatgtgtacattacggctgataagcagaagaacggcattaaggccaatttcaagatccg
ccacaacgttgagggtggatatccctacgacgtgcccgattatgcgggcggatatccgtatgatgttccagattatgctg
gtggtggaggctatccctatgatgtccccgactacgccggaggttacccatacgatgtgcccgactacgctggaggcgat
ggcagcgtgcagctggcagatcattatcaacagaatacccccataggtgatggccccgttctgcttccagataatcacta
cctttccacccagagcgtgctttcgaaagacccgaacgaaaagcgtgatcacatggtcctgctggagtttgtgaccgcgg
ctggaatcaccctgggtatggacgaactctacaagggtggctacccctacgatgtgccggattacgctggatatccgtat
gacgtaccggactatgccggttacccgtacgatgtcccggactacgcttaagtttaaactgaaggacccgcccatcctag
acgtagagctaagttagacctaggtaacttgttaacgaattccgcctctttccatcaactgcctaaacttaagactaagc
caatacctagaattgtagtcagaaggaaggattgtgttgtatagtatgtatagggacgcgagtgagagccaggacactct
ggctcagaaggatcaggtcgttcataaggcttcatggcaacagaacgaacattatgcttagctaatcacactgccagtgt
cgatgtgtgccaataattgggtggctgatcgcccactgtggcagcaacattcatgcgatgtactatactatcatttgtta
ggttatctttaagtcaatacccgcgtggatcatccgaaacacgcggccagtgcatgctgacacaaaatactcaacgtgga
gctgaaccgcagtggcctatttgtacactcgacgacatatcgaggatagttctttttaaatgaataatcattgatgattt
gatggattgtaaaactttttaaaacaataaatagaagcatcaaacagatattcgatatttctcactcatcaagctacgcg
gccgcggacatatgcacacctgcgatcataacttcgtataatgtatgctatacgaagttatcgtacgggatctaattcaa
ttagagactaattcaattagagctaattcaattaggatccaagcttatcgatttcgaaccctcgaccgccggagtataaa
tagaggcgcttcgtctacggagcgacaattcaattcaaacaagcaaagtgaacacgtcgctaagcgaaagctaagcaaat
aaacaagcgcagctgaacaagctaaacaatcggctcgaagccggtcgccaccatggcctcctccgaggacgtcatcaagg
agttcatgcgcttcaaggtgcgcatggagggctccgtgaacggccacgagttcgagatcgagggcgagggcgagggccgc
ccctacgagggcacccagaccgccaagctgaaggtgaccaagggcggccccctgcccttcgcctgggacatcctgtcccc
ccagttccagtacggctccaaggtgtacgtgaagcaccccgccgacatccccgactacaagaagctgtccttccccgagg
gcttcaagtgggagcgcgtgatgaacttcgaggacggcggcgtggtgaccgtgacccaggactcctccctccaggacggc
tccttcatctacaaggtgaagttcatcggcgtgaacttcccctccgacggccccgtaatgcagaagaagactatgggctg
ggaggcgtccaccgagcgcctgtaccccgcgacggcgtgctgaagggcgagatccacaaggccctgaagctgaaggacg
gcggccactacctggtggagttcaagtccatctacatggccaagaagcccgtgcagctgcccggctactactacgtggac

tccaagctggacatcacctcccacaacgaggactacaccatcgtggagcagtacgagcgcgccgagggccgccaccacct
gttcctgtaggggccgcgactctagatcataatcagccataccacatttgtagaggttttacttgctttaaaaaacctcc
cacacctccccctgaacctgaaacataaaatgaatgcaattgttgttgttaacttgtttattgcagcttataatggttac
aaataaagcaatagcatcacaaatttcacaaataaagcatttttttcactgcattctagttgtggtttgtccaaactcat
caatgtatcttaaccggtataacttcgtataatgtatgctatacgaagttatagaagagcactagtaaagatctcgatat
ttctcactctcatcaagcatcggtcatcccaaggataagctacatccgaacccaccttcggtagtgggataagctacatc
cgaacccaccttcggtagtggagaccctaccccaagtttagaccaatacccgcatattttcggcttagagccctggtact
atacgaataactttattttcgggaaagggtagaaacattactagacatcgcttgcgggacaacattgataaccagctatg
atagataatagtattagacacccatatggaaacgtcgtgtatgctcgcgttatcagcgagttaccttagtggaataaaag
gttgcccttcttggcggctatgggcaagcggtcgccctggaggaacgggtttatgaccgcgtcgtccagccgaggatcgg
gcatcacaggatatccggccttcttagccaggctgttaacgttgttcaggctgttctcctggcccatctgcatgatgtac
tgctccacggcggagttcaggtcacccttgtcgccgcccacatcagcactatcctgttccgagcagtcgaacatatgctc
gttgtcgcaggtacagtcctgagcggcctggaactcccggctgaagatggccgtgttttcgaagtcgacaatgcagctgc
cgccttgcgtctccatgtcgtagccctcagggcagggattaggagggttgcagtaggcaggcaggccggcgtcgttcttc
actatagtgggatacctgttgggaccctctcccattccgccggagatgaactggtgaccccacagcgaactgtgctgcag
aaactcgtcatcccgcagcgaaggatgggaggcagctcccgatgccgagcccgattgcaggatgcagtcgcttccgccgt
cacacagcctcgcgtactccgaccgggagaccaggtccacgttgtccttggagccagcggctgcctcgttgcccacgtcg
taatagcccacctgcatgtctttgtccatgcgattcagcagatccgtcatcaagacatcggcaagaatatccttggcgct
ataggactgcacctggtagccactcaaggcgaccacgagcatcgccaccatcatgtacacacctggaaaaacatgggcta
ttctcttacctcgaggctcttccgtcaatcgagttcaagggcgacacaaaatttattctaaatgcataataaatactgat
aacatcttatagtttgtattatattttgtattatcgttgacatgtataattttgatatcaaaaactgattttccctttat
tattttcgagatttattttcttaattctctttaacaaactagaaatattgtatatacaaaaaatcataaataatagatga
atagtttaattataggtgttcatcaatcgaaaaagcaacgtatcttatttaaagtgcgttgctttttctcatttataag
gttaaataattctcatatatcaagcaaagtgacaggcgcccttaaatattctgacaaatgctctttccctaaactccccc
cataaaaaaacccgccgaagcgggtttttacgttatttgcggattaacgattactcgttatcagaaccgcccagggggcc
cgagcttaagactggccgtcgttttacaacacagaaagagtttgtagaaacgcaaaaaggccatccgtcaggggccttct
gcttagtttgatgcctggcagttccctactctcgccttccgcttcctcgctcactgactcgctgcgctcggtcgttcggc
tgcggcgagcggtatcagctcactcaaaggcggtaatacggttatccacagaatcaggggataacgcaggaaagaacatg
tgagcaaaaggccagcaaaaggccaggaaccgtaaaaaggccgcgttgctggcgtttttccataggctccgcccccctga
cgagcatcacaaaaatcgacgctcaagtcagaggtggcgaaacccgacaggactataaagataccaggcgtttccccctg
gaagctccctcgtgcgctctcctgttccgaccctgccgcttaccggatacctgtccgcctttctcccttcgggaagcgtg
gcgctttctcatagctcacgctgtaggtatctcagttcggtgtaggtcgttcgctccaagctgggctgtgtgcacgaacc
ccccgttcagcccgaccgctgcgccttatccggtaactatcgtcttgagtccaacccggtaagacacgacttatcgccac
tggcagcagccactggtaacaggattagcagagcgaggtatgtaggcggtgctacagagttcttgaagtggtggcctaac
tacggctacactagaagaacagtatttggtatctgcgctctgctgaagccagttaccttcggaaaaagagttggtagctc
ttgatccggcaaacaaaccaccgctggtagcggtggtttttttgtttgcaagcagcagattacgcgcagaaaaaaaggat
ctcaagaagatcctttgatcttttctacggggtctgacgctcagtggaacgacgcgcgcgtaactcacgttaagggattt
tggtcatgagcttgcgccgtcccgtcaagtcagcgtaatgctctgctttt

File S9
> [Kkv::smFP]

MSAMRHRPMAPPGQGPGAGTAGEHVDSDDNNFTDDESSPLTHDIYGGSQRTIQETKGWDVFRDPPIKIETGSTANQECLE
LTVKILKIFAYVITFIIVLTGGVIAKGTMLFMTSQVRKDKKMEYCNKDLGRDKSFVVRLPEEERVAWIWALLIAYALPEI
GALIRSARICFFKTFKVPKTGHFLFVWLMESLSAVGMALLMFVVLPQIDAIQGAMLTNCLCVVPGIFGLLSRTSKEGKRF
VKVIIDLAAVAAQVTGLVIWPLLENRRELWVIPVACVMISCGWWENYVSPQSPLGLVRALGRIKEEMKYTRYFCHIFLSI
WKILLFFTVTLLIYWAQGEEPGNLFAMYGDAFGPHKIIVYELPAGLGGVLPDTLESANIDTVDVDAAYNTVVYVLLLQIF
GAYLCYIFGKFACKILIQGFSYAFPVSLTVPLSVTFLIAACGIRIDDPCFFHDTIPDYLFFTSPSNFRFNNFVTEQMAWA
WILWLLSQTWIALHIWTPKCERLATTEKLFVQPMYSSLLIDQSMALNRRRDDQADVKTEDLSEIEKEKGDEYYETISVHT
DRSSAPNKPSIKSSDNITRIYSCATMWHETKDEMIEFLKSIMRMDEDQCARRVAQKYLRVLDPDYYEFETHIFFDDAFEI
SDHSDDDIQCNRFVKLLIATMDEAASEIHQTTIRLRPPKKYPTPYGGRLVWTLPGKTKFITHLKDKDRIRHRKRWSQVMY
MYYLLGHRLMELPISVDRKDAIAENTYLLTLDGDIDFKPNAVTLLVDLMKKNKNLGAACGRIHPVGSGPMVWYQLFEYAI
GHWLQKATEHMIGCVLCSPGCFSLFRGKALMDDNVMKKYTTRSDEARHYVQYDQGEDRWLCTLLLQRGYRVEYSAASDAY
THCPEGFNEFYNQRRRWVPSTIANIMDLLADAKRTIKINDNISLLYIFYQMMLMGGTILGPGTIFLMLVGAFVAAFRIDN
WTSFHYNIVPILAFMFICFTCKSNIQLFVAQVLSTAYALIMMAVIVGTALQLGEDGIGSPSAIFLISMVGSFFIAACLHP
QEFWCITCGLIYLLSIPSMYLLLILYSIINLNVVSWGTREVVAKKTKKELEAEKKAAEEAKKRVKQKSMLSFLQSGIGDN
GDEEGSVEFSLAGLFRCIFCTHGKTSDEKQQLTSIAESLDTIKHRMDTIESAVDPHGHHASRHGRRRTTSSGSKDHHLLT
SVAEKSGDESDESDSDTSAEPKQERDFLTNPYWIEDPDVRKGEVDFLSSTEIQFWKDLIDQYLYPIDNDPVEQARIAKDL
KELRDSSVFAFFMINALFVLIVFLLQLNKDNIHVKWPFGVRTNITYDESTQEVHISKEYLQLEPIGLVFVFFFALILIIQ
FTAMLFHRFGTISHILASTELNFCKKKSEDLTQDQLIDKHAVEIVKNLQRLQGIDGDYDNDSGSGPDRIARRKTIQNLEK
ARQPRRQIGTLDVAFKKRFLKLTADAENNPATPILTRRLTMRAETIRALEVRKNSVMAERRKSAMQTLGAKNEYGITTGA
PINNNGALPNQRSGRVSNAGISIKDVFNVNGGGAEQIYGSNGGGTINQGYEHVIDEDGDGNSLRLTTRNPHPHPHHQVSW
GQNTNGGGGNGTGRLGMYPYDVPDYAGYPYDVPDYAGYPYDVPDYAGGVSKGEELFTGVVPILVELDGDVNGHKFSVRGE
GEGDATNGKLTLKFICTTGKLPVPWPTLVTTLGGGVQCFSRYPDHMKQHDFFKSAMPEGYVQERTISFKDDGTYKTRAEV
KFEGDTLVNRIELKGIDFKEDGNILGHKLEYNFNSHNVYITADKQKNGIKANFKIRHNVEGGYPYDVPDYAGGYPYDVPD
YAGGGGYPYDVPDYAGGYPYDVPDYAGGDGSVQLADHYQQNTPIGDGPVLLPDNHYLSTQSVLSKDPNEKRDHMVLLEFV
TAAGITLGMDELYKGGYPYDVPDYAGYPYDVPDYAGYPYDVPDYA

File S10
> [Kkv::NG] protein from edited gene

MSAMRHRPMAPPGQGPGAGTAGEHVDSDDNNFTDDES**S**PLTHDIYGGSQRTIQETKGWDVFRDPPIKIETGSTANQECLE
LTVKILKIFAYVITFIIVLTGGVIAKGTMLFMTSQVRKDKKMEYCNKDLGRDKSFVVRLPEEERVAWIWALLIAYALPEI
GALIRSARICFFKTFKVPKTGHFLFVWLMESLSAVGMALLMFVVLPQIDAIQGAMLTNCLCVVPGIFGLLSRTSKEGKRF
VKVIIDLAAVAAQVTGLVIWPLLENRRELWVIPVACVMISCGWWENYVSPQSPLGLVRALGRIKEEMKYTRYFCHIFLSI
WKILLFFTVTLLIYWAQGEEPGNLFAMYGDAFGPHKIIVYELPAGLGGVLPDTLESANIDTVDVDAAYNTVVYVLLLQIF
GAYLCYIFGKFACKILIQGFSYAFPVSLTVPLSVTFLIAACGIRIDDPCFFHDTIPDYLFFTSPSNFRFNNFVTEQMAWA
WILWLLSQTWIALHIWTPKCERLATTEKLFVQPMYSSLLIDQSMALNRRRDDQADVKTEDLSEIEKEKGDEYYETISVHT
DRSSAPNKPSIKSSDNITRIYSCATMWHETKDEMIEFLKSIMRMDEDQCARRVAQKYLRVLDPDYYEFETHIFFDDAFEI
SDHSDDDIQCNRFVKLLIATMDEAASEIHQTTIRLRPPKKYPTPYGGRLVWTLPGKTKFITHLKDKDRIRHRKRWSQVMY
MYYLLGHRLMELPISVDRKDAIAENTYLLTLDGDIDFKPNAVTLLVDLMKKNKNLGAACGRIHPVGSGPMVWYQLFEYAI
GHWLQKATEHMIGCVLCSPGCFSLFRGKALMDDNVMKKYTTRSDEARHYVQYDQGEDRWLCTLLLQRGYRVEYSAASDAY
THCPEGFNEFYNQRRRWVPSTIANIMDLLADAKRTIKINDNISLLYIFYQMMLMGGTILGPGTIFLMLVGAFVAAFRIDN
WTSFHYNIVPILAFMFICFTCKSNIQLFVAQVLSTAYALIMMAVIVGTALQLGEDGIGSPSAIFLISMVGSFFIAACLHP
QEFWCITCGLIYLLSIPSMYLLLILYSIINLNVVSWGTREVVAKKTKKELEAEKKAAEEAKKRVKQKSMLSFLQSGIGDN
GDEEGSVEFSLAGLFRCIFCTHGKTSDEKQQLTSIAESLDTIKHRMDTIESAVDPHGHHASRHGRRRTTSSGSKDHHLLT
SVAEKSGDESDESDSDTSAEPKQERDFLTNPYWIEDPDVRKGEVDFLSSTEIQFWKDLIDQYLYPIDNDPVEQARIAKDL
KELRDSSVFAFFMINALFVLIVFLLQLNKDNIHVKWPFGVRTNITYDESTQEVHISKEYLQLEPIGLVFVFFFALILIIQ
FTAMLFHRFGTISHILASTELNFCKKKSEDLTQDQLIDKHAVEIVKNLQRLQGIDGDYDNDSGSGPDRIARRKTIQNLEK
ARQPRRQIGTLDVAFKKRFLKLTADAENNPATPILTRRLTMRAETIRALEVRKNSVMAERRKSAMQTLGAKNEYGITTGA
PINNNGALPNQRSGRVSNAGISIKDVFNVNGGGAEQIYGSNGGGTINQGYEHVIDEDGDGNSLRLTTRNPHPHPHHQVSW
GQNTNGGGGNGTGRLAGMVSKGEEDNMASLPATHELHIFGSINGVDFDMVGQGTGNPNDGYEELNLKSTKGDLQFSPWILVP
HIGYGFHQYLPYPDGMSPFQAAMVDGSGYQVHRTMQFEDGASLTVNYRYTYEGSHIKGEAQVKGTGFPADGPVMTNSLTA
ADWCRSKKTYPNDKTIISTFKWSYTTGNGKRYRSTARTTYTFAKPMAANYLKNQPMYVFRKTELKHSKTELNFKEWQKAF
DVMGMDELYK

File S11

Pairs of oligonucleotides used for construction of gRNA expressing plasmids D01 and DH1.

DO1: GTCGTTTCTCACTCTCATCAAACT 3′ to kkv in intergenic region
AAAGAGTGAGAGTAGTTTGACAAA

DH1: GTCGCTGGTGGTGCGGGTGGGGAT 5′ part of last exon
GACCACCACGCCCACCCCTACAAA
